# Whole genome phylogeny of Cyanobacteria documents a distinct evolutionary trajectory of marine picocyanobacteria

**DOI:** 10.1101/2021.05.26.445609

**Authors:** Otakar Strunecký, Markéta Wachtlová, Michal Koblížek

## Abstract

During their 2.7 Gyr long evolution cyanobacteria radiated into a large number of different lineages. To better understand the evolution of cyanobacteria we compared a whole genome phylogenetic tree using 1,047 concatenated single copy orthologues of *Prochlorococcus marinus* MIT9313 obtained from 93 reference prokaryotic species with traditional phylogenies inferred from 16S rRNA gene and 109 genes found in all used genomes. In contrast to the traditional phylogeny, our genome wide analysis shows a split between picocyanobacteria encompassing all marine *Prochlorococcus* species, marine “*Synechococcus*” species, and members of freshwater *Cyanobium* genus (altogether referred here as the “PSC” clade) and the remaining groups of extant cyanobacteria. To determine the influence of the horizontal gene transfer on the overall tree topology we removed the 374 genes identified as potentially transferred genes. A newly calculated tree utilizing the remaining 673 genes displayed the same topology as the former whole genome tree with the PSC clade as a basal group of all cyanobacteria. The picocyanobacteria also exhibited distinctly larger similarity to reference bacteria’s cell and genome size, carboxyzome architecture, various metabolic pathways, and chlorophyll synthesis than to the other cyanobacteria. Potentially horizontally transferred genes were found in connected chains throughout fundamental metabolic pathways suggesting that evolution of these genes was severely limited and or brought to a standstill. The environment related genes connected with metabolism of nitrogen, sulfur, and scarce seawater metals were more conserved in PSC group as they were already best tuned for its poor and stable environment. Other genes were found predominantly sequentially static as they were already accurately adapted with rare beneficial mutations. The PSC clade thus represents an isolated cyanobacterial lineage which followed a distinct evolutionary trajectory driven by its specific environment.

## Introduction

Cyanobacteria represent one of the most ecologically important groups of microorganisms in the entire history of Earth. Their origin dates back to around 2.7 Gyr ago during the Archean eon (Tomitani et al. 2006). Oxygenic photosynthesis that first appeared in their last common ancestor is regarded among the most significant innovations in the evolution of life, changing entirely the chemistry of the planet via the Great Oxygenation Event (Lyons et al. 2014). The capacity to harvest light through photosynthesis provides an inexhaustible source of energy which fuels all but few of the ecosystems on the planet. In particular marine picocyanobacteria *Prochlorococcus* and *Synechococcus* are responsible for at least one quarter of the global primary production (Garcia-Pichel et al. 2003; Rocap et al. 2002; Whitton 2012; Whitton and Potts 2000). Cyanobacteria have colonized all of the natural habitats within the photic zone - oceans, lakes, soils, hot and cold deserts, polar ice, and thermal springs (Brock 1978; Hess et al. 2016; Komárek 1999; Michaud et al. 2012; Paerl 2012) during the course of their long evolutional history. This dispersion into a wide range of habitats resulted in multiple lineages with a complex evolution.

Cyanobacteria were, together with algae, originally classified as primitive plants (Haeckel 1866) through the application of the botanical classification based on their photosynthetic pigments, cellular size and morphology. Morphologic determination has remained the only method of classification of cyanobacteria for a long time. The situation changed dramatically during the second half of the 20^th^ century with the introduction of modern laboratory techniques such as electron microscopy (i.e. Rippka et al. 1974), flow cytometry (i.e. Chisholm et al. 1988) and a variety of molecular methods (i.e. Nübel et al. 1997). In particular, the use of 16S rRNA sequences and the adoption of mathematical algorithms to infer relationships of individual species offered a straightforward method to analyze prokaryotic phylogeny. The main result of this “molecular revolution” was an abandonment of the unfitting botanical classification and the clear establishment of Cyanobacteria as an independent phylum within the domain Bacteria (Woese 1987). Despite the original wide acceptance of 16S rRNA derived phylogenies, it was soon recognized that they do not represent an “ultimate” solution as any phylogeny based on a single gene provides ground only for a partial reconstruction of the true phylogeny. This conundrum was solved using concatenated protein marker gene trees derived from isolated and population genomes which are much less susceptible to chimeric artifacts (Parks et al. 2015). The 16S rRNA database is currently two orders of magnitude larger than the genome database and must be used in congruence with genome-based taxonomy to provide taxonomic continuity in the literature (Hugenholtz et al. 2016).

In general inferring prokaryotic phylogeny represents an extremely challenging task. The original concept of the tree of life was developed for multicellular organisms (Darwin 1859). By definition it assumes only vertical evolution through a branching lineage of ancestors and descendants. In the case of prokaryotic organisms with a much more flexible genetic inventory and frequent horizontal gene transfer events (HGT) the whole concept of the “natural” phylogenetic tree is questionable. Despite their inherent problems phylogenetic trees are widely used as easy-to-understand tools illustrating the relationships among individual organisms.

A frequently applied approach is to select several genes with presumably vertical evolution and infer the overall phylogeny only on the selected subset (Hug et al. 2016; Komárek et al. 2014). Such an approach leads to significant bias since it may omit a large number of genes or traits which may be critical for the fitness and evolutionary success of individual organisms. Robust phylogeny using a multiple gene approach has been debated for a long time (Bhandari et al. 2012; Swingley et al. 2008). Shall we restrict our analyses to the phylogeny of several conservative genes or shall we use as many genes as possible, including the genes transferred horizontally or genes with rapid evolution? The selection of phylogenetic markers is frequently subjective, considering the housekeeping genes as being better for a “true” phylogenetical relationship assuming that they are passed on by vertical inheritance only (i.e. Cody et al. 2014; Kurmayer et al. 2015). Prokaryotic phylogeny based on a small set of conservative genes may reduce or even contradict the phylogenetic signal of the majority of genes present in the organism. All these genes are important for the evolutionary success of the organism. Reducing the phylogenetic analysis to a small set of standard phylogenetic markers may result only into a coarse approximation to the true phylogenetic relationships.

Another standard approach is the use of an outgroup that roots the tree and sets the distance from the last common ancestor of the studied groups. The majority of the studies presenting 16S rRNA phylogeny of cyanobacteria do use an outgroup. Contrary to this practice, the multi gene phylogenies inferred for cyanobacteria using 27 to 135 genes for 32 to 131 cyanobacteria (Blank and Sanchez-Baracaldo 2010; Dagan et al. 2013; Larsson et al. 2011; Sanchez-Baracaldo et al. 2005) do not present any outgroup. They most frequently rely on cyanobacterium *Gloeobacter* as an outgroup which is usually considered to be the most primitive cyanobacterium (e.g. Mareš et al. 2013; Seo and Yokota 2003). A recent study using 756 orthologues of a meticulously selected group of 65 cyanobacteria (Schirrmeister et al. 2015) also shows the standard cyanobacterial phylogeny with *Gloeobacter* at its base using maximum likelihood inference without outgroup composed of reference bacteria.

To advance the understanding and insight into the cyanobacterial phylogeny we decided to base our analysis on the complete genetic inventory of the studied species without *a priori* rejecting any gene. The highest number of common genes through the domain bacteria were concatenated with an assumption of minimizing the influence of HGTs distorting phylogenic relations. In this study we present a phylogenetic tree of Cyanobacteria based on a set of over 1000 genes shared by the majority of analyzed species and compared them with selected representatives of other bacterial phyla forming a significant outgroup of 45 reference bacteria that has never been used in similar studies in the past.

## Material and Methods

A data set comprehensively covering the complete genomes of cyanobacteria and the reference bacteria was generated using publicly available genomes from NCBI (ftp://ftp.ncbi.nih.gov/genomes/Bacteria/) and from the Joint Genome Institute’s IMG-M database (img.jgi.doe.gov). The number of analyzed genomes was optimized to represent a wide range of phylogenetic groups that would not distort the computed phylogenies due to computational artefacts and a reasonable computer time needed for phylogenetic analysis of very long alignments. We included 48 genomes of cyanobacteria that represented complete and well-annotated cyanobacterial genomes. At least one genome from every clade of cyanobacteria was chosen to equally represent individual lineages of cyanobacteria. This independent selection very well agreed with another *ad hoc* selection of genomes made by Schirrmeister et al. (2016) who calculated molecular clocks of cyanobacteria using multigene comparison. We then included 27 bacterial genomes of neighboring phyla to the PSC clade which were selected from organisms identified as potential donors of horizontally transferred genes. Third group contained 18 diverse non-cyanobacterial genomes from the remaining bacterial phyla. We were unable to use the Melainabacteria in our dataset due to its highly fragmented genome scaffold sequence in the GenBank.

A phylogenetic analysis of 16S rRNA genes of the 93 selected bacteria and cyanobacteria (Supplementary Table S2) was conducted. 1,460 nucleotides from 16S rRNA gene (from a position 271475 to 272935 *Escherichia coli* str. K-12 CP009685) were aligned using MAFFT --localpair --maxiterate 1000 and secondary structure of RNA option-Q-INS-I included. The topology of the phylogenetic tree was based on Bayesian analysis (MB) computed in MrBayes 3.2.2 (Ronquist and Huelsenbeck 2003). In the Bayesian analysis, four runs of four Markov chains were calculated for 12 million generations, sampling every 1000 generations. The initial 25% generations were discarded as burn-in. The detailed settings were as follows: Rates= equal, Nucmodel= 4by4, statefreqpr = dirichlet(1.0), mcmc ngen=12000000 nruns=4 nchains=4 temp=0.200 swapfreq=1 nswaps=1 samplefreq=1000 mcmcdiagn=Yes minpartfreq=0.1 Stopval=0.01, burninfrac=0.25. The tree was validated by maximum likelihood (ML) method in RAxML 7.0.4 (Stamatakis 2006) under GTR +G+I model with 1,000 bootstrap repetitions, and neighbor joining (NJ) under maximum composite likelihood method +G in MEGA 6.06 (Tamura et al. 2011) with 1,000 bootstrap repetitions using all sites.

The whole genomic phylogenetic tree was inferred using a custom BLAST database from the 93 genomes and BLAST+ 2.2.28 (Camacho et al. 2009) with an E-value of −10. *Prochlorococcus marinus*, strain MIT9313 (NC_005071.1) was selected as a reference organism due to its relatively small and well-annotated genome and intermediate phylogenetic position between *Prochlorococcus* and marine *Synechococcus* strains. Amino acid sequences of its genes were blasted against the constructed BLAST database. The obtained 2269 sets of amino acid sequences were aligned and reordered using MAFFT with --localpair --maxiterate 1000 settings (Katoh and Standley 2013). Sequences shorter than 80% of the reference sequence and duplicate sequences were deleted from further processing using a custom PERL script. The PERL scripts are available at /sourceforge.net/projects/pscgroup/. Gene alignments missing more than one third of the organisms were discarded. Including amino acid positions with 25% to 40% gaps in alignments is common practice (Criscuolo and Gribaldo 2011; Schirrmeister et al. 2015) and commonly increases the tree accuracy (Jiang et al. 2014; Wiens 2006). The remaining 1,047 alignments were concatenated using a PERL script providing the final 406,617 amino acid alignment. The validity of 1,047 used gene alignments were checked by OD-seq (Jehl et al. 2015) and Guidance v.2.02 (Sela et al. 2015), however no sequence outliers were detected by this approach. The phylogenetic analysis was performed using the NJ algorithm with the JTT model in MEGA 6.06 (Tamura et al. 2011) including pairwise deletion of missing amino acid sites with 500 bootstrap repetitions. The ML phylogeny was inferred using the RAxML algorithm (Stamatakis 2006) with proteins under 25 categories with GTR PROTCAT protein substitution model with 500 bootstrap repetitions (due to exceptionally time demanding comparison of very long alignments) using the CIPRES Science Gateway (Miller et al. 2010). The phylogenic analysis of 109 proteins that were present in all 93 genomes was done for 61,874 amino acids in the same way as for the whole genomic tree. The phylogenic analysis of no-HGT inferred trees was made for 673 protein alignments with a combined length of 301,420 amino acids in the same way as for the whole genomic tree.

To analyze the phylogeny of photosynthesis in cyanobacteria we selected 10 genes encoding various subunits of photosystem I and 7 genes encoding the subunits of photosystem II (Supplementary Table S3). Phylogeny of PSI and PSII with 5,656 amino acid long alignment was calculated by MrBayes 3.2.2 with four runs of four Markov chains for 5 million generations, sampling every 1000 generations (Rates= equal, Codon=Universal, Coding=All, statefreqpr = dirichlet, mcmc ngen=5000000 nruns=4 nchains=4 temp=0.200 swapfreq=1 nswaps=1 samplefreq=1000 mcmcdiagn=Yes minpartfreq=0.1 allchains=No relburnin=Yes burnin=0, Stopval=0.01, burninfrac=0.25). The initial 25% generations were discarded as burn-in. The NJ and ML trees were inferred using the JTT model in MEGA 6.06 with 1,000 bootstrap repetitions and the RAxML algorithm with proteins under 25 categories with GTR PROTCAT protein substitution model with 1,000 bootstraps, respectively.

Horizontally transferred genes were identified as follows: 2,269 designated open reading frames from the genome of *Prochlorococcus marinus*, strain MIT9313 (NC_005071.1) were blasted against non-redundant GenBank protein database. Blastp was performed by BLAST+ 2.2.28 (Altschul et al. 1990) limited by E-value of −10. Blasted sequences were aligned according to the query. Redundant sequences from any particular organism and sequences shorter than 80 percent of query sequence were deleted from further processing using a PERL script. Aligned genes were used for phylogenetic reconstruction in FastTreeMP (Price et al. 2010) with -lg -gamma settings. GI of all sequences were used to download the taxonomy information from GenBank for all organisms in phylogenetic trees. The phylogenetically closest organisms to PSC clade were recorded for every tree using a custom PERL script. If the node of PSC clade was inside of clades formed by reference bacteria and not adjacent to other cyanobacteria then the genes were considered as HGT. All gene trees identified as HGT were manually validated.

To evaluate phylogenetic relationship within 673 common proteins which were used in multi-locus genomic tree the previously obtained protein alignments were used. Particular phylogenetic trees were calculated in FastTreeMP (Price et al. 2010) with -lg –gamma settings. Acquired phylogenetic trees were visually checked and evaluated.

Exported phylogenetic trees were then drawn in Adobe Illustrator (CS6). Metabolic pathways were identified using KEGG BlastKOALA shown in KEGG pathway map (http://www.kegg.jp/) for the representative genome of *P. marinus* MIT 9313 (Kanehisa et al. 2016). Transmission electron microscopy (TEM) was performed as described earlier (Bohunicka et al. 2015).

## Results and Discussion

### Whole genome phylogeny

To infer the robust phylogeny of Cyanobacteria we calculated the concatenated phylogenetic tree based on 1,047 genes shared among the bacterial genomes (Fig. 1). For comparison we also calculated the standard 16S rRNA gene tree (Fig. 2) and a gene tree from the core set of 109 genes found in all 93 analyzed genomes (Supplementary Figure S2, Supplementary Table S4). Our 16S rRNA tree shared the same basic topology as shown in previous studies (Mareš et al. 2013; Shih et al. 2013; Schirrmeister et al. 2013) with *Gloeobacter* and thermophilic *Synechococcus* species placed as basal groups. The 109 gene tree showed *Gloeobacter* with high bootstrap support at the base of cyanobacterial phylogeny followed by thermal *Synechococcus* strains JA-3-3Ab and JA-2-3Ba next to the PSC clade. The topology of the concatenated tree based on 1,047 genes differed from the 16S rRNA gene tree remarkably. While most of the cyanobacteria clustered together, the *Prochlorococcus*, marine *Synechococcus* and *Cyanobium* species deviated from the main group forming a distinct, separate clade (Fig. 1). Other bacterial clades in the 1,047 gene tree were congruent with their standard classification (Madigan 2012, p. 357), supporting the plausibility of the inferred phylogenetic relations. The PSC clade mainly included marine species. *Prochlorococcus* spp. and marine “*Synechococcus*” species represent the most abundant component of marine phytoplankton (Flombaum et al. 2013). The PSC clade also contained species affiliated to the genus *Cyanobium* (Komarek et al. 1999).Interestingly while *Cyanobium gracile* PCC 6307 is a freshwater species, *Cyanobium* sp. PCC 7001 is a marine strain with a broad salinity tolerance (Stanier et al. 1971). *Prochlorococcus* strains MIT 9313 and MIT 9303 were phylogenetically more closely related to strains from the *Synechococcus* clade. According to the 16S rRNA gene phylogeny these two strains are located between the other *Prochlorococcus* strains and marine *Synechococcus* and are also shown in studies of multiple PSC strains resolved in *Prochlorococcus* clade IV (e.g. Rocap et al. 2002). The interlaid phylogeny of two strains of *Prochlorococcus* (MIT9303, MIT9313) and *Synechococcus* seems to be a frequent phenomenon which was shown before for various genes (Scanlan et al. 2009).

**Figure 1.**
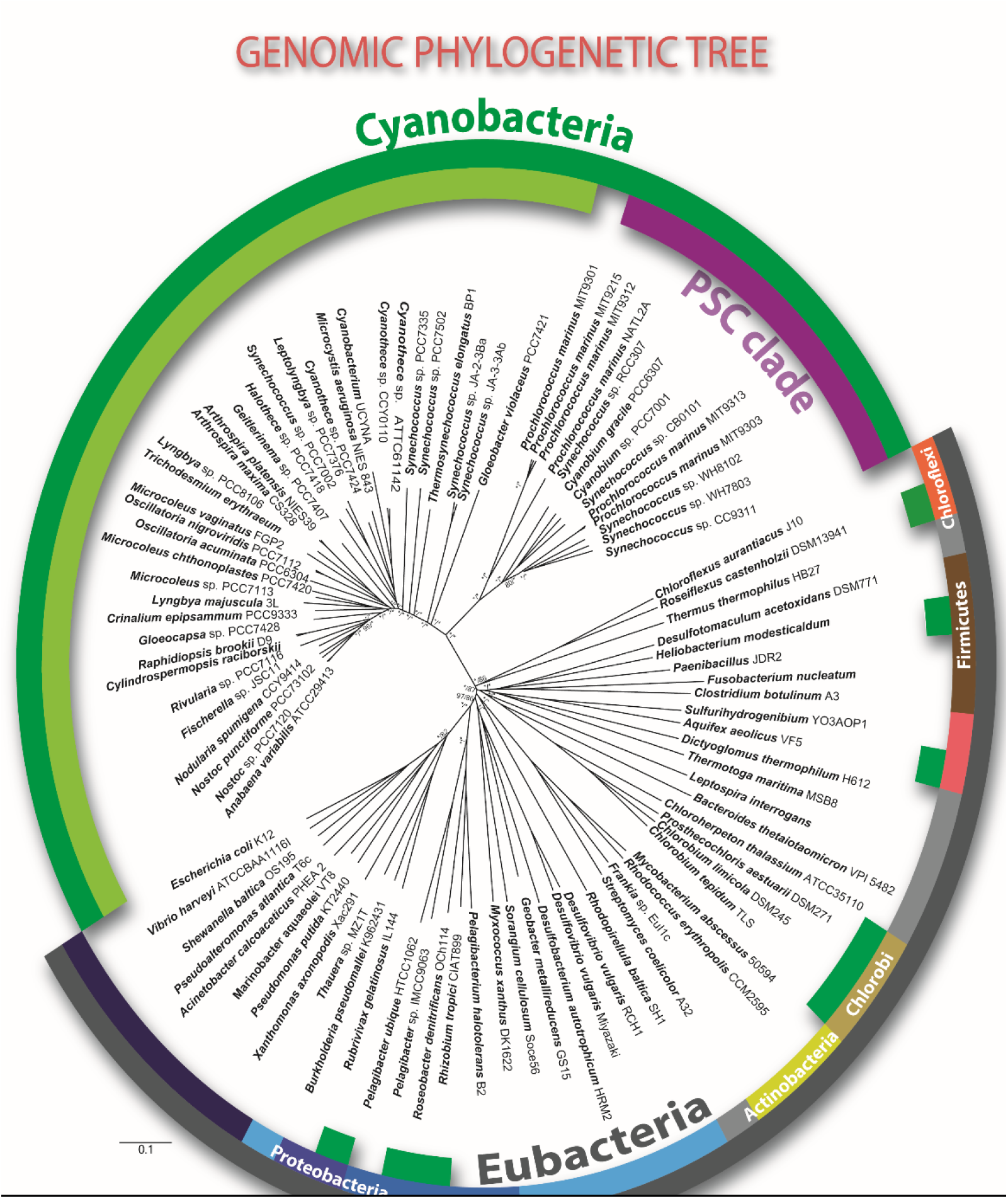
Whole genome phylogenetic analysis of selected members of Cyanobacteria and reference bacteria. The phylogenetic tree is based on concatenated amino acid sequences of 1,047 genes (406,617 aa length) shared among the selected species. The sector for Cyanobacteria is marked in

**Figure 2.**
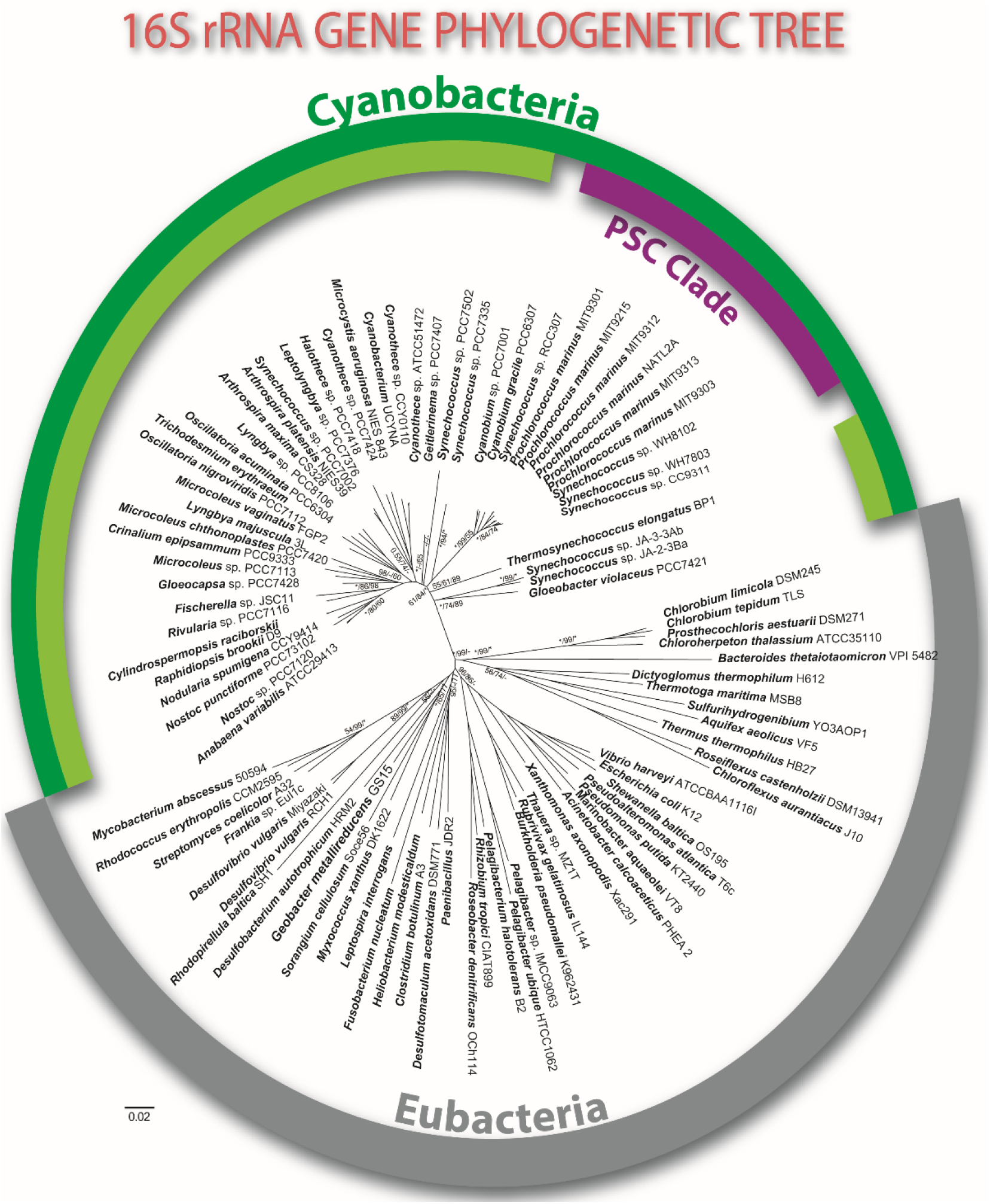
16S rRNA gene phylogenetic analysis shows traditional branching of several distinct cyanobacterial clades, *Gloeobacter* is located near the base of all the cyanobacterial lineages, followed by several clades designated mostly as *Synechococcus*. The PSC clade represents the special cluster next to Gloeobacterales, separated from the other coccoid and filamentous cyanobacterial taxa. Bayesian posterior analysis, Neighbor Joining and Maximum Likelihood bootstraps values with 1,000 repetitions are shown; values equal 1, respective 100% are indicated by asterisk.

In contrast to the whole genomic tree the phylogenic tree based on the core-set of 109 genes was more similar to the 16S RNA phylogeny. It however showed a smaller bootstrap support in basal clades of reference bacteria (Supplementary Figure S2). The similarity of these two phylogenic trees probably reflects the fact that the core-set contains mostly genes responsible for DNA replication and maintenance, transcription, ribosomal architecture and for enzymes of amino acid synthesis. Only 16 genes are responsible for other metabolic functions such glycolysis and the pentose phosphate cycle (Supplementary Table S4). These genes are considered conservative with slow evolution rates similar to the evolution rate of 16S rRNA.

The selection of genes for multiple gene phylogeny thus remains one of the crucial factors influencing tree topology at the basal part of cyanobacterial tree of life. The variability of inferred tree topologies for various sets of genes had already been shown in previous research (Capella-Gutierrez et al. 2014; Shih et al. 2013) and remains one of the crucial problems in cyanobacterial taxonomy.

### Unique characteristics of the PSC clade

The separation of the PSC clade from other cyanobacterial lineages reflects some remarkable differences between these two groups. Marine picocyanobacteria represent the species with the smallest cells (<1.5 μm). *Prochlorococcus* and marine *Synechococcus* share similar simple morphology and ultrastructure such as the isolated centroplasm distinct from the thylakoids at the cell’s periphery (Fig. 3). The ultrastructure of *Prochlorococcus* exhibits particularly radical minimization of the cell size (Fig. 3) and biochemical pathways (Supplementary Figure S3).

**Figure 3.**
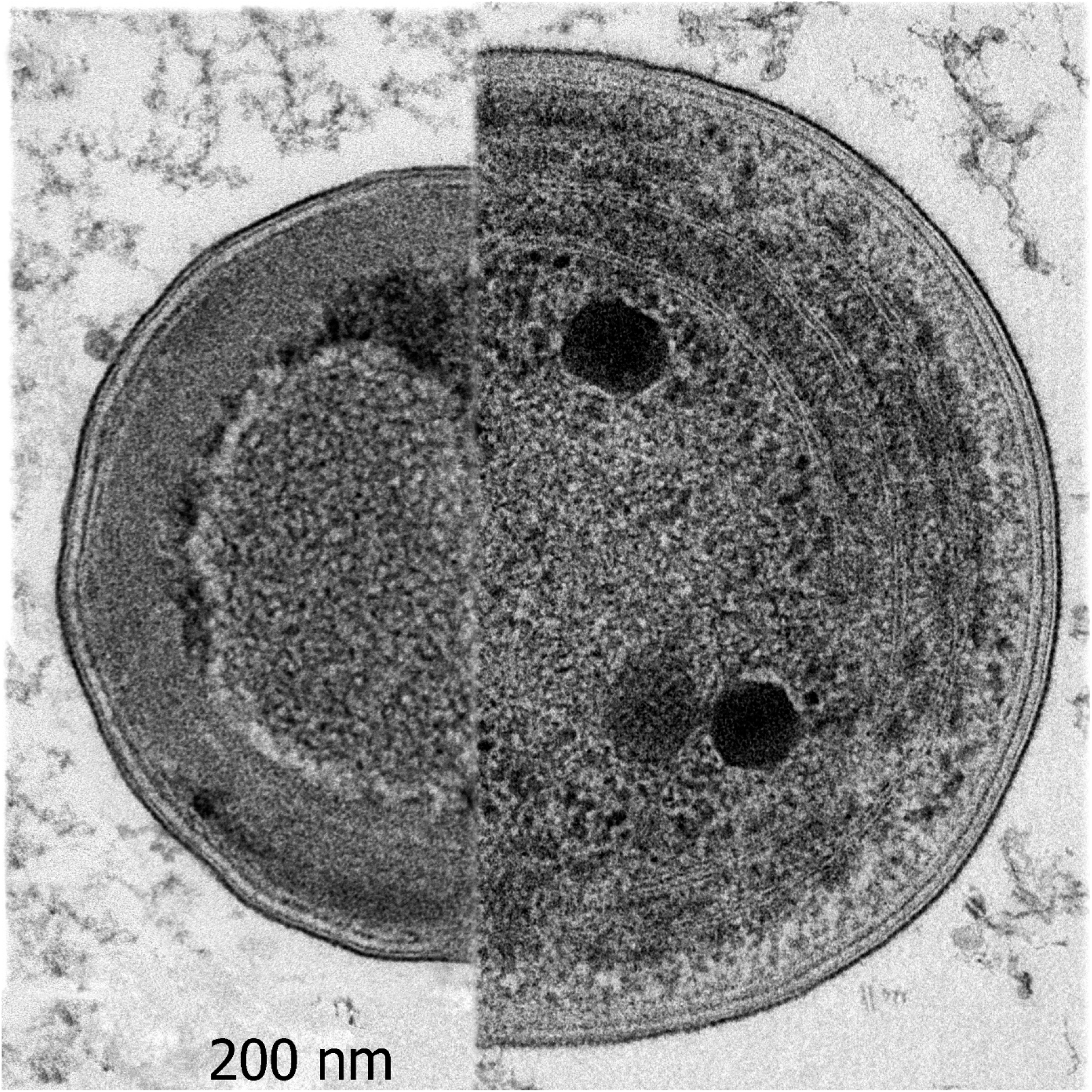
Composite image of *Prochlorococcus marinus* (left) and *Synechococcus marinus* (right). The ultrastructure of both species is similar, with a characteristic and distinct centroplasma (white arrow) with radial thylakoids (black arrow), accompanied with supreme minimalization of the cyanobacterial cell. α-type carboxysomes - marked by asterisk – can be distinguished in the *Synechococcus* cell.

Another noticeable feature is that the PSC species contain significantly smaller genomes in comparison with other cyanobacteria (Fig. 4). While most of the PSC species have genomes smaller than 3 Mbp (*Prochlorococcus* species 1.64–2.68 Mbp, marine *Synechococcus* 2.22–2.86 Mbp and *Cyanobium gracile* PCC 6307 2.83-3.34 Mbp) the genomes of the other cyanobacteria span from 3 to 9 Mbp (Hess 2011; Scanlan et al. 2009). The small genomes of *Prochlorococcus* have been usually interpreted as the effect of genomic streamlining (García-Fernández et al. 2004; Partensky and Garczarek 2010). It is highly likely that the same process occurred in the marine *Synechococcus* species genome.

**Figure 4.**
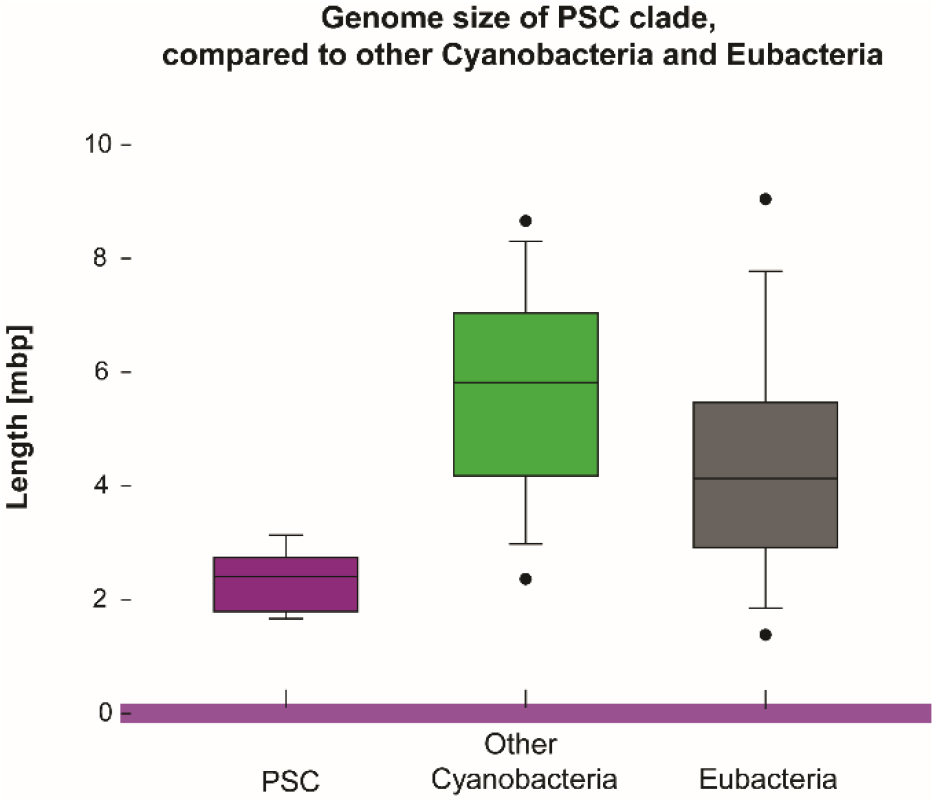
Comparison of genome sizes of the PSC clade, other Cyanobacteria and selected reference bacteria.

The members of the PSC clade are also unique in their photosynthetic apparatus. Since photosynthesis genes were not included in the full genome tree we built a concatenated phylogenetic tree from 17 protein subunits of Photosystem I and Photosystem II present in 45 cyanobacterial genomes. The obtained tree displayed the same separation of PSC clade from the rest of Cyanobacteria as the full genome phylogeny (Fig. 5).

**Figure 5.**
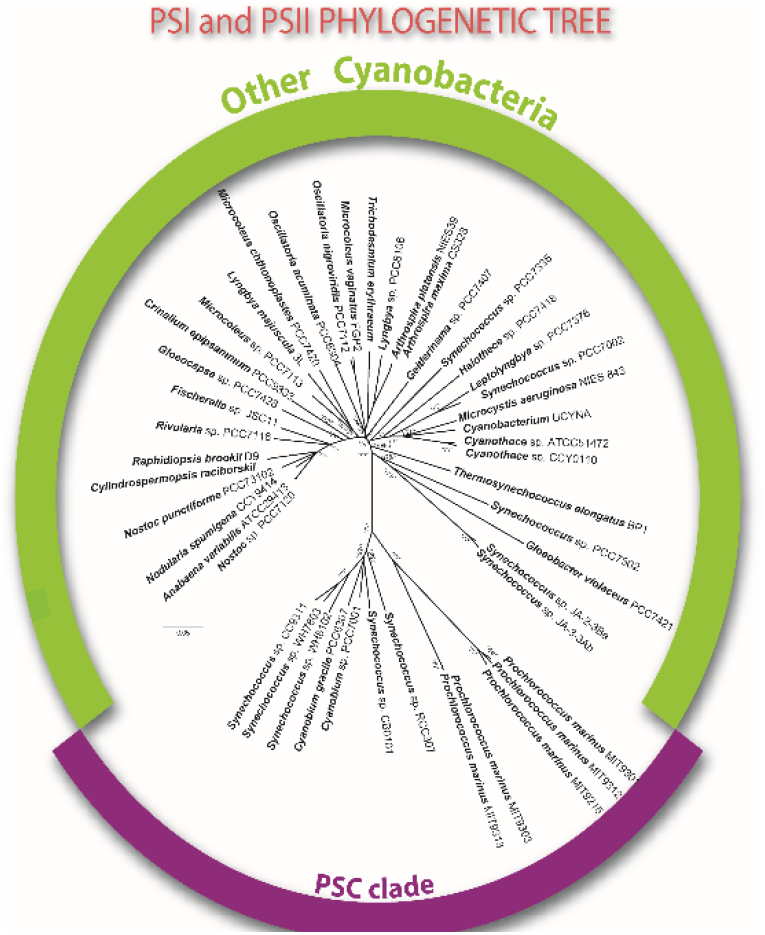
Phylogenetic analysis of 17 concatenated genes encoding various subunits of Photosystem I and Photosystem II. The deep split between PSC clade and other Cyanobacteria is shown. Bayesian posterior analysis, Neighbor Joining and Maximum Likelihood bootstraps values with 1,000 repetitions are shown; values equal 1, respective 100% are indicated by asterisk.

### Three evolutionary scenarios

The obtained whole genome phylogenetic tree indicates that marine picocyanobacteria and members of genus *Cyanobium* (PSC clade) significantly deviate from the other Cyanobacterial species. The separation of PSC clade from other Cyanobacterial lineages may originate from three possible evolutionary scenarios. (1) PSC clade originates from one of the early branches of Cyanobacteria, embarking on a very different evolutionary trajectory due to unique environmental conditions. (2) PSC lineage evolved as one of many Cyanobacterial lineages. The PSC clade however later adopted a large number of genes via HGT. The presence of many horizontally transferred genes distorted the whole genome phylogeny, but it had no influence on the 16S phylogeny. (3) PSC clade descended from a fusion of a primitive cyanobacterium with an ancient bacterium. This scenario implies a massive gene rearrangement which makes the genetic inventory of the present day PSC species very different from other Cyanobacteria.

### Extent of horizontal gene transfer

The presence of reference bacterial genes within cyanobacterial genomes has been noticed and discussed before (i.e. Shi and Falkowski 2008; Zhaxybayeva et al. 2006). To investigate the extent of horizontal gene transfer events in the PSC clade we analyzed individual phylogenies of all 2,269 genes present in *Prochlorococcus* sp. MIT9313. We identified 374 genes which were phylogenetically more related to various bacterial lineages than other Cyanobacteria (Supplementary Table S1). These potentially transferred genes were found in the complete set of studied PSC genomes with the only exception of four genes which were not found in *Cyanobium gracile*. The identified genes were phylogenetically related to Proteobacteria in 157 cases, Firmicutes in 125 cases and Actinobacteria and Chloroflexi in 33 and 11 cases respectively. These results undermine the plausibility of the fusion hypothesis (# 3) due to the only modest extent of the transferred genes and their variable phylogenetic origin.

Many of the genes which were identified as potentially horizontally transferred encode enzymes involved in important metabolic pathways (Supplementary Figure S3). An example of horizontally transferred genes are the enzymes of the Benson-Calvin cycle. Here a large subunit of RuBisCO, phosphoribulokinase, glyceraldehyde 3-phosphate dehydrogenase, fructose 1,6-bisphosphatase, glucose-6-phosphate isomerase, and sedoheptulose-1,7-bisphosphatase were found to be phylogenetically closer to reference bacteria than to other cyanobacteria (Supplementary Table S1). The close similarity of RuBisCO (large and small subunits), phosphoribulokinase, and fructose 1,6-bisphosphatase to Proteobacteria (such as *Thiobacillus* species) was already noted by Dufresne et al. (2003) in *P. marinus*. The difference in RuBisCO genes is manifested also at the microscopic level in the architecture of the carboxysomes. Hence, functional connectedness of RuBisCO is strongly related to carboxysome proteins itself and coevolved simultaneously. The difference of the carboxysome architecture between marine picocyanobacteria and other cyanobacterial species was already noted before (Rae et al. 2013). Differences in carbon fixation led Badger and Price (2003) to divide cyanobacteria into two groups according the protein systems needed for transport, concentration and fixation of dissolved inorganic carbon: α-cyanobacteria which correspond to the PSC clade, and β-cyanobacteria encompassing all the other cyanobacterial genera (Kerfeld and Melnicki 2016). A clear difference between PSC species and other cyanobacteria can also be documented in fundamental metabolic pathways such as the tricarboxylic acid cycle (TCA). TCA as a central pathway for recovering energy from major metabolites is substantially linked with production of sugars in the Benson-Calvin cycle.

While other Cyanobacteria use 2-oxoglutarate decarboxylase and succinate-semialdehyde dehydrogenase to convert 2-oxo-glutarate to succinate (Zhang and Bryant 2011) the PSC species have an additional 2-oxoglutarate dehydrogenase and simultaneously they lack the enzyme that converts succinyl-CoA to succinate or vice versa to close the TCA cycle. It is possible they can convert isocitrate to succinate via the glyoxylate shunt that was recently described by Zhang et al. (2016). Such differences from other cyanobacteria indicate a considerably modified operation of the TCA cycle.

Another example of horizontally transferred genes can be found in the chlorophyll-biosynthesis pathway. While most of the chlorophyll biosynthetic genes document common evolution of all Cyanobacteria, light-independent protochlorophyllide oxidoreductase (DPOR) shows a more complex evolution. The PSC clade species contain completely different forms of DPOR than the rest of the Cyanobacteria. The phylogenetic analysis showed that DPOR genes in PSC species are closely related to those found in purple bacteria (phototrophic Proteobacteria) indicating their ancient horizontal transfer between PSC group and Proteobacteria (Gupta 2012) or in the opposite direction (Zeng et al. 2014).

High sequence similarity of a gene in a particular organism to one present in a representative of a phylogenetically distant group is considered as telltale sign of HGT. Enzymatic complexes composed of several protein subunits (often topologically distant within the genome) are highly unlikely to be encoded by a mix of indigenous and HGT genes. Perturbation of weak intermolecular interactions associated with the adoption of an alternative protein subunit (through HGT) will most likely result in a change in the amino acid residues at the protein-protein interface of the protein subunits. Even minor changes may result in a malfunction of the multi-subunit enzymatic complex, whose functionality is indispensable for the organism’s survival at any point during its evolution. This holds especially true for the enzymes of the core metabolism. Contrary to this expectation, subunits of multiple key enzymes in species of the PSC clade are suspected to be contracted through HGT as they are more related to the other bacteria than to the cyanobacterial variants (Supplemental Table 4). These aforementioned enzymes are not only playing part in the core metabolism (Supplementary Figure S3), but also take part in carbon fixation, chlorophyll synthesis etc. The representatives of PSC clade include both marine species and freshwater species, that are not prone to the strong viral HGT pressure. Then the rational consequence is that many of genes considered as HGT might simply originate from common ancestor of cyanobacteria and reference bacteria remaining unchanged in the PSC lineage. However, this hypothesis of frozen evolution should be elucidated by further more focused studies on this topic.

### Whole genome phylogeny without HGT genes

To evaluate the influence of HGT on the whole genome phylogeny we calculated the new concatenated tree without the genes identified as potentially horizontally transferred. Interestingly the new tree constructed from 673 concatenated genes displayed very similar overall topology as the original whole genome tree. The PSC clade formed the same basal clade to all other cyanobacteria (Supplementary Figure S1). Only minor rearrangements were inferred within other cyanobacteria with repositioning of the *Prochlorococcus* strains MIT 9313 and 9303 back into intermediate position between *Prochlorococcus* and *Synechococcus* strains.

Comparison of the genomic tree without the putative horizontally transferred genes with two other phylogenic trees inferred from 16S rRNA and from the core-set of 109 genes found in all studied genomes raises questions about rate of evolution within different sets of genes. Different rates of evolution (Whelan et al. 2011) among 109 genes responsible for DNA and RNA management and protein synthesis and the other genes more related to environmental conditions in oceans may explain the differences in phylogenies (Supplementary Table S4). Rates of evolution can be derived from the length of branches in phylogenetic trees showing a relatively similar number of mutations through whole phylogeny of cyanobacteria. If substantial heterotachy occurs in PSC clade we would expect a much longer branches in this group which were not found. If the mutation rate in PSC group was faster than the rest of cyanobacteria we would expect that its phylogenetic position would be more distant from bacteria, however, this was not the case.

### Analysis of non HGT genes

To further understand the function and phylogenetic position of particular genes that were not considered as potentially HGT we used 673 single phylogenetic trees which were not previously sorted out as HGT by comparison with non-redundant GenBank protein database. To simplify the evaluation of a phylogenetic signal we split them into three groups. The first group contained phylogenetic trees with *Gloeobacter* closer to reference bacterial outgroup than PSC clade which had 29% of total trees. We also identified 17 phylogenetic trees with *Gloeobacter* inside the reference bacterial lineages suggesting HGT to *Gloeobacter* (Supplementary Table 5). The second group with PSC group basal to *Gloeobacter* included 38% of phylogenetic trees. The last group contained no outgroup (i.e. no reference bacteria) and contained 32% of 673 evaluated gene trees (Supplementary Table 5). The association of many proteins to particular metabolic pathways from first and second group is ambiguous, however, several assumptions can be made. *Gloeobacter* lineage is phylogenetically closer to reference bacteria in proteins responsible for DNA repair and replication, synthesis of tRNA, mRNA and rRNA as well as other proteins used in transcription. Proteins responsible for RNA, heme and protein degradation also fall into this category (Supplementary Table S5). Proteins of PSC clade which are phylogenetically closer to reference bacteria than *Gloeobacter* belong to many various metabolic pathways related to reactions concerning nitrogen, sulfur, iron, molybdenum, cobalt or zinc (Supplementary Table S5). They comprise sulfate and ferric transporters, regulators and numerous other symporters, antiporters and permeases. Nitrogen regulatory proteins can be found exclusively in this group as well as four proteins involved in urea transport and metabolism. Proteins for urea utilization, which might be important source of nitrogen in oceans, complement the metabolic pathway that was already identified as potential HGT (Supplementary Figure S3) by García-Fernández et al. (2004). Proteins taking part in circadian clock machinery as well as proteins involved in cell division machinery were predominantly found in phylogenetic trees with PSC being closer to reference bacteria than to *Gloeobacter*.

Third category of phylogenetic trees with no outgroup reference bacteria naturally comprise many genes connected with photosynthetic pathways. This includes proteins connected to thylakoid synthesis, the transport of substrates and management of photosynthetic products (Supplementary Table S5).

## Conclusion

These results document that the separation of the PSC clade within the whole genomic tree is not caused by the presence of horizontally transferred genes (#2). Despite the fact that HGT events were identified in genomes of PSC species (Supplementary Figure S3) the impact of this process on whole genome phylogeny of Cyanobacteria was only minimal, therefore, the phylogenetic position of the PSC species reflects a fundamental historical separation (Dvořák et al. 2014) from the all other Cyanobacteria (#1). Evolution of non-marine cyanobacteria in many diverse environments allowed for further specialization and rapid speciation. This is clearly reflected in much larger genomes of non-marine Cyanobacteria (Fig. 4). In contrast stable conditions in the oligotrophic oceans kept rates of speciation low due to constant genome optimization and streamlining. Selection pressure on PSC genomes was much higher causing reduced rates of mutation and specialization.

Individual genes are predominantly sequentially static and are already best tuned for oligotrophic and stable environments with rare beneficial mutations.

The PSC clade most likely represents an isolated cyanobacterial lineage which followed a distinct evolutionary trajectory which was driven by its specific environment.

