## Supplemental for "Whole genome phylogeny of Cyanobacteria documents a distinct evolutionary trajectory of marine picocyanobacteria": Supplementary_Table_S1.docx

| Supplementary_Table_S1- Potential HGT genes  Genes phylogenetically more related to various Eubacteria than other Cyanobacteria. | | | | | | | |
| --- | --- | --- | --- | --- | --- | --- | --- |
| Genebank accesion numbers for *P. marinus* str. MIT 9313 | Not in Cyano-bium | Gene | Phylum | Class | Order | Family | Genus |
| 33862583 |  | glycosyltransferase | Firmicutes | Bacilli | Bacillales | Alicyclobacillaceae | Alicyclobacillus |
| 33862584 |  | galactosyl-1-phosphate transferase | Acidobacteria | Holophagae | Holophagales | Holophagaceae | Holophaga |
| 33862585 |  | cobalamin biosynthetic protein | Actinobacteria | Actinobacteria | Actinomycetales | Micromonosporaceae | Actinoplanes |
| 33862586 |  | ketol-acid reductoisomerase | Firmicutes | Bacilli | Bacillales | Alicyclobacillaceae | Alicyclobacillus |
| 33862589 |  | PIN domain superfamily protein | Bacteroidetes | Flavobacteriia | Flavobacteriales | Flavobacteriaceae | Bergeyella |
| 33862591 |  | 3-methyl-2-oxobutanoate hydroxymethyltransferase | Firmicutes | Clostridia | Clostridiales | Peptococcaceae | Desulfosporosinus |
| 33862600 |  | FAD/FMN-containing dehydrogenase | Firmicutes | Clostridia | Thermoanaerobacterales | Thermoanaerobacteraceae | Thermoanaerobacter |
| 33862603 |  | bifunctional dihydrofolate/folylpolyglutamate synthase | Proteobacteria | Gammaproteobacteria | Acidithiobacillales | Acidithiobacillaceae | Acidithiobacillus |
| 33862604 |  | acetylornithine aminotransferase | Firmicutes | Clostridia | Clostridiales | Peptococcaceae | Desulfosporosinus |
| 33862608 |  | dihydrolipoamide dehydrogenase | Firmicutes | Bacilli | Bacillales | Paenibacillaceae | Paenibacillus |
| 33862609 |  | indole-3-glycerol-phosphate synthase | Proteobacteria | Betaproteobacteria | Rhodocyclales | Rhodocyclaceae | Azoarcus |
| 33862614 |  | FKBP-type peptidyl-prolyl cis-trans isomerase (PPIase) | Synergistetes | Synergistia | Synergistales | Synergistaceae | Anaerobaculum |
| 33862618 |  | hypothetical protein PMT0345 | Proteobacteria | Alphaproteobacteria | Rhizobiales | Rhizobiaceae | Agrobacterium |
| 33862620 |  | pyruvate dehydrogenase E1 alpha subunit | Firmicutes | Bacilli | Bacillales | Bacillaceae | Bacillus |
| 33862628 |  | GTP-binding protein Era | Bacteroidetes | Flavobacteriia | Flavobacteriales | Flavobacteriaceae | Capnocytophaga |
| 33862637 |  | riboflavin kinase/FAD synthase | Proteobacteria | Deltaproteobacteria | Myxococcales | Polyangiaceae | Sorangium |
| 33862642 |  | hypothetical protein PMT0369 | Firmicutes | Bacilli | Bacillales | Bacillaceae | Caldalkalibacillus |
| 33862643 |  | precorrin-6y methylase | Firmicutes | Bacilli | Bacillales | Bacillaceae | Anoxybacillus |
| 33862648 |  | bifunctional diaminohydroxyphosphoribosylaminopyrimidine deaminase/5-amino-6-(5-phosphoribosylamino)uracil reductase | Proteobacteria | Alphaproteobacteria | Rhodospirillales | Rhodospirillaceae | Azospirillum |
| 33862650 |  | hypothetical protein PMT0377 | Firmicutes | Bacilli | Bacillales | Paenibacillaceae | Paenibacillus |
| 33862652 |  | ornithine carbamoyltransferase | Proteobacteria | Betaproteobacteria | Burkholderiales | Alcaligenaceae | Bordetella |
| 33862662 |  | acetyl-coenzyme A synthetase | Proteobacteria | Betaproteobacteria | Neisseriales | Neisseriaceae | Kingella |
| 33862666 |  | aspartate and glutamate racemase:glutamate racemase | Proteobacteria | Gammaproteobacteria | Methylococcales | Methylococcaceae | Methylomicrobium |
| 33862668 |  | nitrilase | Chloroflexi | Chloroflexi | Chloroflexales | Chloroflexaceae | Roseiflexus |
| 33862670 |  | hypothetical protein PMT0397 | Firmicutes | Negativicutes | Selenomonadales | Veillonellaceae | Veillonella |
| 33862673 |  | bifunctional N-acetylglucosamine-1-phosphate uridyltransferase/glucosamine-1-phosphate acetyltransferase | Firmicutes | Bacilli | Bacillales | Alicyclobacillaceae | Alicyclobacillus |
| 33862678 |  | naphthoate synthase | Firmicutes | Bacilli | Lactobacillales | Carnobacteriaceae | Carnobacterium |
| 33862679 |  | menaquinone biosynthesis 2-succinyl-6-hydroxy-2,4-cyclohexadiene-1-carboxylate synthase | Actinobacteria | Actinobacteria | Actinomycetales | Nocardiaceae | Rhodococcus |
| 33862690 |  | ABC transporter ATP-binding protein | Proteobacteria | Gammaproteobacteria | Thiotrichales | Piscirickettsiaceae | Methylophaga |
| 33862697 |  | threonyl-tRNA synthetase | Proteobacteria | Gammaproteobacteria | Alteromonadales | Shewanellaceae | Shewanella |
| 33862701 |  | phosphofructokinase | Actinobacteria | Actinobacteria | Actinomycetales | Frankiaceae | Frankia |
| 33862729 |  | RNA polymerase sigma-70 factor family protein | Firmicutes | Clostridia | Thermoanaerobacterales | Thermoanaerobacteraceae | Ammonifex |
| 33862738 |  | tryptophan synthase subunit alpha | Planctomycetes | Phycisphaerae | Phycisphaerales | Phycisphaeraceae | Phycisphaera |
| 33862742 |  | CaCA family, sodium/calcium exchanger | Proteobacteria | Gammaproteobacteria | Chromatiales | Chromatiaceae | Marichromatium |
| 33862743 |  | glutathione reductase (NADPH) | Actinobacteria | Actinobacteria | Actinomycetales | Micromonosporaceae | Micromonospora |
| 33862765 |  | peptide methionine sulfoxide reductase | Proteobacteria | Betaproteobacteria | Burkholderiales | Comamonadaceae | Delftia |
| 33862774 |  | ABC transporter ATP-binding protein | Planctomycetes | Planctomycetia | Planctomycetales | Planctomycetaceae | Rhodopirellula |
| 33862777 |  | endonuclease | Chlorobi | Chlorobia | Chlorobiales | Chlorobiaceae | Chlorobium |
| 33862793 |  | 1-(5-phosphoribosyl)-5-[(5-phosphoribosylamino)methylideneamino] imidazole-4-carboxamide isomerase | Proteobacteria | Gammaproteobacteria | Pseudomonadales | Pseudomonadaceae | Pseudomonas |
| 33862802 |  | gamma-glutamyl kinase | Firmicutes | Bacilli | Lactobacillales | Enterococcaceae | Enterococcus |
| 33862807 |  | acetyl-CoA carboxylase subunit beta | Firmicutes | Bacilli | Bacillales | Bacillaceae | Bacillus |
| 33862812 |  | phosphoribosylformylglycinamidine synthase I | Proteobacteria | Betaproteobacteria | Burkholderiales | Comamonadaceae | Albidiferax |
| 33862814 |  | oxidoreductase, Fe-S subunit | Firmicutes | Bacilli | Bacillales | Alicyclobacillaceae | Alicyclobacillus |
| 33862816 |  | glycosyl transferase family protein | Firmicutes | Bacilli | Bacillales | Bacillaceae | Bacillus |
| 33862826 |  | glycine betaine/proline ABC transporter ATP-binding protein | Proteobacteria | Alphaproteobacteria | Rhodospirillales | Acetobacteraceae | Acidiphilium |
| 33862833 |  | dihydroxy-acid dehydratase | Proteobacteria | Gammaproteobacteria | Enterobacteriales | Enterobacteriaceae | Plesiomonas |
| 33862837 |  | 6-phosphogluconolactonase | Firmicutes | Clostridia | Clostridiales | Peptococcaceae | Desulfitobacterium |
| 33862840 |  | glutamyl-tRNA reductase | Firmicutes | Bacilli | Bacillales | Bacillaceae | Bacillus |
| 33862841 |  | fructose 1,6-bisphosphatase II | Firmicutes | Bacilli | Bacillales | Paenibacillaceae | Brevibacillus |
| 33862842 |  | ribulose-phosphate 3-epimerase | Fibrobacteres | Fibrobacteria | Fibrobacterales | Fibrobacteraceae | Fibrobacter |
| 33862845 |  | ABC transporter ATP-binding protein | Chloroflexi | Chloroflexia | Chloroflexales | Roseiflexaceae | Roseiflexus |
| 33862850 |  | aromatic-ring hydroxylase (flavoprotein monooxygenase) | Proteobacteria | Deltaproteobacteria | Desulfuromonadales | Geobacteraceae | Geobacter |
| 33862851 |  | glycyl-tRNA synthetase subunit beta | Firmicutes | Bacilli | Bacillales | Bacillaceae | Bacillus |
| 33862858 |  | glycosyl transferase family protein | Proteobacteria | Alphaproteobacteria | Rhodospirillales | Acetobacteraceae | Roseomonas |
| 33862860 |  | ABC transporter ATP-binding protein | Proteobacteria | Deltaproteobacteria | Bdellovibrionales | Bdellovibrionaceae | Bdellovibrio |
| 33862864 |  | Zn-dependent peptidase | Proteobacteria | Alphaproteobacteria | Rhodospirillales | Rhodospirillaceae | Azospirillum |
| 33862865 |  | insulinase family protein | Actinobacteria | Actinobacteria | Bifidobacteriales | Bifidobacteriaceae | Gardnerella |
| 33862872 |  | cytidine/deoxycytidylate deaminase | Proteobacteria | Gammaproteobacteria | Vibrionales | Vibrionaceae | Vibrio |
| 33862891 |  | pyruvate dehydrogenase E1 beta subunit | Proteobacteria | Gammaproteobacteria | 0 | 0 | Congregibacter |
| 33862896 |  | glutamate dehydrogenase | Firmicutes | Negativicutes | Selenomonadales | Veillonellaceae | Veillonella |
| 33862914 |  | DNA primase | Actinobacteria | Actinobacteria | Bifidobacteriales | Bifidobacteriaceae | Bifidobacterium |
| 33862918 |  | 30S ribosomal protein S15 | Firmicutes | Clostridia | Thermoanaerobacterales | Thermoanaerobacteraceae | Caldanaerobacter |
| 33862921 |  | aspartyl/glutamyl-tRNA amidotransferase subunit A | Actinobacteria | Actinobacteria | Actinomycetales | Actinomycetaceae | Actinomyces |
| 33862923 |  | tRNA/rRNA methyltransferase SpoU | Proteobacteria | Deltaproteobacteria | Desulfobacterales | Desulfobacteraceae | Desulfobacula |
| 33862928 |  | multidrug ABC transporter | Proteobacteria | Alphaproteobacteria | Sphingomonadales | Sphingomonadaceae | Sphingobium |
| 33862934 |  | quinolinate synthetase | Proteobacteria | Betaproteobacteria | Rhodocyclales | Rhodocyclaceae | Azoarcus |
| 33862956 |  | threonine dehydratase | Proteobacteria | Gammaproteobacteria | Chromatiales | Chromatiaceae | Marichromatium |
| 33862957 |  | 1-deoxy-D-xylulose-5-phosphate synthase | Firmicutes | Bacilli | Lactobacillales | Lactobacillaceae | Lactobacillus |
| 33862965 |  | sugar ABC transporter | Chlamydiae | Chlamydiia | Chlamydiales | Parachlamydiaceae | Parachlamydia |
| 33862968 |  | heat shock protein 90 | Proteobacteria | Deltaproteobacteria | Bdellovibrionales | Bdellovibrionaceae | Bdellovibrio |
| 33862971 |  | inositol phosphatase/fructose-1,6-bisphosphatase | Actinobacteria | Actinobacteria | Actinomycetales | Nocardioidaceae | Aeromicrobium |
| 33862972 |  | phosphate ABC transporter ATP-binding protein | Proteobacteria | Gammaproteobacteria | Enterobacteriales | Enterobacteriaceae | Salmonella |
| 33862973 |  | phosphate ABC transporter | Proteobacteria | Gammaproteobacteria | Pseudomonadales | Pseudomonadaceae | Pseudomonas |
| 33862974 |  | phosphate ABC transporter | Firmicutes | Bacilli | Bacillales | Bacillaceae | Bacillus |
| 33862978 |  | N-acetylmuramic acid 6-phosphate etherase | Proteobacteria | Epsilonproteobacteria | Campylobacterales | Campylobacteraceae | Campylobacter |
| 33862981 |  | N-acetyl-gamma-glutamyl-phosphate reductase | Proteobacteria | Gammaproteobacteria | Oceanospirillales | Alcanivoracaceae | Alcanivorax |
| 33862982 |  | phosphoribosylglycinamide formyltransferase | Proteobacteria | Betaproteobacteria | Burkholderiales | Comamonadaceae | Albidiferax |
| 33862983 |  | PDZ domain-containing protein | Proteobacteria | Alphaproteobacteria | Rhodospirillales | Acetobacteraceae | Acidocella |
| 33862986 |  | glucose-6-phosphate isomerase | Firmicutes | Negativicutes | Selenomonadales | Veillonellaceae | Megamonas |
| 33862987 |  | leucyl-tRNA synthetase | Proteobacteria | Gammaproteobacteria | Alteromonadales | Shewanellaceae | Shewanella |
| 33862994 |  | phosphopantetheine adenylyltransferase | Actinobacteria | Actinobacteria | Actinomycetales | Mycobacteriaceae | Mycobacterium |
| 33863010 |  | phenylalanyl-tRNA synthetase subunit beta | Chlorobi | Chlorobia | Chlorobiales | Chlorobiaceae | Chlorobium |
| 33863015 |  | methionyl-tRNA synthetase | Actinobacteria | Actinobacteria | Actinomycetales | Mycobacteriaceae | Mycobacterium |
| 33863021 |  | cobinamide kinase | Proteobacteria | Deltaproteobacteria | Desulfovibrionales | Desulfovibrionaceae | Desulfovibrio |
| 33863022 |  | SpoU family tRNA/rRNA methyltransferase | Firmicutes | Bacilli | Bacillales | Bacillaceae | Amphibacillus |
| 33863024 |  | LysM motif-containing protein | Nitrospinae | Nitrospinia | Nitrospinales | Nitrospinaceae | Nitrospina |
| 33863026 |  | thioredoxin peroxidase | Firmicutes | Clostridia | Clostridiales | Clostridiaceae | Clostridium |
| 33863044 |  | methionyl-tRNA formyltransferase | Firmicutes | Clostridia | Clostridiales | Clostridiaceae | Clostridium |
| 33863048 |  | carboxyl-terminal processing protease | Synergistetes | Synergistia | Synergistales | Synergistaceae | Anaerobaculum |
| 33863050 |  | uracil-DNA glycosylase | Archaea | Thaumarchaeota | Nitrosopumilales | Nitrosopumilaceae | Nitrosopumilus |
| 33863073 |  | flavodoxin FldA | Proteobacteria | Betaproteobacteria | Burkholderiales | Burkholderiaceae | Burkholderia |
| 33863077 |  | two-component response regulator | Actinobacteria | Actinobacteria | Actinomycetales | Pseudonocardiaceae | Actinosynnema |
| 33863088 |  | dihydropteroate synthase | Firmicutes | Clostridia | Thermoanaerobacterales | Thermoanaerobacteraceae | Thermoanaerobacter |
| 33863089 |  | triosephosphate isomerase | Firmicutes | Negativicutes | Selenomonadales | Veillonellaceae | Selenomonas |
| 33863091 |  | ABC transporter | Proteobacteria | Betaproteobacteria | Burkholderiales | Alcaligenaceae | Achromobacter |
| 33863094 |  | carbamoyl phosphate synthase large subunit | Firmicutes | Clostridia | Clostridiales | Clostridiaceae | Clostridium |
| 33863097 |  | L-asparaginase II | Firmicutes | Clostridia | Clostridiales | Peptococcaceae | Desulfotomaculum |
| 33863147 |  | O-Acetyl homoserine sulfhydrylase | Deinococcus | Deinococci | Deinococcales | Deinococcaceae | Deinococcus |
| 33863168 |  | amino acid ABC transporter | Proteobacteria | Alphaproteobacteria | Rhodospirillales | Rhodospirillaceae | Azospirillum |
| 33863169 |  | amino acid ABC transporter substrate-binding protein | Proteobacteria | Alphaproteobacteria | Rhodospirillales | Rhodospirillaceae | Tistrella |
| 33863173 |  | hypothetical protein PMT0901 | Firmicutes | Bacilli | Bacillales | Bacillaceae | Bacillus |
| 33863576 |  | glutamyl-tRNA synthetase | Firmicutes | Clostridia | Clostridiales | Lachnospiraceae | Blautia |
| 33863583 |  | DNA-binding/iron metalloprotein/AP endonuclease | Firmicutes | Bacilli | Bacillales | Bacillaceae | Bacillus |
| 33863591 |  | prolipoprotein diacylglyceryl transferase | Chlorobi | Chlorobia | Chlorobiales | Chlorobiaceae | Chlorobaculum |
| 33863594 |  | exopolyphosphatase | Firmicutes | Bacilli | Lactobacillales | Carnobacteriaceae | Carnobacterium |
| 33863597 |  | 2-C-methyl-D-erythritol 4-phosphate cytidylyltransferase | Firmicutes | Clostridia | Clostridiales | Clostridiaceae | Clostridium |
| 33863600 |  | 3-oxoacyl-ACP reductase | Firmicutes | Bacilli | Bacillales | Bacillaceae | Caldalkalibacillus |
| 33863601 |  | chaperonin GroEL | Proteobacteria | Gammaproteobacteria | Vibrionales | Vibrionaceae | Aliivibrio |
| 33863602 |  | N-acetylmannosamine-6-phosphate 2-epimerase | Actinobacteria | Actinobacteria | Actinomycetales | Micromonosporaceae | Actinoplanes |
| 33863605 |  | multidrug efflux ABC transporter | Spirochaetes | Spirochaetia | Spirochaetales | Leptospiraceae | Leptospira |
| 33863606 |  | protoheme IX farnesyltransferase | Proteobacteria | Gammaproteobacteria | Xanthomonadales | Xanthomonadaceae | Xanthomonas |
| 33863613 |  | riboflavin synthase subunit alpha | Chloroflexi | Chloroflexi | Chloroflexales | Chloroflexaceae | Chloroflexus |
| 33863625 |  | M23/M37 familypeptidase | Firmicutes | Bacilli | Bacillales | Bacillaceae | Bacillus |
| 16135004 |  | seryl-tRNA synthetase | Proteobacteria | Deltaproteobacteria | Myxococcales | Kofleriaceae | Haliangium |
| 33863645 |  | 3'-phosphoadenosine-5'-phosphosulfate (PAPS) 3'-phosphatase | Firmicutes | Clostridia | Clostridiales | Clostridiaceae | Clostridium |
| 33863646 |  | tetrapyrrole methylase family protein | Actinobacteria | Actinobacteria | Actinomycetales | Nocardioidaceae | Aeromicrobium |
| 33863661 |  | glutamyl/glutaminyl-tRNA synthetase | Chloroflexi | Chloroflexi | Chloroflexales | Chloroflexaceae | Roseiflexus |
| 33863666 |  | transporter membrane component | Firmicutes | Bacilli | Bacillales | Bacillaceae | Bacillus |
| 33863668 |  | membrane protein | Synergistetes | Synergistia | Synergistales | Synergistaceae | Anaerobaculum |
| 33863674 |  | leucyl aminopeptidase | Proteobacteria | Betaproteobacteria | Methylophilales | Methylophilaceae | Methylobacillus |
| 33863678 |  | (3R)-hydroxymyristoyl-ACP dehydratase | Bacteroidetes | Flavobacteriia | Flavobacteriales | Flavobacteriaceae | Bergeyella |
| 33863681 |  | phosphoribosylaminoimidazole-succinocarboxamide synthase | Firmicutes | Clostridia | Clostridiales | Peptococcaceae | Desulfitobacterium |
| 33863683 |  | phosphoribosylamine--glycine ligase | Firmicutes | Clostridia | Clostridiales | Clostridiaceae | Clostridium |
| 33863684 |  | two-component sensor histidine kinase | Actinobacteria | Actinobacteria | Actinomycetales | Mycobacteriaceae | Mycobacterium |
| 33863689 |  | tRNA pseudouridine synthase B | Synergistetes | Synergistia | Synergistales | Synergistaceae | Anaerobaculum |
| 33863690 |  | hypothetical protein PMT1423 | Firmicutes | Bacilli | Lactobacillales | Lactobacillaceae | Lactobacillus |
| 33863697 |  | 50S ribosomal protein L11 methyltransferase | Firmicutes | Bacilli | Bacillales | Bacillaceae | Halobacillus |
| 33863701 |  | UDP-N-acetylmuramoyl-L-alanyl-D-glutamate synthetase | Proteobacteria | Gammaproteobacteria | Cardiobacteriales | Cardiobacteriaceae | Cardiobacterium |
| 33863706 |  | hypothetical protein PMT1439 | Actinobacteria | Actinobacteria | Actinomycetales | Corynebacteriaceae | Corynebacterium |
| 33863712 |  | bifunctional pyrimidine regulatory protein PyrR/uracil phosphoribosyltransferase | Proteobacteria | Gammaproteobacteria | Alteromonadales | Alteromonadaceae | Alishewanella |
| 33863713 |  | phosphoglyceromutase | Chloroflexi | Chloroflexi | Chloroflexales | Chloroflexaceae | Chloroflexus |
| 33863716 |  | chaperonin GroEL | Actinobacteria | Actinobacteria | Acidimicrobiales | Acidimicrobiaceae | Ilumatobacter |
| 33863729 |  | alanine dehydrogenase | Firmicutes | Bacilli | Bacillales | Bacillaceae | Bacillus |
| 33863733 |  | F0F1 ATP synthase subunit gamma | Proteobacteria | Epsilonproteobacteria | Campylobacterales | Campylobacteraceae | Campylobacter |
| 33863745 |  | 7-cyano-7-deazaguanine reductase | Firmicutes | Negativicutes | Selenomonadales | Acidaminococcaceae | Acidaminococcus |
| 33863749 |  | rRNA methyltransferase | Proteobacteria | Gammaproteobacteria | Chromatiales | Ectothiorhodospiraceae | Nitrococcus |
| 33863760 |  | diaminopelargonic acid synthase | Firmicutes | Bacilli | Lactobacillales | Lactobacillaceae | Lactobacillus |
| 33863763 |  | 16S rRNA methyltransferase GidB | Firmicutes | Clostridia | Halanaerobiales | Halobacteroidaceae | Acetohalobium |
| 33863770 |  | ribosomal RNA large subunit methyltransferase N | Proteobacteria | Alphaproteobacteria | Sphingomonadales | Erythrobacteraceae | Erythrobacter |
| 33863813 |  | DNA mismatch repair protein MutS family protein | Proteobacteria | Deltaproteobacteria | Myxococcales | Nannocystaceae | Plesiocystis |
| 33863814 |  | delta-aminolevulinic acid dehydratase | Planctomycetes | Planctomycetia | Planctomycetales | Planctomycetaceae | Rhodopirellula |
| 33863845 |  | RNA methylase | Proteobacteria | Gammaproteobacteria | Chromatiales | Chromatiaceae | Thiocapsa |
| 33863863 |  | HIT (histidine triad) family protein | Proteobacteria | Gammaproteobacteria | Thiotrichales | Thiotrichaceae | Beggiatoa |
| 33863865 |  | ATPase AAA | Proteobacteria | Alphaproteobacteria | Rhizobiales | Beijerinckiaceae | Beijerinckia |
| 33863871 |  | ABC transporter | Proteobacteria | Gammaproteobacteria | Enterobacteriales | Enterobacteriaceae | Escherichia |
| 33862402 |  | RND family multidrug efflux transporter | Proteobacteria | Alphaproteobacteria | Rhodospirillales | Rhodospirillaceae | Caenispirillum |
| 33863884 |  | hypothetical protein PMT1617 | Chloroflexi | Caldilineae | Caldilineales | Caldilineaceae | Caldilinea |
| 33863885 |  | recombination factor protein RarA/unknown domain fusion protein | Actinobacteria | Actinobacteria | Coriobacteriales | Coriobacteriaceae | Eggerthella |
| 33863893 |  | GTP-binding protein, transport associated | Firmicutes | Clostridia | Halanaerobiales | Halobacteroidaceae | Halobacteroides |
| 33863894 |  | DASS family sodium/sulfate transporter | Firmicutes | Clostridia | Halanaerobiales | Halanaerobiaceae | Halothermothrix |
| 33863895 |  | Trk family sodium transporter | Firmicutes | Bacilli | Bacillales | Bacillaceae | Bacillus |
| 33863896 |  | VIC family potassium channel protein | Acidobacteria | Holophagae | Holophagales | Holophagaceae | Holophaga |
| 33863897 |  | anhydro-N-acetylmuramic acid kinase | Actinobacteria | Actinobacteria | Actinomycetales | Micrococcaceae | Arthrobacter |
| 33863917 |  | PDZ domain-containing protein | Firmicutes | Clostridia | Thermoanaerobacterales | Thermoanaerobacterales | Caldicellulosiruptor |
| 33863920 |  | septum site-determining protein MinD | Proteobacteria | Gammaproteobacteria | Acidithiobacillales | Acidithiobacillaceae | Acidithiobacillus |
| 33863927 |  | SAM-binding motif-containing protein | Proteobacteria | Alphaproteobacteria | Rhodobacterales | Rhodobacteraceae | Jannaschia |
| 33863938 |  | carbohydrate kinase | Bacteroidetes | Bacteroidia | Bacteroidales | Rikenellaceae | Alistipes |
| 33863955 |  | low molecular weight phosphotyrosine protein phosphatase | Firmicutes | Bacilli | Bacillales | Bacillaceae | Bacillus |
| 33863958 |  | phosphoribosylaminoimidazole synthetase | Firmicutes | Bacilli | Bacillales | Paenibacillaceae | Paenibacillus |
| 33863980 |  | phosphoenolpyruvate carboxylase | Firmicutes | Bacilli | Bacillales | Paenibacillaceae | Paenibacillus |
| 33863982 |  | recombination protein F | Firmicutes | Clostridia | Clostridiales | Clostridiaceae | Clostridium |
| 33863984 |  | hypothetical protein PMT1717 | Bacteroidetes | Sphingobacteriia | Sphingobacteriales | Sphingobacteriaceae | Sphingobacterium |
| 33864000 |  | 50S ribosomal protein L4 | Firmicutes | Bacilli | Bacillales | Bacillaceae | Geobacillus |
| 33864003 |  | 30S ribosomal protein S19 | Firmicutes | Clostridia | Clostridiales | Clostridiaceae | Clostridium |
| 33864008 |  | 30S ribosomal protein S17 | Proteobacteria | Gammaproteobacteria | Xanthomonadales | Xanthomonadaceae | Xanthomonas |
| 33864018 |  | adenylate kinase | Chlorobi | Chlorobia | Chlorobiales | Chlorobiaceae | Chlorobaculum |
| 33864029 |  | HNH endonuclease family protein | Proteobacteria | Betaproteobacteria | Burkholderiales | Comamonadaceae | Alicycliphilus |
| 33864033 |  | glycosyl transferase family protein | Proteobacteria | Gammaproteobacteria | Alteromonadales | Shewanellaceae | Shewanella |
| 33864049 |  | elongation factor Tu | Firmicutes | Clostridia | Clostridiales | Clostridiaceae | Clostridium |
| 33864053 |  | chorismate mutase-prephenate dehydratase | Firmicutes | Clostridia | Clostridiales | Clostridiaceae |  |
| 33864055 |  | ribonuclease HII | Firmicutes | Clostridia | Halanaerobiales | Halobacteroidaceae | Acetohalobium |
| 33864056 |  | ribonuclease E/G | Proteobacteria | Deltaproteobacteria | Myxococcales | Nannocystineae | Nannocystis |
| 33864062 |  | chorismate synthase | Proteobacteria | Betaproteobacteria | Rhodocyclales | Rhodocyclaceae | Azoarcus |
| 33864066 |  | ATP-sulfurylase | Firmicutes | Negativicutes | Selenomonadales | Acidaminococcaceae |  |
| 33864069 |  | p-pantothenate cysteine ligase and p-pantothenenoylcysteine decarboxylase | Proteobacteria | Betaproteobacteria | Rhodocyclales | Rhodocyclaceae | Azoarcus |
| 33864086 |  | hypothetical protein PMT1819 | Proteobacteria | Betaproteobacteria | Rhodocyclales | Rhodocyclaceae | Propionibacter |
| 33864102 |  | peptidyl-tRNA hydrolase | Proteobacteria | Alphaproteobacteria | Rhizobiales | Bradyrhizobiaceae | Bradyrhizobium |
| 33864110 |  | aconitate hydratase, C-terminal, partial | Firmicutes | Clostridia | Clostridiales | Peptococcaceae | Desulfitobacterium |
| 33864112 |  | molybdenum cofactor biosynthesis protein | Proteobacteria | Gammaproteobacteria | Chromatiales | Chromatiaceae | Thiocapsa marina |
| 33864124 |  | bifunctional phosphoribosylaminoimidazolecarboxamide formyltransferase/IMP cyclohydrolase | Firmicutes | Bacilli | Lactobacillales | Carnobacteriaceae | Carnobacterium |
| 33864129 |  | queuine tRNA-ribosyltransferase | Proteobacteria | Gammaproteobacteria | Chromatiales | Ectothiorhodospiraceae | Thioalkalivibrio |
| 33864132 |  | oxidoreductase | Firmicutes | Bacilli | Bacillales | Bacillaceae | Bacillus |
| 33864139 |  | deoxyribonucleotide triphosphate pyrophosphatase | Proteobacteria | Alphaproteobacteria | Rhizobiales | Rhizobiaceae | Rhizobium |
| 33864146 |  | imidazoleglycerol-phosphate dehydratase | Firmicutes | Negativicutes | Selenomonadales | Acidaminococcaceae | Acidaminococcus |
| 33864151 |  | NUDIX hydrolase | Firmicutes | Negativicutes | Selenomonadales | Veillonellaceae | Acetonema |
| 33864155 |  | ABC transporter ATP-binding protein | Firmicutes | Negativicutes | Selenomonadales | Veillonellaceae | Selenomonas |
| 33864156 |  | hypothetical protein PMT1890 | Proteobacteria | Alphaproteobacteria | Rhodospirillales | Acetobacteraceae | Acidiphilium |
| 33864166 |  | 5'-methylthioadenosine phosphorylase | Proteobacteria | Alphaproteobacteria | Rhizobiales | Methylocystaceae | Methylocystis |
| 33864167 |  | methylthioribose-1-phosphate isomerase | Proteobacteria | Betaproteobacteria | Nitrosomonadales | Nitrosomonadaceae | Nitrosomonas |
| 33864169 |  | nucleotide sugar epimerase | Verrucomicrobia | Opitutae | Puniceicoccales | Puniceicoccaceae | Coraliomargarita |
| 33864170 |  | UDP-glucose 6-dehydrogenase | Proteobacteria | Betaproteobacteria | Burkholderiales | Oxalobacteraceae | Janthinobacterium |
| 16135004 |  | histidyl-tRNA synthetase | Planctomycetes | Planctomycetia | Planctomycetales | Planctomycetaceae | rhodospirurella |
| 33864173 |  | UDP-glucose-4-epimerase | Planctomycetes | Planctomycetia | Planctomycetales | Planctomycetaceae | Rhodopirellula |
| 33864192 | X | group 1 glycosyl transferase | Proteobacteria | Epsilonproteobacteria | Nautiliales | Nautiliaceae | Caminibacter |
| 33864195 |  | ABC transporter transmembrane region:ATP/GTP-binding site | Firmicutes | Bacilli | Bacillales | Bacillaceae | Bacillus |
| 33864199 | X | glycosyl transferase family protein | Firmicutes | Bacilli | Bacillales | Paenibacillaceae | Paenibacillus |
| 33864200 |  | isocitrate dehydrogenase | Actinobacteria | Actinobacteria | Actinomycetales | Micromonosporaceae | Actinoplanes |
| 33864204 |  | oxidoreductase | Proteobacteria | Betaproteobacteria | Burkholderiales | Burkholderiaceae | Burkholderia |
| 33864211 |  | phosphorylase | Proteobacteria | Gammaproteobacteria | Chromatiales | Chromatiaceae | Marichromatium |
| 33864218 |  | glucosamine--fructose-6-phosphate aminotransferase | Firmicutes | Bacilli | Bacillales | Bacillaceae | Geobacillus |
| 33864221 |  | 3-oxoacyl-ACP synthase | Proteobacteria | Gammaproteobacteria | Alteromonadales | Alteromonadaceae | Marinobacter |
| 33864223 |  | NAD-dependent epimerase/dehydratase family protein | Proteobacteria | Alphaproteobacteria | Rhizobiales | Rhizobiaceae | Rhizobium |
| 33864224 | X | glycosyl transferase family protein | Proteobacteria | Gammaproteobacteria | Aeromonadales | Aeromonadaceae | Aeromonas |
| 33864238 |  | cell division inhibitor | Chloroflexi | Chloroflexi | Chloroflexales | Chloroflexaceae | Roseiflexus |
| 33862276 |  | phosphoribosylformylglycinamidine synthase II | Actinobacteria | Actinobacteria | Actinomycetales | Corynebacteriaceae | Corynebacterium |
| 33862277 |  | amidophosphoribosyltransferase | Proteobacteria | Gammaproteobacteria | Vibrionales | Vibrionaceae | Aliivibrio |
| 33862280 |  | hypothetical protein PMT0007 | Proteobacteria | Deltaproteobacteria | Desulfovibrionales | Desulfovibrionaceae | Desulfovibrio |
| 33862286 |  | argininosuccinate lyase | Proteobacteria | Betaproteobacteria | Burkholderiales | Burkholderiaceae | Limnobacter |
| 33862289 |  | hypothetical protein PMT0016 | Bacteroidetes | Flavobacteriia | Flavobacteriales | Flavobacteriaceae | Aequorivita |
| 33862291 |  | pili biogenesis protein | Proteobacteria | Deltaproteobacteria | Myxococcales | Polyangiaceae | Sorangium |
| 33862292 |  | PilT1 protein | Synergistetes | Synergistia | Synergistales | Synergistaceae | Aminomonas |
| 33862293 |  | general secretion pathway protein E | Firmicutes | Bacilli | Bacillales | Alicyclobacillaceae | Alicyclobacillus |
| 33862295 |  | chaperone protein DnaJ | Firmicutes | Clostridia | Clostridiales | Lachnospiraceae | Coprococcus |
| 33862299 |  | UDP-N-acetylenolpyruvoylglucosamine reductase | Firmicutes | Negativicutes | Selenomonadales | Veillonellaceae | Selenomonas |
| 33862300 |  | UDP-N-acetylmuramate--L-alanine ligase | Proteobacteria | Deltaproteobacteria | Myxococcales | Polyangiaceae | Sorangium |
| 33862301 |  | glyceraldehyde 3-phosphate dehydrogenase | Firmicutes | Bacilli | Lactobacillales | Carnobacteriaceae | Carnobacterium |
| 33862304 |  | elongation factor P | Proteobacteria | Gammaproteobacteria | Enterobacteriales | Enterobacteriaceae | Escherichia |
| 33862306 |  | 4-hydroxythreonine-4-phosphate dehydrogenase | Firmicutes | Bacilli | Bacillales | Alicyclobacillaceae | Alicyclobacillus |
| 33862323 |  | sulfolipid (UDP-sulfoquinovose) biosynthesis protein | Firmicutes | Bacilli | Bacillales | Bacillaceae | Anoxybacillus |
| 33862325 |  | thiazole synthase | Firmicutes | Bacilli | Bacillales | Paenibacillaceae | Paenibacillus |
| 33862336 |  | trigger factor | Firmicutes | Negativicutes | Selenomonadales | Veillonellaceae | Pelosinus |
| 33862338 |  | dihydrodipicolinate synthase | Firmicutes | Clostridia | Clostridiales | Clostridiaceae | Clostridium |
| 33862339 |  | hypothetical protein PMT0066 | Firmicutes | Bacilli | Bacillales | Paenibacillaceae | Brevibacillus |
| 33862344 |  | excinuclease ABC subunit B | Proteobacteria | Gammaproteobacteria | Acidithiobacillales | Acidithiobacillaceae | Acidithiobacillus |
| 33862350 |  | multidrug ABC transporter | Firmicutes | Clostridia | Clostridiales | Lachnospiraceae | Butyrivibrio |
| 33862352 |  | DNA mismatch repair protein MutS | Firmicutes | Bacilli | Bacillales | Bacillaceae | Anoxybacillus |
| 33862354 |  | 6,7-dimethyl-8-ribityllumazine synthase | Proteobacteria | Deltaproteobacteria | Desulfovibrionales | Desulfovibrionaceae | Desulfovibrio |
| 33862359 |  | UDP-glucose-4-epimerase | Chlorobi | Chlorobia | Chlorobiales | Chlorobiaceae | Pelodictyon |
| 33862366 |  | multidrug efflux family ABC transporter | Proteobacteria | Gammaproteobacteria | Methylococcales | Methylococcaceae | Methylomonas |
| 33862368 |  | capsular polysaccharide biosynthesis protein | Proteobacteria | Deltaproteobacteria | Desulfovibrionales | Desulfovibrionaceae | Desulfovibrio |
| 33862372 |  | imidazoleglycerol-phosphate synthase, glutamine amidotransferase subunit | Proteobacteria | Deltaproteobacteria | Desulfobacterales | Desulfobacteraceae | Desulfatibacillum |
| 33862373 |  | imidazole glycerol phosphate synthase subunit HisF | Proteobacteria | Alphaproteobacteria | Rhizobiales | Beijerinckiaceae | Beijerinckia |
| 33862381 |  | ABC transporter ATP-binding protein | Aquificae | Aquificae | Aquificales | Desulfurobacteriaceae | Desulfurobacterium |
| 33862386 |  | dTDP-4-dehydrorhamnose 3,5-epimerase | Proteobacteria | Gammaproteobacteria | Chromatiales | Ectothiorhodospiraceae | Nitrococcus |
| 33862388 |  | dTDP-glucose-4,6-dehydratase | Firmicutes | Negativicutes | Selenomonadales | Veillonellaceae | Acetonema |
| 33862390 |  | Serine acetyltransferase | Firmicutes | Bacilli | Bacillales | Bacillaceae | Bacillus |
| 33862393 |  | tRNA delta(2)-isopentenylpyrophosphate transferase | Proteobacteria | Alphaproteobacteria | Sphingomonadales | Sphingomonadaceae | Sphingomonas |
| 33862402 |  | RND family multidrug efflux transporter | Bacteroidetes | Sphingobacteriia | Sphingobacteriales | Saprospiraceae | Saprospira |
| 33862405 |  | succinate dehydrogenase flavoprotein subunit | Proteobacteria | Gammaproteobacteria | Oceanospirillales | Alcanivoracaceae | Kangiella |
| 33862411 |  | S-adenosyl-L-homocysteine hydrolase | Firmicutes | Bacilli | Bacillales | Paenibacillaceae | Paenibacillus |
| 33862418 |  | substrate-binding family 1 protein | Firmicutes | Clostridia | Clostridiales | Ruminococcaceae | Acetivibrio |
| 33862419 |  | two-component response regulator | Firmicutes | Bacilli | Lactobacillales | Carnobacteriaceae | Carnobacterium |
| 33862428 |  | SsrA-binding protein | Proteobacteria | Gammaproteobacteria | Alteromonadales | Alteromonadaceae | Alishewanella |
| 33862429 |  | Holliday junction DNA helicase RuvB | Chloroflexi | Chloroflexi | Chloroflexales | Chloroflexaceae | Roseiflexus |
| 33862433 |  | thiamine biosynthesis protein ThiC | Proteobacteria | Gammaproteobacteria | Pseudomonadales | Moraxellaceae | Acinetobacter |
| 33862436 |  | shikimate kinase | Firmicutes | Bacilli | Bacillales | Paenibacillaceae | Paenibacillus |
| 33862450 |  | L-aspartate oxidase | Proteobacteria | Alphaproteobacteria | Rhizobiales | Rhizobiaceae | Agrobacterium |
| 33862453 |  | UDP pyrophosphate phosphatase | Firmicutes | Clostridia | Clostridiales | Clostridiaceae | Alkaliphilus |
| 33862454 |  | Fe-S oxidoreductase | Chlorobi | Chlorobia | Chlorobiales | Chlorobiaceae | Chlorobaculum |
| 33862464 |  | aldehyde dehydrogenase | Proteobacteria | Alphaproteobacteria | Rhizobiales | Phyllobacteriaceae | Parvibaculum |
| 33862466 |  | sodium/alanine symporter family protein | Proteobacteria | Zetaproteobacteria | Mariprofundales | Mariprofundaceae | Mariprofundus |
| 33862468 |  | 3-dehydroquinate dehydratase | Chloroflexi | Chloroflexia | Chloroflexales | Chloroflexaceae | Chloroflexus |
| 33862476 |  | GTP-binding protein EngA | Firmicutes | Negativicutes | Selenomonadales | Veillonellaceae | selenomonas noxia |
| 33862481 |  | Delta 1-pyrroline-5-carboxylate reductase | Proteobacteria | Alphaproteobacteria | Rhodospirillales | Rhodospirillaceae | Azospirillum |
| 33862485 |  | deoxyribose-phosphate aldolase | Chlamydiae | Chlamydiia | Chlamydiales | Parachlamydiaceae | Parachlamydia |
| 33862499 | X | cystathionine gamma-synthase | Proteobacteria | Gammaproteobacteria | Pseudomonadales | Pseudomonadaceae | Pseudomonas |
| 33862503 |  | UDP-N-acetylmuramoylalanyl-D-glutamate--2,6-diaminopimelate ligase | Actinobacteria | Actinobacteria | Coriobacteriales | Coriobacteriaceae | Eggerthella |
| 33862538 |  | two component sensor histidine kinase | Actinobacteria | Actinobacteria | Actinomycetales | Mycobacteriaceae | Mycobacterium |
| 33862539 |  | peptide ABC transporter | Firmicutes | Bacilli | Bacillales | Paenibacillaceae | Paenibacillus |
| 33862543 |  | Sun protein (Fmu protein) | Chloroflexi | Chloroflexia | Chloroflexales | Roseiflexaceae | Roseiflexus |
| 33862545 |  | bacteriochlorophyll/chlorophyll a synthase | Firmicutes | Clostridia | Clostridiales | Clostridiaceae | Clostridium |
| 33862547 |  | imidazole glycerol phosphate synthase subunit HisF | Proteobacteria | Deltaproteobacteria | Desulfuromonadales | Geobacteraceae | Geobacter |
| 33862556 |  | glycyl-tRNA synthetase subunit alpha | Proteobacteria | Gammaproteobacteria | Oceanospirillales | Halomonadaceae | Halomonas |
| 33862559 |  | hydroxylase | Proteobacteria | Gammaproteobacteria | Vibrionales | Vibrionaceae | Photobacterium |
| 33862805 |  | phosphoribulokinase | Proteobacteria | Gammaproteobacteria |  | Sedimenticola |  |
| 33863245 |  | hypothetical protein PMT0974 | Proteobacteria | Gammaproteobacteria | Thiotrichales | Thiotrichaceae | Beggiatoa |
| 33863248 |  | multidrug ABC transporter | Proteobacteria | Gammaproteobacteria | Legionellales | Legionellaceae | Legionella |
| 33863249 |  | multidrug ABC transporter | Actinobacteria | Actinobacteria | Actinomycetales | Corynebacteriaceae | Corynebacterium |
| 33863265 |  | two-component response regulator, phosphate | Proteobacteria | Betaproteobacteria | Rhodocyclales | Rhodocyclaceae | Thauera |
| 33863271 |  | glyceraldehyde 3-phosphate dehydrogenase | Proteobacteria | Gammaproteobacteria | Chromatiales | Chromatiaceae | Allochromatium |
| 33863303 |  | hypothetical protein PMT1032 | Firmicutes | Clostridia | Clostridiales | Peptococcaceae | Desulfotomaculum |
| 33863320 |  | tRNA (uracil-5-)-methyltransferase Gid | Proteobacteria | Gammaproteobacteria | Chromatiales | Ectothiorhodospiraceae | Thioalkalivibrio |
| 33863323 |  | aminopeptidase N | Proteobacteria | Betaproteobacteria | Burkholderiales | Burkholderiaceae | Limnobacter |
| 33863326 |  | hypothetical protein PMT1055 | Firmicutes | Clostridia | Halanaerobiales | Halobacteroidaceae | Acetohalobium |
| 33863328 |  | UDP pyrophosphate synthase | Firmicutes | Clostridia | Clostridiales | Clostridiaceae | Clostridium |
| 33863340 |  | pseudouridylate synthase specific to ribosomal small subunit | Proteobacteria | Zetaproteobacteria | Mariprofundales | Mariprofundaceae | Mariprofundus |
| 33863342 |  | 4-alpha-glucanotransferase | Proteobacteria | Betaproteobacteria | Nitrosomonadales | Nitrosomonadaceae | Nitrosomonas |
| 33863351 |  | multidrug ABC transporter | Planctomycetes | Planctomycetia | Planctomycetales | Planctomycetaceae | Planctomyces |
| 33863354 |  | ATP-dependent RNA helicase | Nitrospirae | Nitrospira | Nitrospirales | Nitrospiraceae | Thermodesulfovibrio |
| 33863362 |  | pyridoxine 5'-phosphate synthase | Proteobacteria | Gammaproteobacteria | Enterobacteriales | Enterobacteriaceae | Escherichia |
| 33863372 |  | oxidoreductase | Actinobacteria | Actinobacteria | Actinomycetales | Streptomycetaceae | Streptomyces |
| 33863373 |  | glucose-6-phosphate 1-dehydrogenase | Firmicutes | Bacilli | Bacillales | Paenibacillaceae | Brevibacillus |
| 33863375 |  | cobyrinic acid a,c-diamide synthase | Firmicutes | Clostridia | Clostridiales | Clostridiaceae | Clostridium |
| 33863381 |  | bifunctional 5,10-methylene-tetrahydrofolate dehydrogenase/ 5,10-methylene-tetrahydrofolate cyclohydrolase | Proteobacteria | Gammaproteobacteria | Alteromonadales | Alteromonadaceae | Alishewanella |
| 33863383 |  | pseudouridylate synthase | Proteobacteria | Gammaproteobacteria | Xanthomonadales | Xanthomonadaceae | Xanthomonas |
| 33863391 |  | binding-protein dependent transport system inner membrane protein | Proteobacteria | Gammaproteobacteria | Methylococcales | Methylococcaceae | Methylomicrobium |
| 33863393 |  | hypothetical protein PMT1122 -1,4-alpha-glucan branching enzyme | Actinobacteria | Actinobacteria | Actinomycetales | Microbacteriaceae | Clavibacter |
| 33863397 |  | inosine 5-monophosphate dehydrogenase | Firmicutes | Clostridia | Clostridiales | Clostridiaceae | Alkaliphillus |
| 33863399 |  | imidazole glycerol phosphate synthase subunit HisH | Firmicutes | Bacilli | Bacillales | Paenibacillaceae | Paenibacillus |
| 33863407 |  | Holliday junction resolvase | Thermotogae | Thermotogae | Thermotogales | Thermotogaceae | Thermotoga |
| 33863425 |  | GTP-dependent nucleic acid-binding protein EngD | Proteobacteria | Betaproteobacteria | Burkholderiales | Comamonadaceae | Comamonas |
| 33863440 |  | pseudouridine synthase | Firmicutes | Negativicutes | Selenomonadales | Veillonellaceae | Dialister |
| 33863453 |  | cobyric acid synthase | Proteobacteria | Gammaproteobacteria | Oceanospirillales | Alcanivoracaceae | Alcanivorax |
| 33863457 |  | chromosomal replication initiation protein | Proteobacteria | Alphaproteobacteria | Rhizobiales | Bradyrhizobiaceae | Bradyrhizobium |
| 33863460 |  | hypothetical protein PMT1190 | Chlorobi | Chlorobia | Chlorobiales | Chlorobiaceae | Chlorobaculum |
| 33863462 |  | multidrug efflux ABC transporter | Proteobacteria | Alphaproteobacteria | Rhodospirillales | Acetobacteraceae | Acetobacter |
| 33863464 |  | hydroxyacylglutathione hydrolase | Firmicutes | Bacilli | Bacillales | Bacillaceae | Bacillus |
| 33863465 |  | hypothetical protein PMT1195 | Proteobacteria | Gammaproteobacteria | Chromatiales | Ectothiorhodospiraceae | Thioalkalivibrio |
| 33863469 |  | carboxysome structural protein CsoS1 | Proteobacteria | Gammaproteobacteria | Chromatiales | Chromatiaceae | Thioflavicoccus |
| 33863475 |  | ribulose bisophosphate carboxylase | Proteobacteria | Gammaproteobacteria | Chromatiales | Ectothiorhodospiraceae | Thioalkalivibrio |
| 33863476 |  | carboxysome shell protein CsoS1 | Proteobacteria | Gammaproteobacteria | Alteromonadales | Pseudoalteromonadaceae | Pseudoalteromonas |
| 33863477 |  | HAM1 family protein | Proteobacteria | Gammaproteobacteria | Chromatiales | Chromatiaceae | Allochromatium |
| 33863478 |  | hypothetical protein PMT1208 | Proteobacteria | Gammaproteobacteria | Chromatiales | Chromatiaceae | Thiocapsa |
| 33863484 |  | light-independent protochlorophyllide reductase subunit N | Proteobacteria | Betaproteobacteria | Rhodocyclales | Rhodocyclaceae | methyloversatilis |
| 33863485 |  | light-independent protochlorophyllide reductase subunit B | Proteobacteria | Betaproteobacteria | Rhodocyclales | Rhodocyclaceae | sulfitobacter |
| 33863486 |  | protochlorophyllide reductase iron-sulfur ATP-binding protein | Proteobacteria | Alphaproteobacteria | Rhodobacterales | Rhodobacteraceae | Rhodobacter |
| 33863487 |  | protochlorophyllide oxidoreductase | Proteobacteria | Gammaproteobacteria | Alteromonadales | Alteromonadaceae | Alteromonas |
| 33863495 |  | GTP cyclohydrolase I | Proteobacteria | Betaproteobacteria | Burkholderiales | Comamonadaceae | Acidovorax |
| 33863497 |  | acetyl-CoA carboxylase carboxyltransferase subunit alpha | Actinobacteria | Actinobacteria | Actinomycetales | Gordoniaceae | Gordonia |
| 33863500 |  | creatininase | Euryarchaeota | Methanomicrobia | Methanomicrobiales | Methanoregulaceae | Methanoregula |
| 33863507 |  | acetolactate synthase 3 catalytic subunit | Aquificae | Aquificae | Aquificales | Aquificaceae | Hydrogenobaculum |
| 33863508 |  | ferrochelatase | Proteobacteria | Gammaproteobacteria | Pseudomonadales | Pseudomonadaceae | Pseudomonas |
| 33863511 |  | cob(I)alamin adenosyltransferase | Actinobacteria | Actinobacteria | Coriobacteriales | Coriobacteriaceae | Eggerthella |
| 33863512 |  | uridylate kinase | Proteobacteria | Gammaproteobacteria | Pasteurellales | Pasteurellaceae | Pasteurella |
| 33863516 |  | transaldolase/EF-hand domain-containing protein | Firmicutes | Bacilli | Bacillales | Bacillaceae | Bacillus |
| 33863525 |  | inorganic pyrophosphatase | Deinococcus | Deinococci | Deinococcales | Deinococcaceae | Deinococcus |
| 33863527 |  | prolyl-tRNA synthetase | Firmicutes | Clostridia | Halanaerobiales | Halobacteroidaceae | Acetohalobium |
| 33863536 |  | acetylglutamate kinase | Firmicutes | Bacilli | Lactobacillales | Lactobacillaceae | Lactobacillus |
| 33863539 |  | primosomal protein N' | Proteobacteria | Gammaproteobacteria | Cardiobacteriales | Cardiobacteriaceae | Dichelobacter |
| 33863543 |  | inorganic pyrophosphatase | Proteobacteria | Alphaproteobacteria | 0 | 0 | pelagibacter |
| 33863552 |  | hypothetical protein PMT1284 | Proteobacteria | Gammaproteobacteria | Pseudomonadales | Pseudomonadaceae | Azotobacter |
| 33863555 |  | iron ABC transporter | Proteobacteria | Deltaproteobacteria | Desulfobacterales | Desulfobulbaceae | Desulfobulbus |
| 33863559 |  | hypothetical protein PMT1291 | Actinobacteria | Actinobacteria | Actinomycetales | Nocardiopsaceae | Nocardiopsis |
| 33863563 |  | glycolate oxidase subunit glcD | Actinobacteria | Actinobacteria | Actinomycetales | Actinomycetaceae | Actinomyces |
| 16135004 |  | nucleotide-binding protein | Firmicutes | Bacilli | Bacillales | Bacillaceae | Bacillus |
| 33864255 |  | thymidylate kinase | Proteobacteria | Deltaproteobacteria | Desulfobacterales | Desulfobacteraceae | Desulfatibacillum |
| 33864256 |  | P-type ATPase transporter for copper | Proteobacteria | Alphaproteobacteria | Rhodobacterales | Rhodobacteraceae | Rhodobacter |
| 33864258 |  | DNA repair protein RadA | Actinobacteria | Actinobacteria | Actinomycetales | Actinomycetaceae | Actinomyces |
| 33864259 |  | two-component response regulator | Spirochaetes | Spirochaetia | Spirochaetales | Leptospiraceae | Leptospira |
| 33864260 |  | glycerol-3-phosphate acyltransferase PlsX | Firmicutes | Clostridia | Thermoanaerobacterales | Thermoanaerobacterales | Caldicellulosiruptor |
| 33864261 |  | 3-oxoacyl-ACP synthase | Firmicutes | Clostridia | Thermoanaerobacterales | Thermoanaerobacterales | Caldicellulosiruptor |
| 33864272 |  | Rubisco transcriptional regulator | Firmicutes | Clostridia | Clostridiales | Lachnospiraceae | Blautia |
| 33864278 |  | methylenetetrahydrofolate reductase | Firmicutes | Bacilli | Bacillales | Listeriaceae | Listeria |
| 33864291 |  | negative regulator of class I heat shock protein | Firmicutes | Bacilli | Lactobacillales | Enterococcaceae | Enterococcus |
| 33864303 |  | phosphoribosylaminoimidazole carboxylase | Proteobacteria | Alphaproteobacteria | Rhodobacterales | Rhodobacteraceae | Ahrensia |
| 33864305 |  | Mg-protoporphyrin IX methyl transferase | Planctomycetes | Planctomycetia | Planctomycetales | Planctomycetaceae | Blastopirellula |
| 33864313 |  | S-adenosyl-methyltransferase MraW | Proteobacteria | Alphaproteobacteria | Rhodobacterales | Rhodobacteraceae | Rhodobacter |
| 33864314 |  | NAD(P)H-quinone oxidoreductase subunit H | Firmicutes | Bacilli | Bacillales | Bacillaceae | Oceanobacillus |
| 33864325 |  | isochorismate synthase | Proteobacteria | Gammaproteobacteria | Acidithiobacillales | Acidithiobacillaceae | Acidithiobacillus |
| 33864326 |  | glutathione synthetase | Proteobacteria | Betaproteobacteria | Neisseriales | Neisseriaceae | Neisseria |
| 33864335 |  | arginyl-tRNA synthetase | Firmicutes | Bacilli | Bacillales | Paenibacillaceae | Paenibacillus |
| 33864337 |  | nicotinate-nucleotide pyrophosphorylase:quinolinate phosphoriobsyl transferase | Firmicutes | Bacilli | Bacillales | Paenibacillaceae | Paenibacillus |
| 33864338 |  | tRNA modification GTPase TrmE | Chloroflexi | Chloroflexi | Chloroflexales | Chloroflexaceae | Roseiflexus |
| 33864348 |  | glyoxalase/bleomycin resistance protein/dioxygenase family protein | Firmicutes | Bacilli | Bacillales | Alicyclobacillaceae | Alicyclobacillus |
| 33864353 |  | 50S ribosomal protein L1 | Bacteroidetes | Flavobacteriia | Flavobacteriales | Flavobacteriaceae | Myroides |
| 33864358 |  | dihydroorotate dehydrogenase 2 | Firmicutes | Bacilli | Lactobacillales | Streptococcaceae | Streptococcus |
| 33864364 |  | class-I aminotransferase | Firmicutes | Bacilli | Bacillales | Bacillaceae | Bacillus |
| 33864365 |  | UDP-PP-MurNAc-pentapeptide-UDPGlcNAc GlcNAc transferase | Proteobacteria | Gammaproteobacteria | Chromatiales | Chromatiaceae | Thiocapsa |
| 33864367 |  | hypothetical protein PMT2103 | Proteobacteria | Deltaproteobacteria | Syntrophobacterales | Syntrophaceae | Desulfomonile |
| 33864369 |  | 3-hydroxyisobutyrate dehydrogenase | Proteobacteria | Alphaproteobacteria | Rhodobacterales | Rhodobacteraceae | Celeribacter |
| 33864374 |  | ABC transporter ATP-binding protein | Firmicutes | Clostridia | Halanaerobiales | Halanaerobiaceae | Halanaerobium |
| 33864400 |  | dephospho-CoA kinase | Proteobacteria | Alphaproteobacteria | Rhodospirillales | Acetobacteraceae | Acidiphilium |
| 33864404 |  | thiamine biosynthesis oxidoreductase | Proteobacteria | Betaproteobacteria | Rhodocyclales | Rhodocyclaceae | Azoarcus |
| 33863248 |  | multidrug ABC transporter | Proteobacteria | Deltaproteobacteria | Myxococcales | Myxococcaceae | Anaeromyxobacter |
| 33864414 |  | arginine decarboxylase | Proteobacteria | Deltaproteobacteria | Desulfovibrionales | Desulfovibrionaceae | Desulfovibrio |
| 33864416 |  | guanylate cyclase | Fusobacteria | Fusobacteriia | Fusobacteriales | Fusobacteriaceae | Fusobacterium |
| 33864421 |  | alanyl-tRNA synthetase | Proteobacteria | Gammaproteobacteria | Alteromonadales | Alteromonadaceae | Alteromonas |
| 33864433 |  | glycine dehydrogenase | Firmicutes | Clostridia | Clostridiales | Peptococcaceae | Desulfotomaculum |
| 33864434 |  | glycine cleavage system protein H | Firmicutes | Bacilli | Bacillales | Bacillaceae | Bacillus |
| 33864452 |  | multidrug efflux membrane protein | Planctomycetes | Phycisphaerae | Phycisphaerales | Phycisphaeraceae | Phycisphaera |
| 33864460 |  | magnesium-protoporphyrin IX monomethyl ester cyclase | Proteobacteria | Alphaproteobacteria | Rhizobiales | Bradyrhizobiaceae | Bradyrhizobium |
| 33864475 |  | nucleoside triphosphate pyrophosphohydrolase | Firmicutes | Clostridia | Clostridiales | Peptococcaceae | Desulfotomaculum |
| 33864480 |  | aspartyl-tRNA synthetase | Deinococcus | Deinococci | Thermales | Thermaceae | Marinithermus |
| 33864484 |  | organic radical activating protein | Proteobacteria | Betaproteobacteria | Burkholderiales | Alcaligenaceae | Achromobacter |
| 33864489 |  | ATP-binding subunit of urea ABC transport system | Proteobacteria | Gammaproteobacteria | Pseudomonadales | Pseudomonadaceae | Pseudomonas |
| 33864490 |  | ATP-binding subunit of urea ABC transport system | Proteobacteria | Epsilonproteobacteria | Campylobacterales | Campylobacteraceae | Arcobacter |
| 33864492 |  | urea ABC transporter | Proteobacteria | Alphaproteobacteria | Rhodospirillales | Rhodospirillaceae | Azospirillum |
| 33864498 |  | urease subunit gamma | Proteobacteria | Alphaproteobacteria | Rhodobacterales | Rhodobacteraceae | Roseobacter |
| 33864499 |  | urease subunit beta | Firmicutes | Negativicutes | Selenomonadales | Veillonellaceae | Thermosinus |
| 33864503 |  | ferredoxin-nitrite reductase | Actinobacteria | Actinobacteria | Actinomycetales | Mycobacteriaceae | Mycobacterium |
| 33864509 |  | polyphosphate kinase | Actinobacteria | Actinobacteria | Rubrobacterales | Rubrobacteraceae | Rubrobacter |
| 33864512 |  | phospho-2-dehydro-3-deoxyheptonate aldolase | Synergistetes | Synergistia | Synergistales | Synergistaceae | Anaerobaculum |
| 33864528 |  | phospho-N-acetylmuramoyl-pentapeptide-transferase | Proteobacteria | Gammaproteobacteria | Oceanospirillales | Alcanivoracaceae | Alcanivorax |
| 33864531 |  | phosphoribosylglycinamide formyltransferase 2 | Actinobacteria | Actinobacteria | Actinomycetales | Actinomycetaceae | Actinomyces |
