## Supplemental for "Whole genome phylogeny of Cyanobacteria documents a distinct evolutionary trajectory of marine picocyanobacteria": Supplementary_Table_S2 .docx

| Supplementary_Table_S2 – List of prokaryotic genomes used in this study  Accession numbers, genome lengths and taxonomy for all genomes is presented. | | | |
| --- | --- | --- | --- |
| Strain | Genbank genome accession number | Genome length [nt] | Taxonomic lineage |
| *Anabaena variabilis* ATCC 29413 | CP000117.1 | 6356727 | Cyanobacteria; Nostocales; Nostocaceae; Anabaena |
| *Arthrospira maxima* CS-328 | NZ_ABYK01000000 | 6003314 | Cyanobacteria; Oscillatoriophycideae; Oscillatoriales; |
| *Arthrospira platensis* NIES-39 | NC_016640 | 6788435 | Cyanobacteria; Oscillatoriophycideae; Oscillatoriales; Arthrospira |
| *Coleofasciculus chthonoplastes* PCC 7420 | NZ_ABRS01000000 | 8651623 | Cyanobacteria; Oscillatoriophycideae; Oscillatoriales; |
| *Crinalium epipsammum* PCC 9333 | NC_019753 | 5315554 | Cyanobacteria; Oscillatoriophycideae; Oscillatoriales; Crinalium |
| *Cyanobacterium* UCYNA | NC_013771 | 1443806 | Cyanobacteria; Oscillatoriophycideae; Chroococcales; Cand. Atelocyanobacterium |
| *Cyanobium gracile* PCC 6307 | NC_019675 | 3342364 | Cyanobacteria; Oscillatoriophycideae; Chroococcales; Cyanobium |
| *Cyanobium* sp. PCC 7001 | NZ_ABSE00000000 | 2834250 | Cyanobacteria; Oscillatoriophycideae; Chroococcales |
| *Cyanothece* sp. ATCC 51142 | CP000806 | 4934271 | Cyanobacteria; Nostocales; Nostocaceae |
| *Cyanothece* sp. ATCC 51142 | CP000806 | 4934271 | Cyanobacteria; Oscillatoriophycideae; Chroococcales |
| *Cyanothece* sp. PCC 7424 | NC_011729 | 5942652 | Cyanobacteria; Oscillatoriophycideae; Chroococcales; Cyanothece |
| *Cylindrospermopsis raciborskii* CS-505 | NZ_ACYA00000000 | 3742440 | Cyanobacteria; Nostocales; Nostocaceae |
| *Fischerella* sp. JSC-11 | NZ_AP017305 | 5821603 | Cyanobacteria; Nostocales; Nostocaceae |
| *Geitlerinema* sp. PCC 7407 | CP003591 | 4681111 | Cyanobacteria; Oscillatoriophycideae; Oscillatoriales; Geitlerinema |
| *Gloeobacter violaceus* PCC 7421 | NC_005125 | 4659019 | Cyanobacteria; Gloeobacteria; Gloeobacterales; Gloeobacter |
| *Gloeocapsa* sp. PCC 7428 | CP003646 | 5431448 | Cyanobacteria; Oscillatoriophycideae; Chroococcales; Gloeocapsa |
| *Halothece* sp. PCC 7418 | CP003945 | 4179170 | Cyanobacteria; Oscillatoriophycideae; Chroococcales; Halothece cluster; Halothece |
| *Leptolyngbya* sp. PCC 7376 | NC_019683 | 5125950 | Cyanobacteria; Oscillatoriophycideae; Oscillatoriales; Leptolyngbya |
| *Lyngbya* sp. PCC 8106 | NZ_AAVU01000000 | 7037511 | Cyanobacteria; Oscillatoriophycideae; Oscillatoriales; |
| *Microcoleus* sp. PCC 7113 | NC_019738 | 7470429 | Cyanobacteria; Oscillatoriophycideae; Oscillatoriales; Microcoleus |
| *Microcoleus vaginatus* FGP-2 | NZ_AFJC01000000 | 6698929 | Cyanobacteria; Oscillatoriophycideae; Oscillatoriales; |
| *Microcystis aeruginosa* NIES-843 DNA | NC_010296 | 5842795 | Cyanobacteria; Oscillatoriophycideae; Chroococcales; Microcystis |
| *Lyngbya majuscula* 3L | NZ_AEPQ01000000 | 8389417 | Cyanobacteria; Oscillatoriophycideae; Oscillatoriales; |
| *Nodularia spumigena* CCY9414 | NZ_CP007203 | 5462271 | Cyanobacteria; Nostocales; Nostocaceae |
| *Nostoc punctiforme* PCC 73102 | NC_010628 | 8234322 | Cyanobacteria; Nostocales; Nostocaceae; Nostoc |
| *Nostoc* sp. PCC 7120 | NC_003272 | 6413771 | Cyanobacteria; Nostocales; Nostocaceae; Nostoc |
| *Oscillatoria acuminata* PCC 6304 | CP003607 | 7689443 | Cyanobacteria; Oscillatoriophycideae; Oscillatoriales; Oscillatoria |
| *Oscillatoria nigro-viridis* PCC 7112 | CP003614 | 7479014 | Cyanobacteria; Oscillatoriophycideae; Oscillatoriales; Oscillatoria |
| *Prochlorococcus marinus* MIT 9313 | NC_005071 | 2410873 | Cyanobacteria; Prochlorales; Prochlorococcaceae; Prochlorococcus |
| *Prochlorococcus marinus* str. MIT 9215 | NC_009840 | 1738790 | Cyanobacteria; Prochlorales; Prochlorococcaceae; Prochlorococcus |
| *Prochlorococcus marinus* str. MIT 9301 | NC_009091 | 1641879 | Cyanobacteria; Prochlorales; Prochlorococcaceae; Prochlorococcus. |
| *Prochlorococcus marinus* str. MIT 9303 | NC_008820 | 2682675 | Cyanobacteria; Prochlorales; Prochlorococcaceae; Prochlorococcus. |
| *Prochlorococcus marinus* str. MIT 9312 | NC_007577 | 1709204 | Cyanobacteria; Prochlorales; Prochlorococcaceae; Prochlorococcus |
| Prochlorococcus marinus str. NATL2A | NC_007335 | 1842899 | Cyanobacteria; Prochlorales; Prochlorococcaceae; Prochlorococcus. |
| *Raphidiopsis brookii* D9 | NZ_ACYB00000000 | 3186511 | Cyanobacteria; Nostocales; Nostocaceae |
| *Rivularia sp.* PCC 7116 | NC_019678 | 8698463 | Cyanobacteria; Nostocales; Rivulariaceae; Rivularia |
| *Synechococcus* PCC 7335 | NZ_ABRV000000000 | 5972042 | Cyanobacteria; Oscillatoriophycideae; Chroococcales |
| *Synechococcus* sp. CB0101 | NZ_ADXL00000000 | 2686395 | Cyanobacteria; Oscillatoriophycideae; Chroococcales |
| *Synechococcus* sp. CC9902 | CP000097 | 2234828 | Cyanobacteria; Oscillatoriophycideae; Chroococcales; Synechococcus. |
| *Synechococcus* sp. JA-2-3B'a | CP000240 | 3046682 | Cyanobacteria; Oscillatoriophycideae; Chroococcales |
| *Synechococcus* sp. JA-3-3Ab | CP000239 | 2932766 | Cyanobacteria; Oscillatoriophycideae; Chroococcales |
| *Synechococcus* sp. PCC 7002 | CP000951 | 3008047 | Cyanobacteria; Oscillatoriophycideae; Chroococcales; Synechococcus. |
| *Synechococcus* sp. PCC 7502 | CP003594 | 3510253 | Cyanobacteria; Oscillatoriophycideae; Chroococcales; Synechococcus. |
| *Synechococcus* sp. RCC307 | CT978603 | 2794318 | Cyanobacteria; Oscillatoriophycideae; Chroococcales; Synechococcus. |
| *Synechococcus* sp. WH 7803 | CT971583 | 2366980 | Cyanobacteria; Oscillatoriophycideae; Chroococcales; Synechococcus. |
| *Synechococcus* sp. WH 8102 | BX548020 | 2434428 | Cyanobacteria; Oscillatoriophycideae; Chroococcales; Synechococcus. |
| *Thermosynechococcus elongatus* BP-1 | NC_004113 | 2593857 | Cyanobacteria; Oscillatoriophycideae; Chroococcales; Thermosynechococcus. |
| *Trichodesmium erythraeum* IMS101 | CP000393 | 7750108 | Cyanobacteria; Oscillatoriophycideae; Oscillatoriales; Trichodesmium. |
| *Thermus thermophilus* HB27 | NC_005835 | 1894877 | Deinococcus-Thermus; Deinococci; Thermales; Thermaceae; Thermus |
| *Dictyoglomus thermophilum* H-6-12 | NC_011297 | 1959987 | Dictyoglomi; Dictyoglomales; Dictyoglomaceae; Dictyoglomus |
| *Clostridium botulinum* A3 str. Loch Maree | NC_010520 | 3992906 | Firmicutes; Clostridia; Clostridiales; Clostridiaceae; Clostridium |
| *Heliobacterium modesticaldum* Ice1 | NC_010337.2 | 3075407 | Firmicutes; Clostridia; Clostridiales; Heliobacteriaceae; Heliobacterium |
| *Fusobacterium nucleatum* subsp. *nucleatum* ATCC 25586 | NC_003454 | 2174500 | Fusobacteria; Fusobacteriales; Fusobacteriaceae; Fusobacterium |
| *Rhodopirellula baltica* SH 1 | NC_005027 | 7145576 | Planctomycetes; Planctomycetia; Planctomycetales; Planctomycetaceae; Rhodopirellula |
| *Candidatus Pelagibacter* sp. IMCC9063 | NC_015380 | 1284727 | Proteobacteria; Alphaproteobacteria; Pelagibacterales; Pelagibacteraceae; Candidatus Pelagibacter |
| *Candidatus Pelagibacter ubique* HTCC1062 | NC_007205 | 1308759 | Proteobacteria; Alphaproteobacteria; Pelagibacterales; Pelagibacteraceae; Candidatus Pelagibacter |
| *Pelagibacterium halotolerans* B2 | NC_016078 | 3944837 | Proteobacteria; Alphaproteobacteria; Rhizobiales; Hyphomicrobiaceae; Pelagibacterium |
| *Rhizobium tropici* CIAT 899 | NC_020059 | 3837060 | Proteobacteria; Alphaproteobacteria; Rhizobiales; Rhizobiaceae; Rhizobium/Agrobacterium group; Rhizobium |
| *Roseobacter denitrificans* OCh 114 | NC_008209 | 4133097 | Proteobacteria; Alphaproteobacteria; Rhodobacterales; Rhodobacteraceae; Roseobacter. |
| *Burkholderia pseudomallei* K96243 | NC_006350 | 4074542 | Proteobacteria; Betaproteobacteria; Burkholderiales; Burkholderiaceae; Burkholderia |
| *Rubrivivax gelatinosus* IL144 DNA | NC_017075 | 5043253 | Proteobacteria; Betaproteobacteria; Burkholderiales; Rubrivivax |
| *Paenibacillus* sp. JDR-2 | NC_012914 | 7184930 | Firmicutes; Bacilli; Bacillales; Paenibacillaceae |
| *Desulfobacterium autotrophicum* HRM2 | NC_012108 | 5589073 | Proteobacteria; Deltaproteobacteria; Desulfobacterales; Desulfobacteraceae; Desulfobacterium |
| *Thauera* sp. MZ1T | NC_011662 | 4496212 | Proteobacteria; Deltaproteobacteria; Desulfobacterales; Desulfobacteraceae; Desulfobacterium |
| *Desulfovibrio vulgaris* str. 'Miyazaki F' | NC_011769 | 4040304 | Proteobacteria; Deltaproteobacteria; Desulfovibrionales; Desulfovibrionaceae; Desulfovibrio |
| *Desulfovibrio vulgaris* RCH1 | NC_017310 | 3532052 | Proteobacteria; Deltaproteobacteria; Desulfovibrionales; Desulfovibrionaceae; Desulfovibrio |
| *Geobacter metallireducens* GS-15 | NC_007517 | 3997420 | Proteobacteria; Deltaproteobacteria; Desulfuromonadales; Geobacteraceae; Geobacter |
| *Myxococcus xanthus* DK 1622 | NC_008095 | 9139763 | Proteobacteria; Deltaproteobacteria; Myxococcales; Cystobacterineae; Myxococcaceae; Myxococcus. |
| *Sorangium cellulosum* 'So ce 56' | NC_010162 | 13033779 | Proteobacteria; Deltaproteobacteria; Myxococcales; Sorangiineae; Polyangiaceae; Sorangium |
| *Marinobacter aquaeolei* VT8 | NC_008740 | 4326849 | Proteobacteria; Gammaproteobacteria; Alteromonadales; Alteromonadaceae; Marinobacter. |
| *Pseudoalteromonas atlantica* T6c | NC_008228 | 5187005 | Proteobacteria; Gammaproteobacteria; Alteromonadales; Pseudoalteromonadaceae; Pseudoalteromonas. |
| *Shewanella baltica* OS195 | NC_009997 | 5347283 | Proteobacteria; Gammaproteobacteria; Alteromonadales; Shewanellaceae; Shewanella |
| *Escherichia coli* str. K12 substr. DH10B | NC_010473 | 4686137 | Proteobacteria; Gammaproteobacteria; Enterobacteriales; Enterobacteriaceae; Escherichia |
| *Acinetobacter calcoaceticus* PHEA-2 | NC_016603 | 3862530 | Proteobacteria; Gammaproteobacteria; Pseudomonadales; Moraxellaceae; Acinetobacter; Acinetobacter calcoaceticus/baumannii |
| *Pseudomonas putida* KT2440 | NC_002947 | 6181863 | Proteobacteria; Gammaproteobacteria; Pseudomonadales; Pseudomonadaceae; Pseudomonas |
| *Vibrio harveyi* ATCC BAA-1116 | NC_009783 | 3765351 | Proteobacteria; Gammaproteobacteria; Vibrionales; Vibrionaceae; Vibrio |
| *Xanthomonas axonopodis* Xac29-1 | NC_020800 | 5153455 | Proteobacteria; Gammaproteobacteria; Xanthomonadales; Xanthomonadaceae; Xanthomonas |
| *Leptospira interrogans* serovar Copenhageni str. Fiocruz L1-130 | NC_005823 | 4277185 | Spirochaetes; Leptospirales; Leptospiraceae; Leptospira |
| *Thermotoga maritima* MSB8 | NC_000853 | 1860725 | Thermotogae; Thermotogales; Thermotogaceae; Thermotoga |
| *Desulfotomaculum acetoxidans* DSM 771 | NC_013216 | 4545624 | Firmicutes; Clostridia; Clostridiales; Peptococcaceae; Desulfotomaculum |
| *Mycobacterium abscessus* subsp. *bolletii* 50594 | NC_021282.1 | 5000473 | Actinobacteria; Corynebacteriales; Mycobacteriaceae; Mycobacterium; Mycobacterium abscessus |
| *Rhodococcus erythropolis* CCM2595 | NC_022115 | 6281198 | Actinobacteria; Corynebacteriales; Nocardiaceae; Rhodococcus |
| *Frankia* sp. EuI1c | NC_014666 | 8815781 | Actinobacteria; Frankiales; Frankiaceae; Frankia |
| *Streptomyces coelicolor* A3 | NC_003888 | 8667507 | Actinobacteria; Streptomycetales; Streptomycetaceae; Streptomyces; Streptomyces albidoflavus group |
| *Aquifex aeolicus* VF5 | NC_000918 | 1551335 | Aquificae; Aquificales; Aquificaceae; Aquifex |
| *Sulfurihydrogenibium* sp. YO3AOP1 | NC_010730 | 1838442 | Aquificae; Aquificales; Hydrogenothermaceae; Sulfurihydrogenibium |
| *Bacteroides thetaiotaomicron* VPI-5482 | NC_004663 | 6260361 | Bacteroidetes; Bacteroidia; Bacteroidales; Bacteroidaceae; Bacteroides |
| *Chlorobium tepidum* TLS | NC_002932 | 2154946 | Chlorobi; Chlorobia; Chlorobiales; Chlorobiaceae; Chlorobaculum |
| *Chlorobium limicola* DSM 245 | NC_010803 | 2763181 | Chlorobi; Chlorobia; Chlorobiales; Chlorobiaceae; Chlorobium/Pelodictyon group; Chlorobium |
| *Chloroherpeton thalassium* ATCC 35110 | NC_011026 | 3293456 | Chlorobi; Chlorobia; Chlorobiales; Chlorobiaceae; Chloroherpeton |
| *Prosthecochloris aestuarii* DSM 271 | NC_011059 | 2512923 | Chlorobi; Chlorobia; Chlorobiales; Chlorobiaceae; Prosthecochloris |
| *Chloroflexus aurantiacus* J-10-fl | NC_010175 | 5258541 | Chloroflexi; Chloroflexia; Chloroflexales; Chloroflexineae; Chloroflexaceae; Chloroflexus. |
| *Roseiflexus castenholzii* DSM 13941 | CP000804 | 5723298 | Chloroflexi; Chloroflexia; Chloroflexales; Roseiflexineae; Roseiflexaceae; Roseiflexus |
