## Supplemental for "Whole genome phylogeny of Cyanobacteria documents a distinct evolutionary trajectory of marine picocyanobacteria": Supplementary_Table_S3 .docx

| Supplementary_Table_S3 – List of genes used for phylogeny of photosynthesis. | |
| --- | --- |
| Genbank accession numbers in genome of *P.* *marinus* MIT 9313 | protein name |
| 895817 | photosystem I assembly protein Ycf3 |
| 895008 | photosystem I assembly protein Ycf4 |
| 893887 | photosystem I assembly-like protein Ycf37 |
| 895597 | photosystem I P700 chlorophyll a apoprotein A1 |
| 895596 | photosystem I P700 chlorophyll a apoprotein A2 |
| 895537 | photosystem I protein PsaD |
| 895144 | photosystem I PsaF protein (subunit III) |
| 895595 | photosystem I reaction center protein subunit XI |
| 894020 | photosystem I reaction center subunit IV |
| 895779 | photosystem I subunit VII |
| 894909 | photosystem II oxygen evolving complex protein PsbP |
| 895359 | photosystem II PsbA protein (D1) |
| 895492 | photosystem II PsbB protein (CP47) |
| 895010 | photosystem II PsbC protein (CP43) |
| 895009 | photosystem II PsbD protein (D2) |
| 894437 | photosystem II reaction center protein Psb28 |
| 895088 | photosystem II reaction center Psb27 protein |
