## Supplemental for "Whole genome phylogeny of Cyanobacteria documents a distinct evolutionary trajectory of marine picocyanobacteria": Supplementary_Table_S4 .docx

| Supplementary_Table_S4 – List of genes found in all studied genomes. | |
| --- | --- |
| KO | Gene Definition |
| K02469 | gyrA; DNA gyrase subunit A [EC:5.99.1.3] |
| K00150 | gap2; glyceraldehyde-3-phosphate dehydrogenase (NAD(P)) [EC:1.2.1.59] |
| K01951 | guaA; GMP synthase (glutamine-hydrolysing) [EC:6.3.5.2] |
| K02343 | DPO3G; DNA polymerase III subunit gamma/tau [EC:2.7.7.7] |
| K03544 | clpX; ATP-dependent Clp protease ATP-binding subunit ClpX |
| K01358 | clpP; ATP-dependent Clp protease, protease subunit [EC:3.4.21.92] |
| K03070 | secA; preprotein translocase subunit SecA |
| K02520 | infC; translation initiation factor IF-3 |
| K02470 | gyrB; DNA gyrase subunit B [EC:5.99.1.3] |
| K14441 | rimO; ribosomal protein S12 methylthiotransferase [EC:2.8.4.4] |
| K01738 | cysK; cysteine synthase A [EC:2.5.1.47] |
| K02986 | RP-S4; small subunit ribosomal protein S4 |
| K01358 | clpP; ATP-dependent Clp protease, protease subunit [EC:3.4.21.92] |
| K01358 | clpP; ATP-dependent Clp protease, protease subunit [EC:3.4.21.92] |
| K06168 | miaB; tRNA-2-methylthio-N6-dimethylallyladenosine synthase [EC:2.8.4.3] |
| K00566 | mnmA; tRNA-uridine 2-sulfurtransferase [EC:2.8.1.13] |
| K00554 | trmD; tRNA (guanine37-N1)-methyltransferase [EC:2.1.1.228] |
| K11753 | ribF; riboflavin kinase / FMN adenylyltransferase [EC:2.7.1.26 2.7.7.2] |
| K01889 | FARSA; phenylalanyl-tRNA synthetase alpha chain [EC:6.1.1.20] |
| K01462 | PDF; peptide deformylase [EC:3.5.1.88] |
| K01867 | WARS; tryptophanyl-tRNA synthetase [EC:6.1.1.2] |
| K01868 | TARS; threonyl-tRNA synthetase [EC:6.1.1.3] |
| K01783 | rpe; ribulose-phosphate 3-epimerase [EC:5.1.3.1] |
| K03655 | recG; ATP-dependent DNA helicase RecG [EC:3.6.4.12] |
| K02357 | tsf; elongation factor Ts |
| K02967 | RP-S2; small subunit ribosomal protein S2 |
| K03072 | secD; preprotein translocase subunit SecD |
| K02337 | DPO3A1; DNA polymerase III subunit alpha [EC:2.7.7.7] |
| K01956 | carA; carbamoyl-phosphate synthase small subunit [EC:6.3.5.5] |
| K01358 | clpP; ATP-dependent Clp protease, protease subunit [EC:3.4.21.92] |
| K04043 | dnaK; molecular chaperone DnaK |
| K01869 | LARS; leucyl-tRNA synthetase [EC:6.1.1.4] |
| K00954 | E2.7.7.3A; pantetheine-phosphate adenylyltransferase [EC:2.7.7.3] |
| K03703 | uvrC; excinuclease ABC subunit C |
| K01890 | FARSB; phenylalanyl-tRNA synthetase beta chain [EC:6.1.1.20] |
| K01874 | MARS; methionyl-tRNA synthetase [EC:6.1.1.10] |
| K03723 | mfd; transcription-repair coupling factor (superfamily II helicase) [EC:3.6.4.-] |
| K01803 | TPI; triosephosphate isomerase (TIM) [EC:5.3.1.1] |
| K01955 | carB; carbamoyl-phosphate synthase large subunit [EC:6.3.5.5] |
| K00134 | GAPDH; glyceraldehyde 3-phosphate dehydrogenase [EC:1.2.1.12] |
| K00806 | uppS; undecaprenyl diphosphate synthase [EC:2.5.1.31] |
| K00948 | PRPS; ribose-phosphate pyrophosphokinase [EC:2.7.6.1] |
| K13789 | GGPS; geranylgeranyl diphosphate synthase, type II [EC:2.5.1.1 2.5.1.10 2.5.1.29] |
| K06177 | rluA; tRNA pseudouridine32 synthase / 23S rRNA pseudouridine746 synthase [EC:5.4.99.28 5.4.99.29] |
| K02469 | gyrA; DNA gyrase subunit A [EC:5.99.1.3] |
| K00088 | IMPDH; IMP dehydrogenase [EC:1.1.1.205] |
| K06942 | ychF; ribosome-binding ATPase |
| K01883 | CARS; cysteinyl-tRNA synthetase [EC:6.1.1.16] |
| K02838 | frr; ribosome recycling factor |
| K01265 | map; methionyl aminopeptidase [EC:3.4.11.18] |
| K02884 | RP-L19; large subunit ribosomal protein L19 |
| K01885 | EARS; glutamyl-tRNA synthetase [EC:6.1.1.17] |
| K04077 | groEL; chaperonin GroEL |
| K03168 | topA; DNA topoisomerase I [EC:5.99.1.2] |
| K01875 | SARS; seryl-tRNA synthetase [EC:6.1.1.11] |
| K00962 | pnp; polyribonucleotide nucleotidyltransferase [EC:2.7.7.8] |
| K04077 | groEL; chaperonin GroEL |
| K02112 | ATPF1B; F-type H+-transporting ATPase subunit beta [EC:3.6.3.14] |
| K02111 | ATPF1A; F-type H+-transporting ATPase subunit alpha [EC:3.6.3.14] |
| K03046 | rpoC; DNA-directed RNA polymerase subunit beta' [EC:2.7.7.6] |
| K03046 | rpoC; DNA-directed RNA polymerase subunit beta' [EC:2.7.7.6] |
| K03043 | rpoB; DNA-directed RNA polymerase subunit beta [EC:2.7.7.6] |
| K02600 | nusA; N utilization substance protein A |
| K02519 | infB; translation initiation factor IF-2 |
| K03979 | obgE; GTPase [EC:3.6.5.-] |
| K01462 | PDF; peptide deformylase [EC:3.5.1.88] |
| K13799 | panC-cmk; pantoate ligase / CMP/dCMP kinase [EC:6.3.2.1 2.7.4.25] |
| K03553 | recA; recombination protein RecA |
| K02906 | RP-L3; large subunit ribosomal protein L3 |
| K02886 | RP-L2; large subunit ribosomal protein L2 |
| K02965 | RP-S19; small subunit ribosomal protein S19 |
| K02982 | RP-S3; small subunit ribosomal protein S3 |
| K02878 | RP-L16; large subunit ribosomal protein L16 |
| K02874 | RP-L14; large subunit ribosomal protein L14 |
| K02931 | RP-L5; large subunit ribosomal protein L5 |
| K02994 | RP-S8; small subunit ribosomal protein S8 |
| K02933 | RP-L6; large subunit ribosomal protein L6 |
| K02881 | RP-L18; large subunit ribosomal protein L18 |
| K02988 | RP-S5; small subunit ribosomal protein S5 |
| K03076 | secY; preprotein translocase subunit SecY |
| K00939 | adk; adenylate kinase [EC:2.7.4.3] |
| K02952 | RP-S13; small subunit ribosomal protein S13 |
| K02948 | RP-S11; small subunit ribosomal protein S11 |
| K03040 | rpoA; DNA-directed RNA polymerase subunit alpha [EC:2.7.7.6] |
| K02871 | RP-L13; large subunit ribosomal protein L13 |
| K02996 | RP-S9; small subunit ribosomal protein S9 |
| K02835 | prfA; peptide chain release factor 1 |
| K02950 | RP-S12; small subunit ribosomal protein S12 |
| K02946 | RP-S10; small subunit ribosomal protein S10 |
| K01736 | aroC; chorismate synthase [EC:4.2.3.5] |
| K00609 | pyrB; aspartate carbamoyltransferase catalytic subunit [EC:2.1.3.2] |
| K01870 | IARS; isoleucyl-tRNA synthetase [EC:6.1.1.5] |
| K03657 | uvrD; DNA helicase II / ATP-dependent DNA helicase PcrA [EC:3.6.4.12] |
| K00820 | glmS; glucosamine--fructose-6-phosphate aminotransferase (isomerizing) [EC:2.6.1.16] |
| K00615 | E2.2.1.1; transketolase [EC:2.2.1.1] |
| K01738 | cysK; cysteine synthase A [EC:2.5.1.47] |
| K04487 | iscS; cysteine desulfurase [EC:2.8.1.7] |
| K03438 | mraW; 16S rRNA (cytosine1402-N4)-methyltransferase [EC:2.1.1.199] |
| K02836 | prfB; peptide chain release factor 2 |
| K01689 | ENO; enolase [EC:4.2.1.11] |
| K00927 | PGK; phosphoglycerate kinase [EC:2.7.2.3] |
| K06180 | rluD; 23S rRNA pseudouridine1911/1915/1917 synthase [EC:5.4.99.23] |
| K01872 | AARS; alanyl-tRNA synthetase [EC:6.1.1.7] |
| K02314 | dnaB; replicative DNA helicase [EC:3.6.4.12] |
| K01873 | VARS; valyl-tRNA synthetase [EC:6.1.1.9] |
| K01876 | DARS; aspartyl-tRNA synthetase [EC:6.1.1.12] |
| K04043 | dnaK; molecular chaperone DnaK |
| K01000 | mraY; phospho-N-acetylmuramoyl-pentapeptide-transferase [EC:2.7.8.13] |
| K03701 | uvrA; excinuclease ABC subunit A |
