## Supplemental for "Whole genome phylogeny of Cyanobacteria documents a distinct evolutionary trajectory of marine picocyanobacteria": Supplementary_Table_S5.docx

Supplementary_Table_S5- Phylogenetic relation of 673 evaluated genes. Highlighted text indicate potential HGT from reference bacteria to *Gloeobacter.*

| **Group #** | **Group description** | **Protein identification** |
| --- | --- | --- |
| 1 | Gloobacter basal to PSC clade | gi\|33862288\|ref\|NP_893848.1\| tRNA-dihydrouridine synthase A [Prochlorococcus marinus str. MIT 9313] |
| 1 | Gloobacter basal to PSC clade | gi\|33862294\|ref\|NP_893854.1\| heat shock protein GrpE [Prochlorococcus marinus str. MIT 9313] |
| 1 | Gloobacter basal to PSC clade | gi\|33862297\|ref\|NP_893857.1\| hypothetical protein PMT0024 [Prochlorococcus marinus str. MIT 9313] |
| 1 | Gloobacter basal to PSC clade | gi\|33862303\|ref\|NP_893863.1\| cyclophilin type peptidyl-prolyl cis-trans isomerase [Prochlorococcus marinus str. MIT 9313] |
| 1 | Gloobacter basal to PSC clade | gi\|33862316\|ref\|NP_893876.1\| soluble hydrogenase small subunit [Prochlorococcus marinus str. MIT 9313] |
| 1 | Gloobacter basal to PSC clade | gi\|33862321\|ref\|NP_893881.1\| penicillin-binding protein [Prochlorococcus marinus str. MIT 9313] |
| 1 | Gloobacter basal to PSC clade | gi\|33862322\|ref\|NP_893882.1\| SqdX [Prochlorococcus marinus str. MIT 9313] |
| 1 | Gloobacter basal to PSC clade | gi\|33862332\|ref\|NP_893892.1\| DNA polymerase III subunits gamma/tau [Prochlorococcus marinus str. MIT 9313] |
| 1 | Gloobacter basal to PSC clade | gi\|33862334\|ref\|NP_893894.1\| ATP-dependent protease ATP-binding subunit ClpX [Prochlorococcus marinus str. MIT 9313] |
| 1 | Gloobacter basal to PSC clade | gi\|33862335\|ref\|NP_893895.1\| Clp protease proteolytic subunit [Prochlorococcus marinus str. MIT 9313] |
| 1 | Gloobacter basal to PSC clade | gi\|33862356\|ref\|NP_893916.1\| preprotein translocase subunit SecA [Prochlorococcus marinus str. MIT 9313] |
| 1 | Gloobacter basal to PSC clade | gi\|33862369\|ref\|NP_893929.1\| capsular polysaccharide biosynthesis protein [Prochlorococcus marinus str. MIT 9313] |
| 1 | Gloobacter basal to PSC clade | gi\|33862376\|ref\|NP_893936.1\| aminotransferase, class III pyridoxal-phosphate dependent [Prochlorococcus marinus str. MIT 9313] |
| 1 | Gloobacter basal to PSC clade | gi\|33862385\|ref\|NP_893945.1\| glucose-1-phosphate thymidylyltransferase [Prochlorococcus marinus str. MIT 9313] |
| 1 | Gloobacter basal to PSC clade | gi\|33862392\|ref\|NP_893952.1\| translation initiation factor IF-3 [Prochlorococcus marinus str. MIT 9313] |
| 1 | Gloobacter basal to PSC clade | gi\|33862394\|ref\|NP_893954.1\| DNA gyrase subunit B [Prochlorococcus marinus str. MIT 9313] |
| 1 | Gloobacter basal to PSC clade | gi\|33862398\|ref\|NP_893958.1\| glutathione peroxidase [Prochlorococcus marinus str. MIT 9313] |
| 1 | Gloobacter basal to PSC clade | gi\|33862399\|ref\|NP_893959.1\| Mg2+ transporter [Prochlorococcus marinus str. MIT 9313] |
| 1 | Gloobacter basal to PSC clade | gi\|33862408\|ref\|NP_893968.1\| adenine glycosylase [Prochlorococcus marinus str. MIT 9313] |
| 1 | Gloobacter basal to PSC clade | gi\|33862414\|ref\|NP_893974.1\| single-stranded DNA-binding protein [Prochlorococcus marinus str. MIT 9313] |
| 1 | Gloobacter basal to PSC clade | gi\|33862415\|ref\|NP_893975.1\| rod shape-determining protein MreB [Prochlorococcus marinus str. MIT 9313] |
| 1 | Gloobacter basal to PSC clade | gi\|33862420\|ref\|NP_893980.1\| lysyl-tRNA synthetase [Prochlorococcus marinus str. MIT 9313] |
| 1 | Gloobacter basal to PSC clade | gi\|33862443\|ref\|NP_894003.1\| hypothetical protein PMT0170 [Prochlorococcus marinus str. MIT 9313] |
| 1 | Gloobacter basal to PSC clade | gi\|33862456\|ref\|NP_894016.1\| inositol monophosphate family protein [Prochlorococcus marinus str. MIT 9313] |
| 1 | Gloobacter basal to PSC clade | gi\|33862459\|ref\|NP_894019.1\| formamidopyrimidine-DNA glycosylase [Prochlorococcus marinus str. MIT 9313] |
| 1 | Gloobacter basal to PSC clade | gi\|33862462\|ref\|NP_894022.1\| DEAD/DEAH box helicase [Prochlorococcus marinus str. MIT 9313] |
| 1 | Gloobacter basal to PSC clade | gi\|33862480\|ref\|NP_894040.1\| hypothetical protein PMT0207 [Prochlorococcus marinus str. MIT 9313] |
| 1 | Gloobacter basal to PSC clade | gi\|33862487\|ref\|NP_894047.1\| lipoate-protein ligase B [Prochlorococcus marinus str. MIT 9313] |
| 1 | Gloobacter basal to PSC clade | gi\|33862493\|ref\|NP_894053.1\| branched-chain alpha-keto acid dehydrogenase subunit E2 [Prochlorococcus marinus str. MIT 9313] |
| 1 | Gloobacter basal to PSC clade | gi\|33862500\|ref\|NP_894060.1\| 30S ribosomal protein S4 [Prochlorococcus marinus str. MIT 9313] |
| 1 | Gloobacter basal to PSC clade | gi\|33862506\|ref\|NP_894066.1\| L-cysteine/cystine lyase [Prochlorococcus marinus str. MIT 9313] |
| 1 | Gloobacter basal to PSC clade | gi\|33862541\|ref\|NP_894101.1\| tRNA/rRNA methyltransferase SpoU [Prochlorococcus marinus str. MIT 9313] |
| 1 | Gloobacter basal to PSC clade | gi\|33862587\|ref\|NP_894147.1\| ATP-dependent Clp protease proteolytic subunit [Prochlorococcus marinus str. MIT 9313] |
| 1 | Gloobacter basal to PSC clade | gi\|33862588\|ref\|NP_894148.1\| ATP-dependent Clp protease-like protein [Prochlorococcus marinus str. MIT 9313] |
| 1 | Gloobacter basal to PSC clade | gi\|33862595\|ref\|NP_894155.1\| D-alanyl-alanine synthetase A [Prochlorococcus marinus str. MIT 9313] |
| 1 | Gloobacter basal to PSC clade | gi\|33862596\|ref\|NP_894156.1\| (dimethylallyl)adenosine tRNA methylthiotransferase [Prochlorococcus marinus str. MIT 9313] |
| 1 | Gloobacter basal to PSC clade | gi\|33862623\|ref\|NP_894183.1\| 30S ribosomal protein S16 [Prochlorococcus marinus str. MIT 9313] |
| 1 | Gloobacter basal to PSC clade | gi\|33862639\|ref\|NP_894199.1\| stationary phase survival protein SurE [Prochlorococcus marinus str. MIT 9313] |
| 1 | Gloobacter basal to PSC clade | gi\|33862660\|ref\|NP_894220.1\| bacterioferritin comigratory protein [Prochlorococcus marinus str. MIT 9313] |
| 1 | Gloobacter basal to PSC clade | gi\|33862667\|ref\|NP_894227.1\| cell wall hydrolase/autolysin [Prochlorococcus marinus str. MIT 9313] |
| 1 | Gloobacter basal to PSC clade | gi\|33862695\|ref\|NP_894255.1\| tryptophanyl-tRNA synthetase [Prochlorococcus marinus str. MIT 9313] |
| 1 | Gloobacter basal to PSC clade | gi\|33862698\|ref\|NP_894258.1\| glucokinase [Prochlorococcus marinus str. MIT 9313] |
| 1 | Gloobacter basal to PSC clade | gi\|33862709\|ref\|NP_894269.1\| gamma-glutamyl phosphate reductase [Prochlorococcus marinus str. MIT 9313] |
| 1 | Gloobacter basal to PSC clade | gi\|33862710\|ref\|NP_894270.1\| ROK family protein [Prochlorococcus marinus str. MIT 9313] |
| 1 | Gloobacter basal to PSC clade | gi\|33862717\|ref\|NP_894277.1\| glycogen branching protein [Prochlorococcus marinus str. MIT 9313] |
| 1 | Gloobacter basal to PSC clade | gi\|33862718\|ref\|NP_894278.1\| uroporphyrinogen decarboxylase [Prochlorococcus marinus str. MIT 9313] |
| 1 | Gloobacter basal to PSC clade | gi\|33862726\|ref\|NP_894286.1\| bifunctional phosphoribosyl-AMP cyclohydrolase/phosphoribosyl-ATP pyrophosphatase [Prochlorococcus marinus str. MIT 9313] |
| 1 | Gloobacter basal to PSC clade | gi\|33862803\|ref\|NP_894363.1\| UDP-3-O-[3-hydroxymyristoyl] glucosamine N-acyltransferase [Prochlorococcus marinus str. MIT 9313] |
| 1 | Gloobacter basal to PSC clade | gi\|33862810\|ref\|NP_894370.1\| fructose-1,6-bisphosphate aldolase [Prochlorococcus marinus str. MIT 9313] |
| 1 | Gloobacter basal to PSC clade | gi\|33862839\|ref\|NP_894399.1\| glucose-1-phosphate adenylyltransferase [Prochlorococcus marinus str. MIT 9313] |
| 1 | Gloobacter basal to PSC clade | gi\|33862852\|ref\|NP_894412.1\| sulfite reductase subunit beta [Prochlorococcus marinus str. MIT 9313] |
| 1 | Gloobacter basal to PSC clade | gi\|33862856\|ref\|NP_894416.1\| elongation factor Ts [Prochlorococcus marinus str. MIT 9313] |
| 1 | Gloobacter basal to PSC clade | gi\|33862857\|ref\|NP_894417.1\| 30S ribosomal protein S2 [Prochlorococcus marinus str. MIT 9313] |
| 1 | Gloobacter basal to PSC clade | gi\|33862861\|ref\|NP_894421.1\| ABC transporter transmembrane protein [Prochlorococcus marinus str. MIT 9313] |
| 1 | Gloobacter basal to PSC clade | gi\|33862871\|ref\|NP_894431.1\| pyridoxal-dependent decarboxylase family protein [Prochlorococcus marinus str. MIT 9313] |
| 1 | Gloobacter basal to PSC clade | gi\|33862917\|ref\|NP_894477.1\| Holliday junction DNA helicase RuvA [Prochlorococcus marinus str. MIT 9313] |
| 1 | Gloobacter basal to PSC clade | gi\|33862926\|ref\|NP_894486.1\| carbamoyl phosphate synthase small subunit [Prochlorococcus marinus str. MIT 9313] |
| 1 | Gloobacter basal to PSC clade | gi\|33862941\|ref\|NP_894501.1\| phosphoribosylaminoimidazole carboxylase ATPase subunit [Prochlorococcus marinus str. MIT 9313] |
| 1 | Gloobacter basal to PSC clade | gi\|33862948\|ref\|NP_894508.1\| ATP-dependent Clp protease proteolytic subunit [Prochlorococcus marinus str. MIT 9313] |
| 1 | Gloobacter basal to PSC clade | gi\|33862950\|ref\|NP_894510.1\| ABC transporter [Prochlorococcus marinus str. MIT 9313] |
| 1 | Gloobacter basal to PSC clade | gi\|33862996\|ref\|NP_894556.1\| excinuclease ABC subunit C [Prochlorococcus marinus str. MIT 9313] |
| 1 | Gloobacter basal to PSC clade | gi\|33863000\|ref\|NP_894560.1\| branched-chain amino acid aminotransferase [Prochlorococcus marinus str. MIT 9313] |
| 1 | Gloobacter basal to PSC clade | gi\|33863012\|ref\|NP_894572.1\| 30S ribosomal protein S18 [Prochlorococcus marinus str. MIT 9313] |
| 1 | Gloobacter basal to PSC clade | gi\|33863042\|ref\|NP_894602.1\| modulator of DNA gyrase TldD [Prochlorococcus marinus str. MIT 9313] |
| 1 | Gloobacter basal to PSC clade | gi\|33863049\|ref\|NP_894609.1\| 4-hydroxy-3-methylbut-2-en-1-yl diphosphate synthase [Prochlorococcus marinus str. MIT 9313] |
| 1 | Gloobacter basal to PSC clade | gi\|33863058\|ref\|NP_894618.1\| 16S ribosomal RNA methyltransferase RsmE [Prochlorococcus marinus str. MIT 9313] |
| 1 | Gloobacter basal to PSC clade | gi\|33863087\|ref\|NP_894647.1\| magnesium chelatase subunit H [Prochlorococcus marinus str. MIT 9313] |
| 1 | Gloobacter basal to PSC clade | gi\|33863287\|ref\|NP_894847.1\| hypothetical protein PMT1016 [Prochlorococcus marinus str. MIT 9313] |
| 1 | Gloobacter basal to PSC clade | gi\|33863322\|ref\|NP_894882.1\| carotenoid isomerase [Prochlorococcus marinus str. MIT 9313] |
| 1 | Gloobacter basal to PSC clade | gi\|33863330\|ref\|NP_894890.1\| diaminopimelate decarboxylase [Prochlorococcus marinus str. MIT 9313] |
| 1 | Gloobacter basal to PSC clade | gi\|33863331\|ref\|NP_894891.1\| ribosomal-protein-alanine acetyltransferase [Prochlorococcus marinus str. MIT 9313] |
| 1 | Gloobacter basal to PSC clade | gi\|33863345\|ref\|NP_894905.1\| ribose-phosphate pyrophosphokinase [Prochlorococcus marinus str. MIT 9313] |
| 1 | Gloobacter basal to PSC clade | gi\|33863348\|ref\|NP_894908.1\| recombination protein RecR [Prochlorococcus marinus str. MIT 9313] |
| 1 | Gloobacter basal to PSC clade | gi\|33863352\|ref\|NP_894912.1\| RNA-binding protein RbpD [Prochlorococcus marinus str. MIT 9313] |
| 1 | Gloobacter basal to PSC clade | gi\|33863356\|ref\|NP_894916.1\| UvrD/REP helicase [Prochlorococcus marinus str. MIT 9313] |
| 1 | Gloobacter basal to PSC clade | gi\|33863366\|ref\|NP_894926.1\| glutaredoxin-like protein [Prochlorococcus marinus str. MIT 9313] |
| 1 | Gloobacter basal to PSC clade | gi\|33863395\|ref\|NP_894955.1\| DNA gyrase subunit A [Prochlorococcus marinus str. MIT 9313] |
| 1 | Gloobacter basal to PSC clade | gi\|33863406\|ref\|NP_894966.1\| protoporphyrin IX magnesium chelatase subunit ChlI [Prochlorococcus marinus str. MIT 9313] |
| 1 | Gloobacter basal to PSC clade | gi\|33863414\|ref\|NP_894974.1\| homoserine dehydrogenase [Prochlorococcus marinus str. MIT 9313] |
| 1 | Gloobacter basal to PSC clade | gi\|33863429\|ref\|NP_894989.1\| cysteinyl-tRNA synthetase [Prochlorococcus marinus str. MIT 9313] |
| 1 | Gloobacter basal to PSC clade | gi\|161350044\|ref\|NP_894991.2\| 1-deoxy-D-xylulose 5-phosphate reductoisomerase [Prochlorococcus marinus str. MIT 9313] |
| 1 | Gloobacter basal to PSC clade | gi\|33863441\|ref\|NP_895001.1\| methyltransferase [Prochlorococcus marinus str. MIT 9313] |
| 1 | Gloobacter basal to PSC clade | gi\|33863451\|ref\|NP_895011.1\| Maf-like protein [Prochlorococcus marinus str. MIT 9313] |
| 1 | Gloobacter basal to PSC clade | gi\|33863463\|ref\|NP_895023.1\| ATP phosphoribosyltransferase catalytic subunit [Prochlorococcus marinus str. MIT 9313] |
| 1 | Gloobacter basal to PSC clade | gi\|33863509\|ref\|NP_895069.1\| phage integrase family protein [Prochlorococcus marinus str. MIT 9313] |
| 1 | Gloobacter basal to PSC clade | gi\|33863514\|ref\|NP_895074.1\| NAD binding site [Prochlorococcus marinus str. MIT 9313] |
| 1 | Gloobacter basal to PSC clade | gi\|33863517\|ref\|NP_895077.1\| peptidoglycan synthetase [Prochlorococcus marinus str. MIT 9313] |
| 1 | Gloobacter basal to PSC clade | gi\|33863520\|ref\|NP_895080.1\| alpha-ribazole-5'-phosphate phosphatase [Prochlorococcus marinus str. MIT 9313] |
| 1 | Gloobacter basal to PSC clade | gi\|33863541\|ref\|NP_895101.1\| porphobilinogen deaminase [Prochlorococcus marinus str. MIT 9313] |
| 1 | Gloobacter basal to PSC clade | gi\|33863562\|ref\|NP_895122.1\| exodeoxyribonuclease III [Prochlorococcus marinus str. MIT 9313] |
| 1 | Gloobacter basal to PSC clade | gi\|33863569\|ref\|NP_895129.1\| hypothetical protein PMT1301 [Prochlorococcus marinus str. MIT 9313] |
| 1 | Gloobacter basal to PSC clade | gi\|33863573\|ref\|NP_895133.1\| methionine aminopeptidase [Prochlorococcus marinus str. MIT 9313] |
| 1 | Gloobacter basal to PSC clade | gi\|33863574\|ref\|NP_895134.1\| 50S ribosomal protein L19 [Prochlorococcus marinus str. MIT 9313] |
| 1 | Gloobacter basal to PSC clade | gi\|33863584\|ref\|NP_895144.1\| photosystem I PsaF protein (subunit III) [Prochlorococcus marinus str. MIT 9313] |
| 1 | Gloobacter basal to PSC clade | gi\|33863586\|ref\|NP_895146.1\| secreted protein MPB70 precursor [Prochlorococcus marinus str. MIT 9313] |
| 1 | Gloobacter basal to PSC clade | gi\|33863589\|ref\|NP_895149.1\| cytochrome b6-f complex iron-sulfur subunit [Prochlorococcus marinus str. MIT 9313] |
| 1 | Gloobacter basal to PSC clade | gi\|33863604\|ref\|NP_895164.1\| multidrug efflux ABC transporter [Prochlorococcus marinus str. MIT 9313] |
| 1 | Gloobacter basal to PSC clade | gi\|33863628\|ref\|NP_895188.1\| chloroplast membrane-associated 30 kD protein [Prochlorococcus marinus str. MIT 9313] |
| 1 | Gloobacter basal to PSC clade | gi\|33863632\|ref\|NP_895192.1\| signal peptide peptidase SppA [Prochlorococcus marinus str. MIT 9313] |
| 1 | Gloobacter basal to PSC clade | gi\|33863638\|ref\|NP_895198.1\| inner membrane protein translocase component YidC [Prochlorococcus marinus str. MIT 9313] |
| 1 | Gloobacter basal to PSC clade | gi\|161350042\|ref\|NP_895201.2\| seryl-tRNA synthetase [Prochlorococcus marinus str. MIT 9313] |
| 1 | Gloobacter basal to PSC clade | gi\|33863644\|ref\|NP_895204.1\| polynucleotide phosphorylase/polyadenylase [Prochlorococcus marinus str. MIT 9313] |
| 1 | Gloobacter basal to PSC clade | gi\|33863669\|ref\|NP_895229.1\| orotidine 5'-phosphate decarboxylase [Prochlorococcus marinus str. MIT 9313] |
| 1 | Gloobacter basal to PSC clade | gi\|33863670\|ref\|NP_895230.1\| tyrosyl-tRNA synthetase [Prochlorococcus marinus str. MIT 9313] |
| 1 | Gloobacter basal to PSC clade | gi\|33863676\|ref\|NP_895236.1\| lipid-A-disaccharide synthase [Prochlorococcus marinus str. MIT 9313] |
| 1 | Gloobacter basal to PSC clade | gi\|33863687\|ref\|NP_895247.1\| 50S ribosomal protein L21 [Prochlorococcus marinus str. MIT 9313] |
| 1 | Gloobacter basal to PSC clade | gi\|33863717\|ref\|NP_895277.1\| co-chaperonin GroES [Prochlorococcus marinus str. MIT 9313] |
| 1 | Gloobacter basal to PSC clade | gi\|33863718\|ref\|NP_895278.1\| F0F1 ATP synthase subunit beta [Prochlorococcus marinus str. MIT 9313] |
| 1 | Gloobacter basal to PSC clade | gi\|33863719\|ref\|NP_895279.1\| F0F1 ATP synthase subunit epsilon [Prochlorococcus marinus str. MIT 9313] |
| 1 | Gloobacter basal to PSC clade | gi\|33863721\|ref\|NP_895281.1\| hypothetical protein PMT1454 [Prochlorococcus marinus str. MIT 9313] |
| 1 | Gloobacter basal to PSC clade | gi\|33863723\|ref\|NP_895283.1\| aminopeptidase P [Prochlorococcus marinus str. MIT 9313] |
| 1 | Gloobacter basal to PSC clade | gi\|33863730\|ref\|NP_895290.1\| hypothetical protein PMT1463 [Prochlorococcus marinus str. MIT 9313] |
| 1 | Gloobacter basal to PSC clade | gi\|33863734\|ref\|NP_895294.1\| F0F1 ATP synthase subunit alpha [Prochlorococcus marinus str. MIT 9313] |
| 1 | Gloobacter basal to PSC clade | gi\|33863739\|ref\|NP_895299.1\| F0F1 ATP synthase subunit A [Prochlorococcus marinus str. MIT 9313] |
| 1 | Gloobacter basal to PSC clade | gi\|33863741\|ref\|NP_895301.1\| methyltransferase [Prochlorococcus marinus str. MIT 9313] |
| 1 | Gloobacter basal to PSC clade | gi\|33863742\|ref\|NP_895302.1\| cell division protein FtsW [Prochlorococcus marinus str. MIT 9313] |
| 1 | Gloobacter basal to PSC clade | gi\|33863751\|ref\|NP_895311.1\| adenylosuccinate lyase [Prochlorococcus marinus str. MIT 9313] |
| 1 | Gloobacter basal to PSC clade | gi\|33863778\|ref\|NP_895338.1\| ribose-5-phosphate isomerase A [Prochlorococcus marinus str. MIT 9313] |
| 1 | Gloobacter basal to PSC clade | gi\|33863782\|ref\|NP_895342.1\| sulfatase [Prochlorococcus marinus str. MIT 9313] |
| 1 | Gloobacter basal to PSC clade | gi\|33863795\|ref\|NP_895355.1\| translation initiation factor IF-2 [Prochlorococcus marinus str. MIT 9313] |
| 1 | Gloobacter basal to PSC clade | gi\|33863811\|ref\|NP_895371.1\| GTPase ObgE [Prochlorococcus marinus str. MIT 9313] |
| 1 | Gloobacter basal to PSC clade | gi\|33863855\|ref\|NP_895415.1\| acetyl-CoA carboxylase biotin carboxylase subunit [Prochlorococcus marinus str. MIT 9313] |
| 1 | Gloobacter basal to PSC clade | gi\|33863870\|ref\|NP_895430.1\| cysteine desulfurase or selenocysteine lyase [Prochlorococcus marinus str. MIT 9313] |
| 1 | Gloobacter basal to PSC clade | gi\|33863873\|ref\|NP_895433.1\| cysteine desulfurase activator complex subunit SufB [Prochlorococcus marinus str. MIT 9313] |
| 1 | Gloobacter basal to PSC clade | gi\|33863915\|ref\|NP_895475.1\| cytochrome b6-f complex subunit IV [Prochlorococcus marinus str. MIT 9313] |
| 1 | Gloobacter basal to PSC clade | gi\|33863934\|ref\|NP_895494.1\| NrdR family transcriptional regulator [Prochlorococcus marinus str. MIT 9313] |
| 1 | Gloobacter basal to PSC clade | gi\|33863935\|ref\|NP_895495.1\| 30S ribosomal protein S1 [Prochlorococcus marinus str. MIT 9313] |
| 1 | Gloobacter basal to PSC clade | gi\|33863952\|ref\|NP_895512.1\| heme oxygenase [Prochlorococcus marinus str. MIT 9313] |
| 1 | Gloobacter basal to PSC clade | gi\|33863973\|ref\|NP_895533.1\| coproporphyrinogen III oxidase [Prochlorococcus marinus str. MIT 9313] |
| 1 | Gloobacter basal to PSC clade | gi\|33863991\|ref\|NP_895551.1\| arogenate dehydrogenase [Prochlorococcus marinus str. MIT 9313] |
| 1 | Gloobacter basal to PSC clade | gi\|33863994\|ref\|NP_895554.1\| recombinase A [Prochlorococcus marinus str. MIT 9313] |
| 1 | Gloobacter basal to PSC clade | gi\|33863999\|ref\|NP_895559.1\| 50S ribosomal protein L3 [Prochlorococcus marinus str. MIT 9313] |
| 1 | Gloobacter basal to PSC clade | gi\|33864001\|ref\|NP_895561.1\| 50S ribosomal protein L23 [Prochlorococcus marinus str. MIT 9313] |
| 1 | Gloobacter basal to PSC clade | gi\|33864002\|ref\|NP_895562.1\| 50S ribosomal protein L2 [Prochlorococcus marinus str. MIT 9313] |
| 1 | Gloobacter basal to PSC clade | gi\|33864004\|ref\|NP_895564.1\| 50S ribosomal protein L22 [Prochlorococcus marinus str. MIT 9313] |
| 1 | Gloobacter basal to PSC clade | gi\|33864005\|ref\|NP_895565.1\| 30S ribosomal protein S3 [Prochlorococcus marinus str. MIT 9313] |
| 1 | Gloobacter basal to PSC clade | gi\|33864009\|ref\|NP_895569.1\| 50S ribosomal protein L14 [Prochlorococcus marinus str. MIT 9313] |
| 1 | Gloobacter basal to PSC clade | gi\|33864010\|ref\|NP_895570.1\| 50S ribosomal protein L24 [Prochlorococcus marinus str. MIT 9313] |
| 1 | Gloobacter basal to PSC clade | gi\|33864011\|ref\|NP_895571.1\| 50S ribosomal protein L5 [Prochlorococcus marinus str. MIT 9313] |
| 1 | Gloobacter basal to PSC clade | gi\|33864013\|ref\|NP_895573.1\| 50S ribosomal protein L6 [Prochlorococcus marinus str. MIT 9313] |
| 1 | Gloobacter basal to PSC clade | gi\|33864015\|ref\|NP_895575.1\| 30S ribosomal protein S5 [Prochlorococcus marinus str. MIT 9313] |
| 1 | Gloobacter basal to PSC clade | gi\|33864016\|ref\|NP_895576.1\| 50S ribosomal protein L15 [Prochlorococcus marinus str. MIT 9313] |
| 1 | Gloobacter basal to PSC clade | gi\|33864017\|ref\|NP_895577.1\| preprotein translocase subunit SecY [Prochlorococcus marinus str. MIT 9313] |
| 1 | Gloobacter basal to PSC clade | gi\|33864020\|ref\|NP_895580.1\| 30S ribosomal protein S13 [Prochlorococcus marinus str. MIT 9313] |
| 1 | Gloobacter basal to PSC clade | gi\|33864028\|ref\|NP_895588.1\| peptide chain release factor 1 [Prochlorococcus marinus str. MIT 9313] |
| 1 | Gloobacter basal to PSC clade | gi\|33864044\|ref\|NP_895604.1\| ferredoxin-dependent glutamate synthase [Prochlorococcus marinus str. MIT 9313] |
| 1 | Gloobacter basal to PSC clade | gi\|33864050\|ref\|NP_895610.1\| 30S ribosomal protein S10 [Prochlorococcus marinus str. MIT 9313] |
| 1 | Gloobacter basal to PSC clade | gi\|33864052\|ref\|NP_895612.1\| SAM-binding motif-containing protein [Prochlorococcus marinus str. MIT 9313] |
| 1 | Gloobacter basal to PSC clade | gi\|33864058\|ref\|NP_895618.1\| L,L-diaminopimelate aminotransferase [Prochlorococcus marinus str. MIT 9313] |
| 1 | Gloobacter basal to PSC clade | gi\|33864061\|ref\|NP_895621.1\| Fe-S oxidoreductase [Prochlorococcus marinus str. MIT 9313] |
| 1 | Gloobacter basal to PSC clade | gi\|33864075\|ref\|NP_895635.1\| methylpurine-DNA glycosylase (MPG) [Prochlorococcus marinus str. MIT 9313] |
| 1 | Gloobacter basal to PSC clade | gi\|33864120\|ref\|NP_895680.1\| ammonium transporter [Prochlorococcus marinus str. MIT 9313] |
| 1 | Gloobacter basal to PSC clade | gi\|33864121\|ref\|NP_895681.1\| 4-hydroxy-3-methylbut-2-enyl diphosphate reductase [Prochlorococcus marinus str. MIT 9313] |
| 1 | Gloobacter basal to PSC clade | gi\|33864145\|ref\|NP_895705.1\| retinal pigment epithelial membrane protein [Prochlorococcus marinus str. MIT 9313] |
| 1 | Gloobacter basal to PSC clade | gi\|33864147\|ref\|NP_895707.1\| enoyl-ACP reductase [Prochlorococcus marinus str. MIT 9313] |
| 1 | Gloobacter basal to PSC clade | gi\|33864153\|ref\|NP_895713.1\| protoporphyrin IX magnesium chelatase subunit ChlD [Prochlorococcus marinus str. MIT 9313] |
| 1 | Gloobacter basal to PSC clade | gi\|33864158\|ref\|NP_895718.1\| NADH dehydrogenase subunit B [Prochlorococcus marinus str. MIT 9313] |
| 1 | Gloobacter basal to PSC clade | gi\|33864159\|ref\|NP_895719.1\| NADH dehydrogenase subunit A [Prochlorococcus marinus str. MIT 9313] |
| 1 | Gloobacter basal to PSC clade | gi\|33864174\|ref\|NP_895734.1\| sugar transferase [Prochlorococcus marinus str. MIT 9313] |
| 1 | Gloobacter basal to PSC clade | gi\|33864191\|ref\|NP_895751.1\| 2-dehydro-3-deoxyphosphooctonate aldolase [Prochlorococcus marinus str. MIT 9313] |
| 1 | Gloobacter basal to PSC clade | gi\|33864222\|ref\|NP_895782.1\| transketolase [Prochlorococcus marinus str. MIT 9313] |
| 1 | Gloobacter basal to PSC clade | gi\|33864228\|ref\|NP_895788.1\| ribosome-binding factor A [Prochlorococcus marinus str. MIT 9313] |
| 1 | Gloobacter basal to PSC clade | gi\|33864229\|ref\|NP_895789.1\| beta-N-acetylglucosaminidase [Prochlorococcus marinus str. MIT 9313] |
| 1 | Gloobacter basal to PSC clade | gi\|33864231\|ref\|NP_895791.1\| uroporphyrinogen III synthase [Prochlorococcus marinus str. MIT 9313] |
| 1 | Gloobacter basal to PSC clade | gi\|33864234\|ref\|NP_895794.1\| hypothetical protein PMT1969 [Prochlorococcus marinus str. MIT 9313] |
| 1 | Gloobacter basal to PSC clade | gi\|33864277\|ref\|NP_895837.1\| sugar-phosphate nucleotidyl transferase [Prochlorococcus marinus str. MIT 9313] |
| 1 | Gloobacter basal to PSC clade | gi\|33864285\|ref\|NP_895845.1\| NADH dehydrogenase subunit H [Prochlorococcus marinus str. MIT 9313] |
| 1 | Gloobacter basal to PSC clade | gi\|33864286\|ref\|NP_895846.1\| citrate synthase [Prochlorococcus marinus str. MIT 9313] |
| 1 | Gloobacter basal to PSC clade | gi\|33864292\|ref\|NP_895852.1\| tryptophan synthase subunit beta [Prochlorococcus marinus str. MIT 9313] |
| 1 | Gloobacter basal to PSC clade | gi\|33864304\|ref\|NP_895864.1\| N-acetylglucosamine-6-phosphate deacetylase [Prochlorococcus marinus str. MIT 9313] |
| 1 | Gloobacter basal to PSC clade | gi\|33864310\|ref\|NP_895870.1\| NifS-like aminotransferase class-V [Prochlorococcus marinus str. MIT 9313] |
| 1 | Gloobacter basal to PSC clade | gi\|33864328\|ref\|NP_895888.1\| peptide chain release factor 2 [Prochlorococcus marinus str. MIT 9313] |
| 1 | Gloobacter basal to PSC clade | gi\|33864333\|ref\|NP_895893.1\| hypothetical protein PMT2068 [Prochlorococcus marinus str. MIT 9313] |
| 1 | Gloobacter basal to PSC clade | gi\|33864351\|ref\|NP_895911.1\| transcription antitermination protein NusG [Prochlorococcus marinus str. MIT 9313] |
| 1 | Gloobacter basal to PSC clade | gi\|33864354\|ref\|NP_895914.1\| 50S ribosomal protein L10 [Prochlorococcus marinus str. MIT 9313] |
| 1 | Gloobacter basal to PSC clade | gi\|33864370\|ref\|NP_895930.1\| phosphoglycerate kinase [Prochlorococcus marinus str. MIT 9313] |
| 1 | Gloobacter basal to PSC clade | gi\|33864372\|ref\|NP_895932.1\| ribosomal biogenesis GTPase [Prochlorococcus marinus str. MIT 9313] |
| 1 | Gloobacter basal to PSC clade | gi\|33864399\|ref\|NP_895959.1\| bifunctional ornithine acetyltransferase/N-acetylglutamate synthase [Prochlorococcus marinus str. MIT 9313] |
| 1 | Gloobacter basal to PSC clade | gi\|33864403\|ref\|NP_895963.1\| aspartyl/glutamyl-tRNA amidotransferase subunit B [Prochlorococcus marinus str. MIT 9313] |
| 1 | Gloobacter basal to PSC clade | gi\|33864412\|ref\|NP_895972.1\| nucleoside diphosphate kinase [Prochlorococcus marinus str. MIT 9313] |
| 1 | Gloobacter basal to PSC clade | gi\|33864430\|ref\|NP_895990.1\| rubredoxin:rubrerythrin:rubredoxin-type Fe(Cys)4 protein [Prochlorococcus marinus str. MIT 9313] |
| 1 | Gloobacter basal to PSC clade | gi\|33864435\|ref\|NP_895995.1\| cystathionine beta-lyase family aluminum resistance protein [Prochlorococcus marinus str. MIT 9313] |
| 1 | Gloobacter basal to PSC clade | gi\|33864436\|ref\|NP_895996.1\| Fatty acid desaturase, type 1 [Prochlorococcus marinus str. MIT 9313] |
| 1 | Gloobacter basal to PSC clade | gi\|33864438\|ref\|NP_895998.1\| Fatty acid desaturase, type 1 [Prochlorococcus marinus str. MIT 9313] |
| 1 | Gloobacter basal to PSC clade | gi\|33864439\|ref\|NP_895999.1\| 50S ribosomal protein L9 [Prochlorococcus marinus str. MIT 9313] |
| 1 | Gloobacter basal to PSC clade | gi\|33864455\|ref\|NP_896015.1\| valyl-tRNA synthetase [Prochlorococcus marinus str. MIT 9313] |
| 1 | Gloobacter basal to PSC clade | gi\|33864459\|ref\|NP_896019.1\| ferredoxin, PetF like protein [Prochlorococcus marinus str. MIT 9313] |
| 1 | Gloobacter basal to PSC clade | gi\|33864472\|ref\|NP_896032.1\| ferric uptake regulator family protein [Prochlorococcus marinus str. MIT 9313] |
| 1 | Gloobacter basal to PSC clade | gi\|33864479\|ref\|NP_896039.1\| glycine cleavage system aminomethyltransferase T [Prochlorococcus marinus str. MIT 9313] |
| 1 | Gloobacter basal to PSC clade | gi\|33864483\|ref\|NP_896043.1\| CTP synthetase [Prochlorococcus marinus str. MIT 9313] |
| 1 | Gloobacter basal to PSC clade | gi\|33864486\|ref\|NP_896046.1\| ATPase [Prochlorococcus marinus str. MIT 9313] |
| 1 | Gloobacter basal to PSC clade | gi\|33864513\|ref\|NP_896073.1\| bifunctional aconitate hydratase 2/2-methylisocitrate dehydratase [Prochlorococcus marinus str. MIT 9313] |
| 1 | Gloobacter basal to PSC clade | gi\|33864517\|ref\|NP_896077.1\| formyltetrahydrofolate deformylase [Prochlorococcus marinus str. MIT 9313] |
| 1 | Gloobacter basal to PSC clade | gi\|33864519\|ref\|NP_896079.1\| molecular chaperone DnaK [Prochlorococcus marinus str. MIT 9313] |
| 1 | Gloobacter basal to PSC clade | gi\|33864532\|ref\|NP_896092.1\| sucrose phosphate synthase [Prochlorococcus marinus str. MIT 9313] |
| 1 | Gloobacter basal to PSC clade | gi\|33864534\|ref\|NP_896094.1\| excinuclease ABC subunit A [Prochlorococcus marinus str. MIT 9313] |
| 1 | Gloobacter basal to PSC clade | gi\|33864538\|ref\|NP_896098.1\| threonine synthase [Prochlorococcus marinus str. MIT 9313] |
| 3 | NO eubacterial outgroup | gi\|33862278\|ref\|NP_893838.1\| DNA gyrase/topoisomerase IV subunit A [Prochlorococcus marinus str. MIT 9313] |
| 3 | NO eubacterial outgroup | gi\|33862279\|ref\|NP_893839.1\| hypothetical protein PMT0006 [Prochlorococcus marinus str. MIT 9313] |
| 3 | NO eubacterial outgroup | gi\|33862283\|ref\|NP_893843.1\| transcription antitermination protein NusB [Prochlorococcus marinus str. MIT 9313] |
| 3 | NO eubacterial outgroup | gi\|33862327\|ref\|NP_893887.1\| photosystem I assembly-like protein Ycf37 [Prochlorococcus marinus str. MIT 9313] |
| 3 | NO eubacterial outgroup | gi\|33862342\|ref\|NP_893902.1\| hypothetical protein PMT0069 [Prochlorococcus marinus str. MIT 9313] |
| 3 | NO eubacterial outgroup | gi\|33862349\|ref\|NP_893909.1\| transporter component [Prochlorococcus marinus str. MIT 9313] |
| 3 | NO eubacterial outgroup | gi\|33862351\|ref\|NP_893911.1\| hypothetical protein PMT0078 [Prochlorococcus marinus str. MIT 9313] |
| 3 | NO eubacterial outgroup | gi\|33862355\|ref\|NP_893915.1\| acetyltransferase, GNAT family [Prochlorococcus marinus str. MIT 9313] |
| 3 | NO eubacterial outgroup | gi\|33862367\|ref\|NP_893927.1\| transporter for efflux [Prochlorococcus marinus str. MIT 9313] |
| 3 | NO eubacterial outgroup | gi\|33862379\|ref\|NP_893939.1\| N-acetylneuraminate synthase [Prochlorococcus marinus str. MIT 9313] |
| 3 | NO eubacterial outgroup | gi\|33862382\|ref\|NP_893942.1\| ABC transporter [Prochlorococcus marinus str. MIT 9313] |
| 3 | NO eubacterial outgroup | gi\|33862391\|ref\|NP_893951.1\| transcriptional regulator [Prochlorococcus marinus str. MIT 9313] |
| 3 | NO eubacterial outgroup | gi\|33862406\|ref\|NP_893966.1\| succinate dehydrogenase/fumarate reductase iron-sulfur subunit [Prochlorococcus marinus str. MIT 9313] |
| 3 | NO eubacterial outgroup | gi\|33862410\|ref\|NP_893970.1\| hypothetical protein PMT0137 [Prochlorococcus marinus str. MIT 9313] |
| 3 | NO eubacterial outgroup | gi\|33862427\|ref\|NP_893987.1\| protein kinase:Serine/threonine protein kinase [Prochlorococcus marinus str. MIT 9313] |
| 3 | NO eubacterial outgroup | gi\|33862437\|ref\|NP_893997.1\| 6-pyruvoyl-tetrahydropterin synthase [Prochlorococcus marinus str. MIT 9313] |
| 3 | NO eubacterial outgroup | gi\|33862445\|ref\|NP_894005.1\| hypothetical protein PMT0172 [Prochlorococcus marinus str. MIT 9313] |
| 3 | NO eubacterial outgroup | gi\|33862449\|ref\|NP_894009.1\| hypothetical protein PMT0176 [Prochlorococcus marinus str. MIT 9313] |
| 3 | NO eubacterial outgroup | gi\|33862452\|ref\|NP_894012.1\| hypothetical protein PMT0179 [Prochlorococcus marinus str. MIT 9313] |
| 3 | NO eubacterial outgroup | gi\|33862455\|ref\|NP_894015.1\| RND family outer membrane efflux protein [Prochlorococcus marinus str. MIT 9313] |
| 3 | NO eubacterial outgroup | gi\|33862465\|ref\|NP_894025.1\| glycoside hydrolase family protein [Prochlorococcus marinus str. MIT 9313] |
| 3 | NO eubacterial outgroup | gi\|33862477\|ref\|NP_894037.1\| cobalt ABC transporter permease [Prochlorococcus marinus str. MIT 9313] |
| 3 | NO eubacterial outgroup | gi\|33862479\|ref\|NP_894039.1\| hypothetical protein PMT0206 [Prochlorococcus marinus str. MIT 9313] |
| 3 | NO eubacterial outgroup | gi\|33862498\|ref\|NP_894058.1\| cystathionine gamma-synthase [Prochlorococcus marinus str. MIT 9313] |
| 3 | NO eubacterial outgroup | gi\|33862549\|ref\|NP_894109.1\| menaquinone biosynthesis methlytransferase, partial [Prochlorococcus marinus str. MIT 9313] |
| 3 | NO eubacterial outgroup | gi\|33862560\|ref\|NP_894120.1\| iron ABC transporter substrate-binding protein [Prochlorococcus marinus str. MIT 9313] |
| 3 | NO eubacterial outgroup | gi\|33862570\|ref\|NP_894130.1\| hypothetical protein PMT0297 [Prochlorococcus marinus str. MIT 9313] |
| 3 | NO eubacterial outgroup | gi\|33862580\|ref\|NP_894140.1\| hypothetical protein PMT0307 [Prochlorococcus marinus str. MIT 9313] |
| 3 | NO eubacterial outgroup | gi\|33862581\|ref\|NP_894141.1\| hydantoinase/oxoprolinase:hydantoinase B/oxoprolinase [Prochlorococcus marinus str. MIT 9313] |
| 3 | NO eubacterial outgroup | gi\|33862607\|ref\|NP_894167.1\| tRNA/rRNA methyltransferase SpoU [Prochlorococcus marinus str. MIT 9313] |
| 3 | NO eubacterial outgroup | gi\|33862629\|ref\|NP_894189.1\| bifuntional enzyme: tRNA methyltransferase; 2-C-methyl-D-erythritol 2,4-cyclodiphosphate synthase [Prochlorococcus marinus str. MIT 9313] |
| 3 | NO eubacterial outgroup | gi\|33862657\|ref\|NP_894217.1\| peptide deformylase [Prochlorococcus marinus str. MIT 9313] |
| 3 | NO eubacterial outgroup | gi\|33862661\|ref\|NP_894221.1\| hypothetical protein PMT0388 [Prochlorococcus marinus str. MIT 9313] |
| 3 | NO eubacterial outgroup | gi\|33862664\|ref\|NP_894224.1\| hypothetical protein PMT0391 [Prochlorococcus marinus str. MIT 9313] |
| 3 | NO eubacterial outgroup | gi\|33862669\|ref\|NP_894229.1\| 2-phosphosulfolactate phosphatase [Prochlorococcus marinus str. MIT 9313] |
| 3 | NO eubacterial outgroup | gi\|33862674\|ref\|NP_894234.1\| cytoplasmic peptidoglycan synthetase [Prochlorococcus marinus str. MIT 9313] |
| 3 | NO eubacterial outgroup | gi\|33862682\|ref\|NP_894242.1\| hypothetical protein PMT0409 [Prochlorococcus marinus str. MIT 9313] |
| 3 | NO eubacterial outgroup | gi\|33862691\|ref\|NP_894251.1\| ABC transporter substrate-binding protein [Prochlorococcus marinus str. MIT 9313] |
| 3 | NO eubacterial outgroup | gi\|33862715\|ref\|NP_894275.1\| hypothetical protein PMT0442 [Prochlorococcus marinus str. MIT 9313] |
| 3 | NO eubacterial outgroup | gi\|33862719\|ref\|NP_894279.1\| hypothetical protein PMT0446 [Prochlorococcus marinus str. MIT 9313] |
| 3 | NO eubacterial outgroup | gi\|33862737\|ref\|NP_894297.1\| hypothetical protein PMT0464 [Prochlorococcus marinus str. MIT 9313] |
| 3 | NO eubacterial outgroup | gi\|33862753\|ref\|NP_894313.1\| short-chain dehydrogenase/reductase [Prochlorococcus marinus str. MIT 9313] |
| 3 | NO eubacterial outgroup | gi\|33862769\|ref\|NP_894329.1\| light-harvesting complex protein [Prochlorococcus marinus str. MIT 9313] |
| 3 | NO eubacterial outgroup | gi\|33862779\|ref\|NP_894339.1\| NADH-flavin reductase [Prochlorococcus marinus str. MIT 9313] |
| 3 | NO eubacterial outgroup | gi\|33862782\|ref\|NP_894342.1\| cytochrome c, class IC:cytochrome c, class I [Prochlorococcus marinus str. MIT 9313] |
| 3 | NO eubacterial outgroup | gi\|33862799\|ref\|NP_894359.1\| hypothetical protein PMT0526 [Prochlorococcus marinus str. MIT 9313] |
| 3 | NO eubacterial outgroup | gi\|33862824\|ref\|NP_894384.1\| sarcosine-dimethylglycine methyltransferase [Prochlorococcus marinus str. MIT 9313] |
| 3 | NO eubacterial outgroup | gi\|33862830\|ref\|NP_894390.1\| hypothetical protein PMT0557 [Prochlorococcus marinus str. MIT 9313] |
| 3 | NO eubacterial outgroup | gi\|33862831\|ref\|NP_894391.1\| uracil phosphoribosyltransferase [Prochlorococcus marinus str. MIT 9313] |
| 3 | NO eubacterial outgroup | gi\|33862849\|ref\|NP_894409.1\| carboxypeptidase [Prochlorococcus marinus str. MIT 9313] |
| 3 | NO eubacterial outgroup | gi\|33862862\|ref\|NP_894422.1\| ABC transporter [Prochlorococcus marinus str. MIT 9313] |
| 3 | NO eubacterial outgroup | gi\|33862863\|ref\|NP_894423.1\| phycocyanobilin:ferredoxin oxidoreductase [Prochlorococcus marinus str. MIT 9313] |
| 3 | NO eubacterial outgroup | gi\|33862868\|ref\|NP_894428.1\| lipoprotein signal peptidase [Prochlorococcus marinus str. MIT 9313] |
| 3 | NO eubacterial outgroup | gi\|33862870\|ref\|NP_894430.1\| hypothetical protein PMT0597 [Prochlorococcus marinus str. MIT 9313] |
| 3 | NO eubacterial outgroup | gi\|33862879\|ref\|NP_894439.1\| hypothetical protein PMT0606 [Prochlorococcus marinus str. MIT 9313] |
| 3 | NO eubacterial outgroup | gi\|33862885\|ref\|NP_894445.1\| methyltransferase [Prochlorococcus marinus str. MIT 9313] |
| 3 | NO eubacterial outgroup | gi\|33862899\|ref\|NP_894459.1\| hypothetical protein PMT0626 [Prochlorococcus marinus str. MIT 9313] |
| 3 | NO eubacterial outgroup | gi\|33862911\|ref\|NP_894471.1\| UmuC protein [Prochlorococcus marinus str. MIT 9313] |
| 3 | NO eubacterial outgroup | gi\|33862925\|ref\|NP_894485.1\| STAS domain-containing protein [Prochlorococcus marinus str. MIT 9313] |
| 3 | NO eubacterial outgroup | gi\|33862937\|ref\|NP_894497.1\| hypothetical protein PMT0665 [Prochlorococcus marinus str. MIT 9313] |
| 3 | NO eubacterial outgroup | gi\|33862938\|ref\|NP_894498.1\| hypothetical protein PMT0666 [Prochlorococcus marinus str. MIT 9313] |
| 3 | NO eubacterial outgroup | gi\|33862944\|ref\|NP_894504.1\| ATP-dependent Clp protease adaptor [Prochlorococcus marinus str. MIT 9313] |
| 3 | NO eubacterial outgroup | gi\|33862947\|ref\|NP_894507.1\| Thf1-like protein [Prochlorococcus marinus str. MIT 9313] |
| 3 | NO eubacterial outgroup | gi\|33862955\|ref\|NP_894515.1\| hypothetical protein PMT0683 [Prochlorococcus marinus str. MIT 9313] |
| 3 | NO eubacterial outgroup | gi\|33862960\|ref\|NP_894520.1\| hypothetical protein PMT0688 [Prochlorococcus marinus str. MIT 9313] |
| 3 | NO eubacterial outgroup | gi\|33862961\|ref\|NP_894521.1\| hypothetical protein PMT0689 [Prochlorococcus marinus str. MIT 9313] |
| 3 | NO eubacterial outgroup | gi\|33862962\|ref\|NP_894522.1\| hypothetical protein PMT0690 [Prochlorococcus marinus str. MIT 9313] |
| 3 | NO eubacterial outgroup | gi\|33862964\|ref\|NP_894524.1\| sugar ABC transporter [Prochlorococcus marinus str. MIT 9313] |
| 3 | NO eubacterial outgroup | gi\|33862967\|ref\|NP_894527.1\| 50S ribosomal protein L28 [Prochlorococcus marinus str. MIT 9313] |
| 3 | NO eubacterial outgroup | gi\|33862976\|ref\|NP_894536.1\| DnaJ2 protein [Prochlorococcus marinus str. MIT 9313] |
| 3 | NO eubacterial outgroup | gi\|33862979\|ref\|NP_894539.1\| cyclophilin type peptidyl-prolyl cis-trans isomerase [Prochlorococcus marinus str. MIT 9313] |
| 3 | NO eubacterial outgroup | gi\|33862992\|ref\|NP_894552.1\| D-Ala-D-Ala carboxypeptidase 3 [Prochlorococcus marinus str. MIT 9313] |
| 3 | NO eubacterial outgroup | gi\|33863003\|ref\|NP_894563.1\| hypothetical protein PMT0731 [Prochlorococcus marinus str. MIT 9313] |
| 3 | NO eubacterial outgroup | gi\|33863007\|ref\|NP_894567.1\| hypothetical protein PMT0735 [Prochlorococcus marinus str. MIT 9313] |
| 3 | NO eubacterial outgroup | gi\|33863020\|ref\|NP_894580.1\| proton extrusion protein PcxA [Prochlorococcus marinus str. MIT 9313] |
| 3 | NO eubacterial outgroup | gi\|33863043\|ref\|NP_894603.1\| modulator of DNA gyrase [Prochlorococcus marinus str. MIT 9313] |
| 3 | NO eubacterial outgroup | gi\|33863056\|ref\|NP_894616.1\| type III sigma factor [Prochlorococcus marinus str. MIT 9313] |
| 3 | NO eubacterial outgroup | gi\|33863057\|ref\|NP_894617.1\| hypothetical protein PMT0785 [Prochlorococcus marinus str. MIT 9313] |
| 3 | NO eubacterial outgroup | gi\|33863059\|ref\|NP_894619.1\| hypothetical protein PMT0787 [Prochlorococcus marinus str. MIT 9313] |
| 3 | NO eubacterial outgroup | gi\|33863065\|ref\|NP_894625.1\| ribonucleotide reductase (class II) [Prochlorococcus marinus str. MIT 9313] |
| 3 | NO eubacterial outgroup | gi\|33863075\|ref\|NP_894635.1\| hypothetical protein PMT0803 [Prochlorococcus marinus str. MIT 9313] |
| 3 | NO eubacterial outgroup | gi\|33863093\|ref\|NP_894653.1\| hypothetical protein PMT0821 [Prochlorococcus marinus str. MIT 9313] |
| 3 | NO eubacterial outgroup | gi\|33863126\|ref\|NP_894686.1\| MscS family mechanosensitive ion channel [Prochlorococcus marinus str. MIT 9313] |
| 3 | NO eubacterial outgroup | gi\|33863133\|ref\|NP_894693.1\| hypothetical protein PMT0861 [Prochlorococcus marinus str. MIT 9313] |
| 3 | NO eubacterial outgroup | gi\|33863149\|ref\|NP_894709.1\| hypothetical protein PMT0877 [Prochlorococcus marinus str. MIT 9313] |
| 3 | NO eubacterial outgroup | gi\|33863152\|ref\|NP_894712.1\| hypothetical protein PMT0880 [Prochlorococcus marinus str. MIT 9313] |
| 3 | NO eubacterial outgroup | gi\|33863171\|ref\|NP_894731.1\| sulfate transporter [Prochlorococcus marinus str. MIT 9313] |
| 3 | NO eubacterial outgroup | gi\|33863187\|ref\|NP_894747.1\| outer membrane protein [Prochlorococcus marinus str. MIT 9313] |
| 3 | NO eubacterial outgroup | gi\|33863282\|ref\|NP_894842.1\| trigger factor [Prochlorococcus marinus str. MIT 9313] |
| 3 | NO eubacterial outgroup | gi\|33863315\|ref\|NP_894875.1\| major facilitator superfamily proline/betaine transporter [Prochlorococcus marinus str. MIT 9313] |
| 3 | NO eubacterial outgroup | gi\|33863317\|ref\|NP_894877.1\| light-harvesting complex protein [Prochlorococcus marinus str. MIT 9313] |
| 3 | NO eubacterial outgroup | gi\|33863324\|ref\|NP_894884.1\| hypothetical protein PMT1053 [Prochlorococcus marinus str. MIT 9313] |
| 3 | NO eubacterial outgroup | gi\|33863334\|ref\|NP_894894.1\| alpha/beta fold family hydrolase [Prochlorococcus marinus str. MIT 9313] |
| 3 | NO eubacterial outgroup | gi\|33863337\|ref\|NP_894897.1\| Orn/Lys/Arg decarboxylase family protein [Prochlorococcus marinus str. MIT 9313] |
| 3 | NO eubacterial outgroup | gi\|33863343\|ref\|NP_894903.1\| LytR-membrane bound transcriptional regulator [Prochlorococcus marinus str. MIT 9313] |
| 3 | NO eubacterial outgroup | gi\|33863357\|ref\|NP_894917.1\| UvrD/REP helicase [Prochlorococcus marinus str. MIT 9313] |
| 3 | NO eubacterial outgroup | gi\|33863365\|ref\|NP_894925.1\| BolA-like protein [Prochlorococcus marinus str. MIT 9313] |
| 3 | NO eubacterial outgroup | gi\|33863374\|ref\|NP_894934.1\| glucose 6-phosphate dehydrogenase effector OpcA [Prochlorococcus marinus str. MIT 9313] |
| 3 | NO eubacterial outgroup | gi\|33863404\|ref\|NP_894964.1\| beta-lactamase [Prochlorococcus marinus str. MIT 9313] |
| 3 | NO eubacterial outgroup | gi\|33863405\|ref\|NP_894965.1\| hypothetical protein PMT1134 [Prochlorococcus marinus str. MIT 9313] |
| 3 | NO eubacterial outgroup | gi\|33863413\|ref\|NP_894973.1\| hypothetical protein PMT1142 [Prochlorococcus marinus str. MIT 9313] |
| 3 | NO eubacterial outgroup | gi\|33863443\|ref\|NP_895003.1\| chaperon-like protein for quinone binding in photosystem II [Prochlorococcus marinus str. MIT 9313] |
| 3 | NO eubacterial outgroup | gi\|33863445\|ref\|NP_895005.1\| alpha/beta hydrolase superfamily protein [Prochlorococcus marinus str. MIT 9313] |
| 3 | NO eubacterial outgroup | gi\|33863447\|ref\|NP_895007.1\| cyclophilin-type peptidyl-prolyl cis-trans isomerase [Prochlorococcus marinus str. MIT 9313] |
| 3 | NO eubacterial outgroup | gi\|33863448\|ref\|NP_895008.1\| photosystem I assembly protein Ycf4 [Prochlorococcus marinus str. MIT 9313] |
| 3 | NO eubacterial outgroup | gi\|33863449\|ref\|NP_895009.1\| photosystem II PsbD protein (D2) [Prochlorococcus marinus str. MIT 9313] |
| 3 | NO eubacterial outgroup | gi\|33863450\|ref\|NP_895010.1\| photosystem II PsbC protein (CP43) [Prochlorococcus marinus str. MIT 9313] |
| 3 | NO eubacterial outgroup | gi\|33863461\|ref\|NP_895021.1\| acetyltransferase [Prochlorococcus marinus str. MIT 9313] |
| 3 | NO eubacterial outgroup | gi\|33863482\|ref\|NP_895042.1\| sodium-dependent bicarbonate transporter [Prochlorococcus marinus str. MIT 9313] |
| 3 | NO eubacterial outgroup | gi\|33863494\|ref\|NP_895054.1\| phosphoribosylanthranilate isomerase [Prochlorococcus marinus str. MIT 9313] |
| 3 | NO eubacterial outgroup | gi\|33863513\|ref\|NP_895073.1\| ribosome recycling factor [Prochlorococcus marinus str. MIT 9313] |
| 3 | NO eubacterial outgroup | gi\|33863522\|ref\|NP_895082.1\| Signal peptidase I [Prochlorococcus marinus str. MIT 9313] |
| 3 | NO eubacterial outgroup | gi\|33863523\|ref\|NP_895083.1\| hypothetical protein PMT1255 [Prochlorococcus marinus str. MIT 9313] |
| 3 | NO eubacterial outgroup | gi\|33863542\|ref\|NP_895102.1\| hypothetical protein PMT1274 [Prochlorococcus marinus str. MIT 9313] |
| 3 | NO eubacterial outgroup | gi\|33863551\|ref\|NP_895111.1\| pterin-4-alpha-carbinolamine dehydratase [Prochlorococcus marinus str. MIT 9313] |
| 3 | NO eubacterial outgroup | gi\|33863560\|ref\|NP_895120.1\| hypothetical protein PMT1292 [Prochlorococcus marinus str. MIT 9313] |
| 3 | NO eubacterial outgroup | gi\|33863570\|ref\|NP_895130.1\| BioD-like N-terminal domain of phosphotransacetylase, partial [Prochlorococcus marinus str. MIT 9313] |
| 3 | NO eubacterial outgroup | gi\|109150048\|ref\|YP_654194.1\| guanylate kinase [Prochlorococcus marinus str. MIT 9313] |
| 3 | NO eubacterial outgroup | gi\|33863590\|ref\|NP_895150.1\| apocytochrome f [Prochlorococcus marinus str. MIT 9313] |
| 3 | NO eubacterial outgroup | gi\|33863592\|ref\|NP_895152.1\| precorrin-4 C(11)-methyltransferase [Prochlorococcus marinus str. MIT 9313] |
| 3 | NO eubacterial outgroup | gi\|33863595\|ref\|NP_895155.1\| prenyltransferase [Prochlorococcus marinus str. MIT 9313] |
| 3 | NO eubacterial outgroup | gi\|33863599\|ref\|NP_895159.1\| VIC family potassium channel protein [Prochlorococcus marinus str. MIT 9313] |
| 3 | NO eubacterial outgroup | gi\|33863607\|ref\|NP_895167.1\| hypothetical protein PMT1340 [Prochlorococcus marinus str. MIT 9313] |
| 3 | NO eubacterial outgroup | gi\|33863611\|ref\|NP_895171.1\| hypothetical protein PMT1344 [Prochlorococcus marinus str. MIT 9313] |
| 3 | NO eubacterial outgroup | gi\|33863614\|ref\|NP_895174.1\| hypothetical protein PMT1347 [Prochlorococcus marinus str. MIT 9313] |
| 3 | NO eubacterial outgroup | gi\|33863617\|ref\|NP_895177.1\| hypothetical protein PMT1350 [Prochlorococcus marinus str. MIT 9313] |
| 3 | NO eubacterial outgroup | gi\|33863620\|ref\|NP_895180.1\| DNA topoisomerase I [Prochlorococcus marinus str. MIT 9313] |
| 3 | NO eubacterial outgroup | gi\|33863623\|ref\|NP_895183.1\| response regulator receiver domain-containing protein [Prochlorococcus marinus str. MIT 9313] |
| 3 | NO eubacterial outgroup | gi\|33863626\|ref\|NP_895186.1\| biotin--acetyl-CoA-carboxylase ligase [Prochlorococcus marinus str. MIT 9313] |
| 3 | NO eubacterial outgroup | gi\|33863630\|ref\|NP_895190.1\| hypothetical protein PMT1363 [Prochlorococcus marinus str. MIT 9313] |
| 3 | NO eubacterial outgroup | gi\|33863631\|ref\|NP_895191.1\| SMR family transporter PecM [Prochlorococcus marinus str. MIT 9313] |
| 3 | NO eubacterial outgroup | gi\|33863633\|ref\|NP_895193.1\| chorismate mutase [Prochlorococcus marinus str. MIT 9313] |
| 3 | NO eubacterial outgroup | gi\|33863637\|ref\|NP_895197.1\| hypothetical protein PMT1370 [Prochlorococcus marinus str. MIT 9313] |
| 3 | NO eubacterial outgroup | gi\|33863643\|ref\|NP_895203.1\| 30S ribosomal protein S14 [Prochlorococcus marinus str. MIT 9313] |
| 3 | NO eubacterial outgroup | gi\|33863671\|ref\|NP_895231.1\| hypothetical protein PMT1404 [Prochlorococcus marinus str. MIT 9313] |
| 3 | NO eubacterial outgroup | gi\|33863672\|ref\|NP_895232.1\| DNA-binding response regulator [Prochlorococcus marinus str. MIT 9313] |
| 3 | NO eubacterial outgroup | gi\|33863680\|ref\|NP_895240.1\| chloroplast outer envelope membrane protein [Prochlorococcus marinus str. MIT 9313] |
| 3 | NO eubacterial outgroup | gi\|33863694\|ref\|NP_895254.1\| cytochrome c-550 [Prochlorococcus marinus str. MIT 9313] |
| 3 | NO eubacterial outgroup | gi\|33863700\|ref\|NP_895260.1\| hypothetical protein PMT1433 [Prochlorococcus marinus str. MIT 9313] |
| 3 | NO eubacterial outgroup | gi\|33863711\|ref\|NP_895271.1\| HSP70 family molecular chaperone [Prochlorococcus marinus str. MIT 9313] |
| 3 | NO eubacterial outgroup | gi\|33863728\|ref\|NP_895288.1\| carbon-nitrogen hydrolase:NAD+ synthase [Prochlorococcus marinus str. MIT 9313] |
| 3 | NO eubacterial outgroup | gi\|33863755\|ref\|NP_895315.1\| 8-amino-7-oxononanoate synthase [Prochlorococcus marinus str. MIT 9313] |
| 3 | NO eubacterial outgroup | gi\|33863764\|ref\|NP_895324.1\| aldo/keto reductase [Prochlorococcus marinus str. MIT 9313] |
| 3 | NO eubacterial outgroup | gi\|33863766\|ref\|NP_895326.1\| hypothetical protein PMT1499 [Prochlorococcus marinus str. MIT 9313] |
| 3 | NO eubacterial outgroup | gi\|33863768\|ref\|NP_895328.1\| hypothetical protein PMT1501 [Prochlorococcus marinus str. MIT 9313] |
| 3 | NO eubacterial outgroup | gi\|33863798\|ref\|NP_895358.1\| hypothetical protein PMT1531 [Prochlorococcus marinus str. MIT 9313] |
| 3 | NO eubacterial outgroup | gi\|33863816\|ref\|NP_895376.1\| hypothetical protein PMT1549 [Prochlorococcus marinus str. MIT 9313] |
| 3 | NO eubacterial outgroup | gi\|33863818\|ref\|NP_895378.1\| FAD-dependent pyridine nucleotide-disulfide oxidoreductase [Prochlorococcus marinus str. MIT 9313] |
| 3 | NO eubacterial outgroup | gi\|33863821\|ref\|NP_895381.1\| pentapeptide repeat-containing protein [Prochlorococcus marinus str. MIT 9313] |
| 3 | NO eubacterial outgroup | gi\|33863825\|ref\|NP_895385.1\| NAD binding site [Prochlorococcus marinus str. MIT 9313] |
| 3 | NO eubacterial outgroup | gi\|33863840\|ref\|NP_895400.1\| ABC transporter [Prochlorococcus marinus str. MIT 9313] |
| 3 | NO eubacterial outgroup | gi\|33863841\|ref\|NP_895401.1\| membrane permease or ABC transporter component [Prochlorococcus marinus str. MIT 9313] |
| 3 | NO eubacterial outgroup | gi\|33863842\|ref\|NP_895402.1\| membrane permease or ABC transporter component [Prochlorococcus marinus str. MIT 9313] |
| 3 | NO eubacterial outgroup | gi\|33863881\|ref\|NP_895441.1\| phosphoglucomutase [Prochlorococcus marinus str. MIT 9313] |
| 3 | NO eubacterial outgroup | gi\|33863891\|ref\|NP_895451.1\| phosphoadenosine phosphosulfate reductase [Prochlorococcus marinus str. MIT 9313] |
| 3 | NO eubacterial outgroup | gi\|33863916\|ref\|NP_895476.1\| cytochrome b6 [Prochlorococcus marinus str. MIT 9313] |
| 3 | NO eubacterial outgroup | gi\|33863921\|ref\|NP_895481.1\| cell division topological specificity factor MinE [Prochlorococcus marinus str. MIT 9313] |
| 3 | NO eubacterial outgroup | gi\|33863931\|ref\|NP_895491.1\| ferredoxin [Prochlorococcus marinus str. MIT 9313] |
| 3 | NO eubacterial outgroup | gi\|33863932\|ref\|NP_895492.1\| photosystem II PsbB protein (CP47) [Prochlorococcus marinus str. MIT 9313] |
| 3 | NO eubacterial outgroup | gi\|33863944\|ref\|NP_895504.1\| CpeS-like protein [Prochlorococcus marinus str. MIT 9313] |
| 3 | NO eubacterial outgroup | gi\|33863945\|ref\|NP_895505.1\| CpeT protein [Prochlorococcus marinus str. MIT 9313] |
| 3 | NO eubacterial outgroup | gi\|33863949\|ref\|NP_895509.1\| phycobilisome protein (phycoerythrin, alpha-subunit) [Prochlorococcus marinus str. MIT 9313] |
| 3 | NO eubacterial outgroup | gi\|33863950\|ref\|NP_895510.1\| phycobilisome protein (phycoerythrin beta-subunit) [Prochlorococcus marinus str. MIT 9313] |
| 3 | NO eubacterial outgroup | gi\|33863957\|ref\|NP_895517.1\| rare lipoprotein A [Prochlorococcus marinus str. MIT 9313] |
| 3 | NO eubacterial outgroup | gi\|33863968\|ref\|NP_895528.1\| hypothetical protein PMT1701 [Prochlorococcus marinus str. MIT 9313] |
| 3 | NO eubacterial outgroup | gi\|33863977\|ref\|NP_895537.1\| photosystem I protein PsaD [Prochlorococcus marinus str. MIT 9313] |
| 3 | NO eubacterial outgroup | gi\|33863979\|ref\|NP_895539.1\| hypothetical protein PMT1712 [Prochlorococcus marinus str. MIT 9313] |
| 3 | NO eubacterial outgroup | gi\|33863983\|ref\|NP_895543.1\| S-adenosylmethionine decarboxylase [Prochlorococcus marinus str. MIT 9313] |
| 3 | NO eubacterial outgroup | gi\|33863996\|ref\|NP_895556.1\| ATPase AAA [Prochlorococcus marinus str. MIT 9313] |
| 3 | NO eubacterial outgroup | gi\|33863997\|ref\|NP_895557.1\| hypothetical protein PMT1730 [Prochlorococcus marinus str. MIT 9313] |
| 3 | NO eubacterial outgroup | gi\|33863998\|ref\|NP_895558.1\| NADH dehydrogenase I subunit N [Prochlorococcus marinus str. MIT 9313] |
| 3 | NO eubacterial outgroup | gi\|33864006\|ref\|NP_895566.1\| 50S ribosomal protein L16 [Prochlorococcus marinus str. MIT 9313] |
| 3 | NO eubacterial outgroup | gi\|33864023\|ref\|NP_895583.1\| 50S ribosomal protein L17 [Prochlorococcus marinus str. MIT 9313] |
| 3 | NO eubacterial outgroup | gi\|33864025\|ref\|NP_895585.1\| 50S ribosomal protein L13 [Prochlorococcus marinus str. MIT 9313] |
| 3 | NO eubacterial outgroup | gi\|33864026\|ref\|NP_895586.1\| 30S ribosomal protein S9 [Prochlorococcus marinus str. MIT 9313] |
| 3 | NO eubacterial outgroup | gi\|33864027\|ref\|NP_895587.1\| 50S ribosomal protein L31 [Prochlorococcus marinus str. MIT 9313] |
| 3 | NO eubacterial outgroup | gi\|33864032\|ref\|NP_895592.1\| hypothetical protein PMT1765 [Prochlorococcus marinus str. MIT 9313] |
| 3 | NO eubacterial outgroup | gi\|33864035\|ref\|NP_895595.1\| photosystem I reaction center protein subunit XI [Prochlorococcus marinus str. MIT 9313] |
| 3 | NO eubacterial outgroup | gi\|33864036\|ref\|NP_895596.1\| photosystem I P700 chlorophyll a apoprotein A2 [Prochlorococcus marinus str. MIT 9313] |
| 3 | NO eubacterial outgroup | gi\|33864037\|ref\|NP_895597.1\| photosystem I P700 chlorophyll a apoprotein A1 [Prochlorococcus marinus str. MIT 9313] |
| 3 | NO eubacterial outgroup | gi\|33864046\|ref\|NP_895606.1\| 30S ribosomal protein S12 [Prochlorococcus marinus str. MIT 9313] |
| 3 | NO eubacterial outgroup | gi\|33864047\|ref\|NP_895607.1\| 30S ribosomal protein S7 [Prochlorococcus marinus str. MIT 9313] |
| 3 | NO eubacterial outgroup | gi\|33864051\|ref\|NP_895611.1\| ATP-dependent protease La [Prochlorococcus marinus str. MIT 9313] |
| 3 | NO eubacterial outgroup | gi\|33864059\|ref\|NP_895619.1\| ATP-dependent Clp protease adaptor protein ClpS [Prochlorococcus marinus str. MIT 9313] |
| 3 | NO eubacterial outgroup | gi\|33864067\|ref\|NP_895627.1\| photosystem II manganese-stabilizing protein [Prochlorococcus marinus str. MIT 9313] |
| 3 | NO eubacterial outgroup | gi\|33864073\|ref\|NP_895633.1\| NAD binding site:FAD-dependent pyridine nucleotide-disulfide [Prochlorococcus marinus str. MIT 9313] |
| 3 | NO eubacterial outgroup | gi\|33864090\|ref\|NP_895650.1\| phosphoglucosamine mutase [Prochlorococcus marinus str. MIT 9313] |
| 3 | NO eubacterial outgroup | gi\|33864093\|ref\|NP_895653.1\| thioredoxin [Prochlorococcus marinus str. MIT 9313] |
| 3 | NO eubacterial outgroup | gi\|33864096\|ref\|NP_895656.1\| cob(I)alamin adenosyltransferase [Prochlorococcus marinus str. MIT 9313] |
| 3 | NO eubacterial outgroup | gi\|33864103\|ref\|NP_895663.1\| twin arginine translocase protein A [Prochlorococcus marinus str. MIT 9313] |
| 3 | NO eubacterial outgroup | gi\|33864109\|ref\|NP_895669.1\| pentapeptide repeat-containing protein [Prochlorococcus marinus str. MIT 9313] |
| 3 | NO eubacterial outgroup | gi\|33864111\|ref\|NP_895671.1\| isopropylmalate isomerase large subunit [Prochlorococcus marinus str. MIT 9313] |
| 3 | NO eubacterial outgroup | gi\|33864114\|ref\|NP_895674.1\| serine hydroxymethyltransferase [Prochlorococcus marinus str. MIT 9313] |
| 3 | NO eubacterial outgroup | gi\|33864150\|ref\|NP_895710.1\| pleiotropic regulatory protein [Prochlorococcus marinus str. MIT 9313] |
| 3 | NO eubacterial outgroup | gi\|33864186\|ref\|NP_895746.1\| group 1 glycosyl transferase [Prochlorococcus marinus str. MIT 9313] |
| 3 | NO eubacterial outgroup | gi\|33864203\|ref\|NP_895763.1\| hypothetical protein PMT1938 [Prochlorococcus marinus str. MIT 9313] |
| 3 | NO eubacterial outgroup | gi\|33864209\|ref\|NP_895769.1\| CPA2 family Na+/H+ antiporter [Prochlorococcus marinus str. MIT 9313] |
| 3 | NO eubacterial outgroup | gi\|33864220\|ref\|NP_895780.1\| acyl carrier protein [Prochlorococcus marinus str. MIT 9313] |
| 3 | NO eubacterial outgroup | gi\|33864230\|ref\|NP_895790.1\| TPR repeat-containing glycosyl transferase [Prochlorococcus marinus str. MIT 9313] |
| 3 | NO eubacterial outgroup | gi\|33864232\|ref\|NP_895792.1\| integral membrane protein [Prochlorococcus marinus str. MIT 9313] |
| 3 | NO eubacterial outgroup | gi\|33864235\|ref\|NP_895795.1\| hypothetical protein PMT1970 [Prochlorococcus marinus str. MIT 9313] |
| 3 | NO eubacterial outgroup | gi\|33864242\|ref\|NP_895802.1\| heat shock protein DnaJ [Prochlorococcus marinus str. MIT 9313] |
| 3 | NO eubacterial outgroup | gi\|33864244\|ref\|NP_895804.1\| porin [Prochlorococcus marinus str. MIT 9313] |
| 3 | NO eubacterial outgroup | gi\|33864257\|ref\|NP_895817.1\| photosystem I assembly protein Ycf3 [Prochlorococcus marinus str. MIT 9313] |
| 3 | NO eubacterial outgroup | gi\|33864265\|ref\|NP_895825.1\| hypothetical protein PMT2000 [Prochlorococcus marinus str. MIT 9313] |
| 3 | NO eubacterial outgroup | gi\|33864266\|ref\|NP_895826.1\| poly A polymerase [Prochlorococcus marinus str. MIT 9313] |
| 3 | NO eubacterial outgroup | gi\|33864267\|ref\|NP_895827.1\| RNA recognition motif-containing protein [Prochlorococcus marinus str. MIT 9313] |
| 3 | NO eubacterial outgroup | gi\|33864268\|ref\|NP_895828.1\| squalene and phytoene synthase [Prochlorococcus marinus str. MIT 9313] |
| 3 | NO eubacterial outgroup | gi\|33864270\|ref\|NP_895830.1\| NADH dehydrogenase I subunit M [Prochlorococcus marinus str. MIT 9313] |
| 3 | NO eubacterial outgroup | gi\|33864324\|ref\|NP_895884.1\| 1,4-dihydroxy-2-naphthoate octaprenyltransferase [Prochlorococcus marinus str. MIT 9313] |
| 3 | NO eubacterial outgroup | gi\|33864327\|ref\|NP_895887.1\| glutaredoxin [Prochlorococcus marinus str. MIT 9313] |
| 3 | NO eubacterial outgroup | gi\|33864356\|ref\|NP_895916.1\| hypothetical protein PMT2092 [Prochlorococcus marinus str. MIT 9313] |
| 3 | NO eubacterial outgroup | gi\|33864410\|ref\|NP_895970.1\| adenylate cyclase [Prochlorococcus marinus str. MIT 9313] |
| 3 | NO eubacterial outgroup | gi\|33864465\|ref\|NP_896025.1\| ABC transporter [Prochlorococcus marinus str. MIT 9313] |
| 3 | NO eubacterial outgroup | gi\|33864482\|ref\|NP_896042.1\| Dps family protein [Prochlorococcus marinus str. MIT 9313] |
| 3 | NO eubacterial outgroup | gi\|33864511\|ref\|NP_896071.1\| hypothetical protein PMT2247 [Prochlorococcus marinus str. MIT 9313] |
| 3 | NO eubacterial outgroup | gi\|33864518\|ref\|NP_896078.1\| NAD binding site:D-amino acid oxidase [Prochlorococcus marinus str. MIT 9313] |
| 3 | NO eubacterial outgroup | gi\|33864533\|ref\|NP_896093.1\| esterase/lipase/thioesterase family protein [Prochlorococcus marinus str. MIT 9313] |
| 2 | PSC clade basal to Gloobacter | gi\|33862284\|ref\|NP_893844.1\| signal recognition particle protein [Prochlorococcus marinus str. MIT 9313] |
| 2 | PSC clade basal to Gloobacter | gi\|33862285\|ref\|NP_893845.1\| protein phosphatase 2C domain-containing protein [Prochlorococcus marinus str. MIT 9313] |
| 2 | PSC clade basal to Gloobacter | gi\|33862287\|ref\|NP_893847.1\| RNA recognition motif-containing protein [Prochlorococcus marinus str. MIT 9313] |
| 2 | PSC clade basal to Gloobacter | gi\|33862302\|ref\|NP_893862.1\| thiamine monophosphate kinase [Prochlorococcus marinus str. MIT 9313] |
| 2 | PSC clade basal to Gloobacter | gi\|33862305\|ref\|NP_893865.1\| acetyl-CoA biotin carboxyl carrier subunit [Prochlorococcus marinus str. MIT 9313] |
| 2 | PSC clade basal to Gloobacter | gi\|33862317\|ref\|NP_893877.1\| cobalt-precorrin-6A synthase [Prochlorococcus marinus str. MIT 9313] |
| 2 | PSC clade basal to Gloobacter | gi\|33862318\|ref\|NP_893878.1\| GMP synthase [Prochlorococcus marinus str. MIT 9313] |
| 2 | PSC clade basal to Gloobacter | gi\|33862328\|ref\|NP_893888.1\| 50S ribosomal protein L20 [Prochlorococcus marinus str. MIT 9313] |
| 2 | PSC clade basal to Gloobacter | gi\|33862330\|ref\|NP_893890.1\| amidase [Prochlorococcus marinus str. MIT 9313] |
| 2 | PSC clade basal to Gloobacter | gi\|33862331\|ref\|NP_893891.1\| glycosyl transferase family protein [Prochlorococcus marinus str. MIT 9313] |
| 2 | PSC clade basal to Gloobacter | gi\|33862337\|ref\|NP_893897.1\| aspartate semialdehyde dehydrogenase [Prochlorococcus marinus str. MIT 9313] |
| 2 | PSC clade basal to Gloobacter | gi\|33862341\|ref\|NP_893901.1\| hypothetical protein PMT0068 [Prochlorococcus marinus str. MIT 9313] |
| 2 | PSC clade basal to Gloobacter | gi\|33862346\|ref\|NP_893906.1\| aspartate kinase [Prochlorococcus marinus str. MIT 9313] |
| 2 | PSC clade basal to Gloobacter | gi\|33862347\|ref\|NP_893907.1\| DNA polymerase III subunit delta [Prochlorococcus marinus str. MIT 9313] |
| 2 | PSC clade basal to Gloobacter | gi\|33862348\|ref\|NP_893908.1\| precorrin-8X methylmutase CobH [Prochlorococcus marinus str. MIT 9313] |
| 2 | PSC clade basal to Gloobacter | gi\|33862387\|ref\|NP_893947.1\| dTDP-4-dehydrorhamnose reductase [Prochlorococcus marinus str. MIT 9313] |
| 2 | PSC clade basal to Gloobacter | gi\|33862401\|ref\|NP_893961.1\| alpha/beta fold family hydrolase [Prochlorococcus marinus str. MIT 9313] |
| 2 | PSC clade basal to Gloobacter | gi\|33862403\|ref\|NP_893963.1\| hypothetical protein PMT0130 [Prochlorococcus marinus str. MIT 9313] |
| 2 | PSC clade basal to Gloobacter | gi\|33862409\|ref\|NP_893969.1\| carbohydrate kinase [Prochlorococcus marinus str. MIT 9313] |
| 2 | PSC clade basal to Gloobacter | gi\|33862425\|ref\|NP_893985.1\| hypothetical protein PMT0152 [Prochlorococcus marinus str. MIT 9313] |
| 2 | PSC clade basal to Gloobacter | gi\|33862431\|ref\|NP_893991.1\| zinc metallopeptidase [Prochlorococcus marinus str. MIT 9313] |
| 2 | PSC clade basal to Gloobacter | gi\|33862448\|ref\|NP_894008.1\| Fe-S oxidoreductase [Prochlorococcus marinus str. MIT 9313] |
| 2 | PSC clade basal to Gloobacter | gi\|33862471\|ref\|NP_894031.1\| precorrin-2 C20-methyltransferase [Prochlorococcus marinus str. MIT 9313] |
| 2 | PSC clade basal to Gloobacter | gi\|33862473\|ref\|NP_894033.1\| nitrogen regulation protein NifR3 family protein [Prochlorococcus marinus str. MIT 9313] |
| 2 | PSC clade basal to Gloobacter | gi\|33862482\|ref\|NP_894042.1\| group 1 glycosyl transferase [Prochlorococcus marinus str. MIT 9313] |
| 2 | PSC clade basal to Gloobacter | gi\|33862483\|ref\|NP_894043.1\| MFS superfamily transporter [Prochlorococcus marinus str. MIT 9313] |
| 2 | PSC clade basal to Gloobacter | gi\|33862486\|ref\|NP_894046.1\| light repressed protein A-like protein [Prochlorococcus marinus str. MIT 9313] |
| 2 | PSC clade basal to Gloobacter | gi\|33862496\|ref\|NP_894056.1\| S-adenosylmethionine--tRNA ribosyltransferase-isomerase [Prochlorococcus marinus str. MIT 9313] |
| 2 | PSC clade basal to Gloobacter | gi\|33862497\|ref\|NP_894057.1\| O-acetylserine (thiol)-lyase A [Prochlorococcus marinus str. MIT 9313] |
| 2 | PSC clade basal to Gloobacter | gi\|33862527\|ref\|NP_894087.1\| NifU-like protein [Prochlorococcus marinus str. MIT 9313] |
| 2 | PSC clade basal to Gloobacter | gi\|33862548\|ref\|NP_894108.1\| SAM-binding motif-containing protein [Prochlorococcus marinus str. MIT 9313] |
| 2 | PSC clade basal to Gloobacter | gi\|33862590\|ref\|NP_894150.1\| oxygen-independent coproporphyrinogen III oxidase [Prochlorococcus marinus str. MIT 9313] |
| 2 | PSC clade basal to Gloobacter | gi\|33862592\|ref\|NP_894152.1\| cell division protein FtsZ [Prochlorococcus marinus str. MIT 9313] |
| 2 | PSC clade basal to Gloobacter | gi\|33862599\|ref\|NP_894159.1\| cytosine deaminase [Prochlorococcus marinus str. MIT 9313] |
| 2 | PSC clade basal to Gloobacter | gi\|33862605\|ref\|NP_894165.1\| UDP-N-acetylglucosamine 1-carboxyvinyltransferase [Prochlorococcus marinus str. MIT 9313] |
| 2 | PSC clade basal to Gloobacter | gi\|161350046\|ref\|NP_894177.2\| tRNA-specific 2-thiouridylase MnmA [Prochlorococcus marinus str. MIT 9313] |
| 2 | PSC clade basal to Gloobacter | gi\|33862622\|ref\|NP_894182.1\| signal recognition particle protein (SRP54) [Prochlorococcus marinus str. MIT 9313] |
| 2 | PSC clade basal to Gloobacter | gi\|33862631\|ref\|NP_894191.1\| circadian phase modifier CpmA-like protein [Prochlorococcus marinus str. MIT 9313] |
| 2 | PSC clade basal to Gloobacter | gi\|33862636\|ref\|NP_894196.1\| thiamine-phosphate pyrophosphorylase [Prochlorococcus marinus str. MIT 9313] |
| 2 | PSC clade basal to Gloobacter | gi\|33862640\|ref\|NP_894200.1\| phenylalanyl-tRNA synthetase subunit alpha [Prochlorococcus marinus str. MIT 9313] |
| 2 | PSC clade basal to Gloobacter | gi\|33862641\|ref\|NP_894201.1\| inorganic polyphosphate/ATP-NAD kinase [Prochlorococcus marinus str. MIT 9313] |
| 2 | PSC clade basal to Gloobacter | gi\|33862653\|ref\|NP_894213.1\| SOS function regulatory protein, LexA repressor [Prochlorococcus marinus str. MIT 9313] |
| 2 | PSC clade basal to Gloobacter | gi\|33862665\|ref\|NP_894225.1\| polyprenyl synthetase; solanesyl diphosphate synthase (sds) [Prochlorococcus marinus str. MIT 9313] |
| 2 | PSC clade basal to Gloobacter | gi\|33862671\|ref\|NP_894231.1\| 3-phosphoshikimate 1-carboxyvinyltransferase [Prochlorococcus marinus str. MIT 9313] |
| 2 | PSC clade basal to Gloobacter | gi\|33862676\|ref\|NP_894236.1\| glycogen synthase [Prochlorococcus marinus str. MIT 9313] |
| 2 | PSC clade basal to Gloobacter | gi\|33862680\|ref\|NP_894240.1\| leader peptidase I [Prochlorococcus marinus str. MIT 9313] |
| 2 | PSC clade basal to Gloobacter | gi\|33862689\|ref\|NP_894249.1\| ABC transporter [Prochlorococcus marinus str. MIT 9313] |
| 2 | PSC clade basal to Gloobacter | gi\|33862699\|ref\|NP_894259.1\| homoserine kinase [Prochlorococcus marinus str. MIT 9313] |
| 2 | PSC clade basal to Gloobacter | gi\|33862781\|ref\|NP_894341.1\| ABC transporter substrate-binding protein, phosphate [Prochlorococcus marinus str. MIT 9313] |
| 2 | PSC clade basal to Gloobacter | gi\|33862792\|ref\|NP_894352.1\| mRNA binding protein [Prochlorococcus marinus str. MIT 9313] |
| 2 | PSC clade basal to Gloobacter | gi\|33862804\|ref\|NP_894364.1\| 3-isopropylmalate dehydrogenase [Prochlorococcus marinus str. MIT 9313] |
| 2 | PSC clade basal to Gloobacter | gi\|33862806\|ref\|NP_894366.1\| type 4 prepilin peptidase [Prochlorococcus marinus str. MIT 9313] |
| 2 | PSC clade basal to Gloobacter | gi\|33862811\|ref\|NP_894371.1\| hypothetical protein PMT0538 [Prochlorococcus marinus str. MIT 9313] |
| 2 | PSC clade basal to Gloobacter | gi\|33862821\|ref\|NP_894381.1\| short-chain dehydrogenase/reductase [Prochlorococcus marinus str. MIT 9313] |
| 2 | PSC clade basal to Gloobacter | gi\|33862838\|ref\|NP_894398.1\| 6-phosphogluconate dehydrogenase [Prochlorococcus marinus str. MIT 9313] |
| 2 | PSC clade basal to Gloobacter | gi\|33862843\|ref\|NP_894403.1\| heme transporter [Prochlorococcus marinus str. MIT 9313] |
| 2 | PSC clade basal to Gloobacter | gi\|33862844\|ref\|NP_894404.1\| permease [Prochlorococcus marinus str. MIT 9313] |
| 2 | PSC clade basal to Gloobacter | gi\|33862853\|ref\|NP_894413.1\| D-Ala-D-Ala dipeptidase [Prochlorococcus marinus str. MIT 9313] |
| 2 | PSC clade basal to Gloobacter | gi\|33862854\|ref\|NP_894414.1\| ATP-dependent DNA helicase RecG [Prochlorococcus marinus str. MIT 9313] |
| 2 | PSC clade basal to Gloobacter | gi\|33862873\|ref\|NP_894433.1\| serine:pyruvate/alanine:glyoxylate aminotransferase [Prochlorococcus marinus str. MIT 9313] |
| 2 | PSC clade basal to Gloobacter | gi\|33862874\|ref\|NP_894434.1\| glutamine synthetase [Prochlorococcus marinus str. MIT 9313] |
| 2 | PSC clade basal to Gloobacter | gi\|33862890\|ref\|NP_894450.1\| preprotein translocase subunit SecD [Prochlorococcus marinus str. MIT 9313] |
| 2 | PSC clade basal to Gloobacter | gi\|33862893\|ref\|NP_894453.1\| 4-diphosphocytidyl-2-C-methyl-D-erythritol kinase [Prochlorococcus marinus str. MIT 9313] |
| 2 | PSC clade basal to Gloobacter | gi\|33862894\|ref\|NP_894454.1\| dimethyladenosine transferase [Prochlorococcus marinus str. MIT 9313] |
| 2 | PSC clade basal to Gloobacter | gi\|33862920\|ref\|NP_894480.1\| DNA polymerase III subunit alpha [Prochlorococcus marinus str. MIT 9313] |
| 2 | PSC clade basal to Gloobacter | gi\|33862927\|ref\|NP_894487.1\| anthranilate phosphoribosyltransferase [Prochlorococcus marinus str. MIT 9313] |
| 2 | PSC clade basal to Gloobacter | gi\|33862939\|ref\|NP_894499.1\| 3-dehydroquinate synthase [Prochlorococcus marinus str. MIT 9313] |
| 2 | PSC clade basal to Gloobacter | gi\|33862951\|ref\|NP_894511.1\| pyruvate kinase [Prochlorococcus marinus str. MIT 9313] |
| 2 | PSC clade basal to Gloobacter | gi\|33862970\|ref\|NP_894530.1\| ATP phosphoribosyltransferase regulatory subunit [Prochlorococcus marinus str. MIT 9313] |
| 2 | PSC clade basal to Gloobacter | gi\|33862975\|ref\|NP_894535.1\| molecular chaperone DnaK [Prochlorococcus marinus str. MIT 9313] |
| 2 | PSC clade basal to Gloobacter | gi\|33862980\|ref\|NP_894540.1\| bifunctional 3,4-dihydroxy-2-butanone 4-phosphate synthase/GTP cyclohydrolase II [Prochlorococcus marinus str. MIT 9313] |
| 2 | PSC clade basal to Gloobacter | gi\|33862988\|ref\|NP_894548.1\| diaminopimelate epimerase [Prochlorococcus marinus str. MIT 9313] |
| 2 | PSC clade basal to Gloobacter | gi\|33862989\|ref\|NP_894549.1\| class-V aminotransferase family cysteine desulfurase [Prochlorococcus marinus str. MIT 9313] |
| 2 | PSC clade basal to Gloobacter | gi\|33862999\|ref\|NP_894559.1\| cobalamin biosynthetic protein CobN [Prochlorococcus marinus str. MIT 9313] |
| 2 | PSC clade basal to Gloobacter | gi\|33863009\|ref\|NP_894569.1\| RNA methyltransferase [Prochlorococcus marinus str. MIT 9313] |
| 2 | PSC clade basal to Gloobacter | gi\|33863013\|ref\|NP_894573.1\| ribonuclease II [Prochlorococcus marinus str. MIT 9313] |
| 2 | PSC clade basal to Gloobacter | gi\|33863033\|ref\|NP_894593.1\| DHH domain-containing protein [Prochlorococcus marinus str. MIT 9313] |
| 2 | PSC clade basal to Gloobacter | gi\|33863038\|ref\|NP_894598.1\| acetazolamide conferring resistance protein Zam [Prochlorococcus marinus str. MIT 9313] |
| 2 | PSC clade basal to Gloobacter | gi\|33863039\|ref\|NP_894599.1\| aromatic acid decarboxylase [Prochlorococcus marinus str. MIT 9313] |
| 2 | PSC clade basal to Gloobacter | gi\|33863047\|ref\|NP_894607.1\| transcriptional-repair coupling factor [Prochlorococcus marinus str. MIT 9313] |
| 2 | PSC clade basal to Gloobacter | gi\|33863082\|ref\|NP_894642.1\| adenine phosphoribosyltransferase [Prochlorococcus marinus str. MIT 9313] |
| 2 | PSC clade basal to Gloobacter | gi\|33863084\|ref\|NP_894644.1\| 2-octaprenyl-6-methoxyphenol 4-monooxygenase [Prochlorococcus marinus str. MIT 9313] |
| 2 | PSC clade basal to Gloobacter | gi\|33863086\|ref\|NP_894646.1\| dihydrodipicolinate reductase [Prochlorococcus marinus str. MIT 9313] |
| 2 | PSC clade basal to Gloobacter | gi\|33863096\|ref\|NP_894656.1\| hypothetical protein PMT0824 [Prochlorococcus marinus str. MIT 9313] |
| 2 | PSC clade basal to Gloobacter | gi\|33863120\|ref\|NP_894680.1\| NADH dehydrogenase (complex I) subunit [Prochlorococcus marinus str. MIT 9313] |
| 2 | PSC clade basal to Gloobacter | gi\|33863130\|ref\|NP_894690.1\| ferric uptake regulator family protein [Prochlorococcus marinus str. MIT 9313] |
| 2 | PSC clade basal to Gloobacter | gi\|33863143\|ref\|NP_894703.1\| short-chain dehydrogenase/reductase [Prochlorococcus marinus str. MIT 9313] |
| 2 | PSC clade basal to Gloobacter | gi\|33863264\|ref\|NP_894824.1\| ABC transporter substrate-binding protein, phosphate [Prochlorococcus marinus str. MIT 9313] |
| 2 | PSC clade basal to Gloobacter | gi\|33863268\|ref\|NP_894828.1\| two component sensor histidine kinase fragment, pseudogene, partial [Prochlorococcus marinus str. MIT 9313] |
| 2 | PSC clade basal to Gloobacter | gi\|33863272\|ref\|NP_894832.1\| ArsR family regulatory protein [Prochlorococcus marinus str. MIT 9313] |
| 2 | PSC clade basal to Gloobacter | gi\|33863283\|ref\|NP_894843.1\| Alkyl hydroperoxide reductase/ Thiol specific antioxidant/ Mal allergens family protein [Prochlorococcus marinus str. MIT 9313] |
| 2 | PSC clade basal to Gloobacter | gi\|33863302\|ref\|NP_894862.1\| dienelactone hydrolase [Prochlorococcus marinus str. MIT 9313] |
| 2 | PSC clade basal to Gloobacter | gi\|33863327\|ref\|NP_894887.1\| biotin synthase [Prochlorococcus marinus str. MIT 9313] |
| 2 | PSC clade basal to Gloobacter | gi\|33863336\|ref\|NP_894896.1\| phosphatidate cytidylyltransferase [Prochlorococcus marinus str. MIT 9313] |
| 2 | PSC clade basal to Gloobacter | gi\|33863338\|ref\|NP_894898.1\| kinase [Prochlorococcus marinus str. MIT 9313] |
| 2 | PSC clade basal to Gloobacter | gi\|33863347\|ref\|NP_894907.1\| lipoyl synthase [Prochlorococcus marinus str. MIT 9313] |
| 2 | PSC clade basal to Gloobacter | gi\|33863350\|ref\|NP_894910.1\| membrane bound transcriptional regulator [Prochlorococcus marinus str. MIT 9313] |
| 2 | PSC clade basal to Gloobacter | gi\|33863355\|ref\|NP_894915.1\| sugar-phosphate nucleotidyl transferase [Prochlorococcus marinus str. MIT 9313] |
| 2 | PSC clade basal to Gloobacter | gi\|33863363\|ref\|NP_894923.1\| phospholipid/glycerol acyltransferase [Prochlorococcus marinus str. MIT 9313] |
| 2 | PSC clade basal to Gloobacter | gi\|33863379\|ref\|NP_894939.1\| hypothetical protein PMT1108 [Prochlorococcus marinus str. MIT 9313] |
| 2 | PSC clade basal to Gloobacter | gi\|33863380\|ref\|NP_894940.1\| polyprenyl synthetase [Prochlorococcus marinus str. MIT 9313] |
| 2 | PSC clade basal to Gloobacter | gi\|33863382\|ref\|NP_894942.1\| hypothetical protein PMT1111 [Prochlorococcus marinus str. MIT 9313] |
| 2 | PSC clade basal to Gloobacter | gi\|33863392\|ref\|NP_894952.1\| 2-isopropylmalate synthase [Prochlorococcus marinus str. MIT 9313] |
| 2 | PSC clade basal to Gloobacter | gi\|33863403\|ref\|NP_894963.1\| tRNA/rRNA methyltransferase [Prochlorococcus marinus str. MIT 9313] |
| 2 | PSC clade basal to Gloobacter | gi\|33863417\|ref\|NP_894977.1\| peptide ABC transporter [Prochlorococcus marinus str. MIT 9313] |
| 2 | PSC clade basal to Gloobacter | gi\|33863426\|ref\|NP_894986.1\| membrane fusion protein [Prochlorococcus marinus str. MIT 9313] |
| 2 | PSC clade basal to Gloobacter | gi\|33863427\|ref\|NP_894987.1\| DNA polymerase I [Prochlorococcus marinus str. MIT 9313] |
| 2 | PSC clade basal to Gloobacter | gi\|33863434\|ref\|NP_894994.1\| nicotinamide nucleotide transhydrogenase subunit beta [Prochlorococcus marinus str. MIT 9313] |
| 2 | PSC clade basal to Gloobacter | gi\|33863435\|ref\|NP_894995.1\| nicotinamide nucleotide transhydrogenase subunit alpha 2 (A2) [Prochlorococcus marinus str. MIT 9313] |
| 2 | PSC clade basal to Gloobacter | gi\|33863436\|ref\|NP_894996.1\| nicotinamide nucleotide transhydrogenase subunit alpha 1 (A1) [Prochlorococcus marinus str. MIT 9313] |
| 2 | PSC clade basal to Gloobacter | gi\|33863446\|ref\|NP_895006.1\| acetolactate synthase 3 regulatory subunit [Prochlorococcus marinus str. MIT 9313] |
| 2 | PSC clade basal to Gloobacter | gi\|33863474\|ref\|NP_895034.1\| ribulose bisphosphate carboxylase, small chain [Prochlorococcus marinus str. MIT 9313] |
| 2 | PSC clade basal to Gloobacter | gi\|33863483\|ref\|NP_895043.1\| sulfate transporter [Prochlorococcus marinus str. MIT 9313] |
| 2 | PSC clade basal to Gloobacter | gi\|33863492\|ref\|NP_895052.1\| protein ligase [Prochlorococcus marinus str. MIT 9313] |
| 2 | PSC clade basal to Gloobacter | gi\|33863498\|ref\|NP_895058.1\| hypothetical protein PMT1230 [Prochlorococcus marinus str. MIT 9313] |
| 2 | PSC clade basal to Gloobacter | gi\|33863515\|ref\|NP_895075.1\| short-chain dehydrogenase/reductase [Prochlorococcus marinus str. MIT 9313] |
| 2 | PSC clade basal to Gloobacter | gi\|33863529\|ref\|NP_895089.1\| adenylosuccinate synthetase [Prochlorococcus marinus str. MIT 9313] |
| 2 | PSC clade basal to Gloobacter | gi\|33863530\|ref\|NP_895090.1\| carbohydrate kinase [Prochlorococcus marinus str. MIT 9313] |
| 2 | PSC clade basal to Gloobacter | gi\|161350043\|ref\|NP_895126.2\| nucleotide-binding protein [Prochlorococcus marinus str. MIT 9313] |
| 2 | PSC clade basal to Gloobacter | gi\|33863567\|ref\|NP_895127.1\| hypothetical protein PMT1299 [Prochlorococcus marinus str. MIT 9313] |
| 2 | PSC clade basal to Gloobacter | gi\|33863577\|ref\|NP_895137.1\| CPA1 family Na+/H+ antiporter [Prochlorococcus marinus str. MIT 9313] |
| 2 | PSC clade basal to Gloobacter | gi\|33863580\|ref\|NP_895140.1\| UDP-N-acetylglucosamine 2-epimerase [Prochlorococcus marinus str. MIT 9313] |
| 2 | PSC clade basal to Gloobacter | gi\|33863587\|ref\|NP_895147.1\| Tat family protein secretion protein [Prochlorococcus marinus str. MIT 9313] |
| 2 | PSC clade basal to Gloobacter | gi\|33863596\|ref\|NP_895156.1\| hypothetical protein PMT1329 [Prochlorococcus marinus str. MIT 9313] |
| 2 | PSC clade basal to Gloobacter | gi\|33863608\|ref\|NP_895168.1\| cytochrome c oxidase, subunit 2 [Prochlorococcus marinus str. MIT 9313] |
| 2 | PSC clade basal to Gloobacter | gi\|33863609\|ref\|NP_895169.1\| cytochrome c oxidase subunit I [Prochlorococcus marinus str. MIT 9313] |
| 2 | PSC clade basal to Gloobacter | gi\|33863610\|ref\|NP_895170.1\| cytochrome c oxidase subunit III [Prochlorococcus marinus str. MIT 9313] |
| 2 | PSC clade basal to Gloobacter | gi\|33863615\|ref\|NP_895175.1\| aldo/keto reductase [Prochlorococcus marinus str. MIT 9313] |
| 2 | PSC clade basal to Gloobacter | gi\|33863621\|ref\|NP_895181.1\| NAD(P)H-quinone oxidoreductase subunit 2 [Prochlorococcus marinus str. MIT 9313] |
| 2 | PSC clade basal to Gloobacter | gi\|33863627\|ref\|NP_895187.1\| class-I aminotransferase [Prochlorococcus marinus str. MIT 9313] |
| 2 | PSC clade basal to Gloobacter | gi\|33863659\|ref\|NP_895219.1\| hypothetical protein PMT1392 [Prochlorococcus marinus str. MIT 9313] |
| 2 | PSC clade basal to Gloobacter | gi\|33863662\|ref\|NP_895222.1\| histone-like DNA-binding protein [Prochlorococcus marinus str. MIT 9313] |
| 2 | PSC clade basal to Gloobacter | gi\|33863664\|ref\|NP_895224.1\| isoamylase [Prochlorococcus marinus str. MIT 9313] |
| 2 | PSC clade basal to Gloobacter | gi\|33863665\|ref\|NP_895225.1\| GPH family sugar transporter [Prochlorococcus marinus str. MIT 9313] |
| 2 | PSC clade basal to Gloobacter | gi\|33863675\|ref\|NP_895235.1\| peptide methionine sulfoxide reductase [Prochlorococcus marinus str. MIT 9313] |
| 2 | PSC clade basal to Gloobacter | gi\|33863677\|ref\|NP_895237.1\| UDP-N-acetylglucosamine acyltransferase [Prochlorococcus marinus str. MIT 9313] |
| 2 | PSC clade basal to Gloobacter | gi\|33863679\|ref\|NP_895239.1\| UDP-3-O-[3-hydroxymyristoyl] N-acetylglucosamine deacetylase [Prochlorococcus marinus str. MIT 9313] |
| 2 | PSC clade basal to Gloobacter | gi\|33863685\|ref\|NP_895245.1\| circadian clock protein KaiC [Prochlorococcus marinus str. MIT 9313] |
| 2 | PSC clade basal to Gloobacter | gi\|33863686\|ref\|NP_895246.1\| circadian clock protein KaiB [Prochlorococcus marinus str. MIT 9313] |
| 2 | PSC clade basal to Gloobacter | gi\|33863692\|ref\|NP_895252.1\| sporulation protein SpoIID [Prochlorococcus marinus str. MIT 9313] |
| 2 | PSC clade basal to Gloobacter | gi\|33863696\|ref\|NP_895256.1\| 2Fe-2S ferredoxin [Prochlorococcus marinus str. MIT 9313] |
| 2 | PSC clade basal to Gloobacter | gi\|33863698\|ref\|NP_895258.1\| D-3-phosphoglycerate dehydrogenase [Prochlorococcus marinus str. MIT 9313] |
| 2 | PSC clade basal to Gloobacter | gi\|33863735\|ref\|NP_895295.1\| F0F1 ATP synthase subunit delta [Prochlorococcus marinus str. MIT 9313] |
| 2 | PSC clade basal to Gloobacter | gi\|33863744\|ref\|NP_895304.1\| c-type cytochrome biogenesis protein Ccs1 [Prochlorococcus marinus str. MIT 9313] |
| 2 | PSC clade basal to Gloobacter | gi\|33863748\|ref\|NP_895308.1\| nitrogen regulatory protein P-II [Prochlorococcus marinus str. MIT 9313] |
| 2 | PSC clade basal to Gloobacter | gi\|33863752\|ref\|NP_895312.1\| fumarate hydratase [Prochlorococcus marinus str. MIT 9313] |
| 2 | PSC clade basal to Gloobacter | gi\|33863754\|ref\|NP_895314.1\| DNA helicase [Prochlorococcus marinus str. MIT 9313] |
| 2 | PSC clade basal to Gloobacter | gi\|33863772\|ref\|NP_895332.1\| DNA-directed RNA polymerase subunit beta' [Prochlorococcus marinus str. MIT 9313] |
| 2 | PSC clade basal to Gloobacter | gi\|33863773\|ref\|NP_895333.1\| DNA-directed RNA polymerase subunit gamma [Prochlorococcus marinus str. MIT 9313] |
| 2 | PSC clade basal to Gloobacter | gi\|33863774\|ref\|NP_895334.1\| DNA-directed RNA polymerase subunit beta [Prochlorococcus marinus str. MIT 9313] |
| 2 | PSC clade basal to Gloobacter | gi\|33863776\|ref\|NP_895336.1\| 30S ribosomal protein S20 [Prochlorococcus marinus str. MIT 9313] |
| 2 | PSC clade basal to Gloobacter | gi\|33863777\|ref\|NP_895337.1\| histidinol dehydrogenase [Prochlorococcus marinus str. MIT 9313] |
| 2 | PSC clade basal to Gloobacter | gi\|33863793\|ref\|NP_895353.1\| transcription elongation factor NusA [Prochlorococcus marinus str. MIT 9313] |
| 2 | PSC clade basal to Gloobacter | gi\|33863824\|ref\|NP_895384.1\| oxidoreductase [Prochlorococcus marinus str. MIT 9313] |
| 2 | PSC clade basal to Gloobacter | gi\|33863832\|ref\|NP_895392.1\| cytochrome P450 enzyme [Prochlorococcus marinus str. MIT 9313] |
| 2 | PSC clade basal to Gloobacter | gi\|33863851\|ref\|NP_895411.1\| SMC ATPase superfamily chromosome segregation protein [Prochlorococcus marinus str. MIT 9313] |
| 2 | PSC clade basal to Gloobacter | gi\|33863853\|ref\|NP_895413.1\| methionine sulfoxide reductase family protein [Prochlorococcus marinus str. MIT 9313] |
| 2 | PSC clade basal to Gloobacter | gi\|33863854\|ref\|NP_895414.1\| hypothetical protein PMT1587 [Prochlorococcus marinus str. MIT 9313] |
| 2 | PSC clade basal to Gloobacter | gi\|33863868\|ref\|NP_895428.1\| peptide deformylase [Prochlorococcus marinus str. MIT 9313] |
| 2 | PSC clade basal to Gloobacter | gi\|33863874\|ref\|NP_895434.1\| AsnC family regulatory protein [Prochlorococcus marinus str. MIT 9313] |
| 2 | PSC clade basal to Gloobacter | gi\|33863889\|ref\|NP_895449.1\| bacterioferritin comigratory (BCP) protein [Prochlorococcus marinus str. MIT 9313] |
| 2 | PSC clade basal to Gloobacter | gi\|33863892\|ref\|NP_895452.1\| NADH dehydrogenase, transport associated [Prochlorococcus marinus str. MIT 9313] |
| 2 | PSC clade basal to Gloobacter | gi\|33863906\|ref\|NP_895466.1\| DEAD/DEAH box helicase [Prochlorococcus marinus str. MIT 9313] |
| 2 | PSC clade basal to Gloobacter | gi\|33863928\|ref\|NP_895488.1\| SMF family protein [Prochlorococcus marinus str. MIT 9313] |
| 2 | PSC clade basal to Gloobacter | gi\|33863937\|ref\|NP_895497.1\| S-adenosylmethionine synthetase [Prochlorococcus marinus str. MIT 9313] |
| 2 | PSC clade basal to Gloobacter | gi\|33863970\|ref\|NP_895530.1\| ribonuclease D [Prochlorococcus marinus str. MIT 9313] |
| 2 | PSC clade basal to Gloobacter | gi\|33863974\|ref\|NP_895534.1\| ATPase [Prochlorococcus marinus str. MIT 9313] |
| 2 | PSC clade basal to Gloobacter | gi\|33863975\|ref\|NP_895535.1\| hypothetical protein PMT1708 [Prochlorococcus marinus str. MIT 9313] |
| 2 | PSC clade basal to Gloobacter | gi\|33863978\|ref\|NP_895538.1\| anthranilate synthase component I/chorismate-binding protein [Prochlorococcus marinus str. MIT 9313] |
| 2 | PSC clade basal to Gloobacter | gi\|33863986\|ref\|NP_895546.1\| molybdopterin biosynthesis protein [Prochlorococcus marinus str. MIT 9313] |
| 2 | PSC clade basal to Gloobacter | gi\|33864014\|ref\|NP_895574.1\| 50S ribosomal protein L18 [Prochlorococcus marinus str. MIT 9313] |
| 2 | PSC clade basal to Gloobacter | gi\|33864021\|ref\|NP_895581.1\| 30S ribosomal protein S11 [Prochlorococcus marinus str. MIT 9313] |
| 2 | PSC clade basal to Gloobacter | gi\|33864022\|ref\|NP_895582.1\| DNA-directed RNA polymerase subunit alpha [Prochlorococcus marinus str. MIT 9313] |
| 2 | PSC clade basal to Gloobacter | gi\|33864024\|ref\|NP_895584.1\| tRNA pseudouridine synthase A [Prochlorococcus marinus str. MIT 9313] |
| 2 | PSC clade basal to Gloobacter | gi\|33864030\|ref\|NP_895590.1\| alanine racemase [Prochlorococcus marinus str. MIT 9313] |
| 2 | PSC clade basal to Gloobacter | gi\|33864039\|ref\|NP_895599.1\| bifunctional cbiH protein and precorrin-3B C17-methyltransferase [Prochlorococcus marinus str. MIT 9313] |
| 2 | PSC clade basal to Gloobacter | gi\|33864042\|ref\|NP_895602.1\| lipoyl synthase [Prochlorococcus marinus str. MIT 9313] |
| 2 | PSC clade basal to Gloobacter | gi\|33864057\|ref\|NP_895617.1\| hypothetical protein PMT1790 [Prochlorococcus marinus str. MIT 9313] |
| 2 | PSC clade basal to Gloobacter | gi\|33864074\|ref\|NP_895634.1\| aspartate carbamoyltransferase catalytic subunit [Prochlorococcus marinus str. MIT 9313] |
| 2 | PSC clade basal to Gloobacter | gi\|33864081\|ref\|NP_895641.1\| creatininase [Prochlorococcus marinus str. MIT 9313] |
| 2 | PSC clade basal to Gloobacter | gi\|33864082\|ref\|NP_895642.1\| aspartyl/glutamyl-tRNA amidotransferase subunit C [Prochlorococcus marinus str. MIT 9313] |
| 2 | PSC clade basal to Gloobacter | gi\|33864083\|ref\|NP_895643.1\| beta carotene hydroxylase [Prochlorococcus marinus str. MIT 9313] |
| 2 | PSC clade basal to Gloobacter | gi\|33864084\|ref\|NP_895644.1\| isoleucyl-tRNA synthetase [Prochlorococcus marinus str. MIT 9313] |
| 2 | PSC clade basal to Gloobacter | gi\|33864087\|ref\|NP_895647.1\| tRNA (guanine-N(7)-)-methyltransferase [Prochlorococcus marinus str. MIT 9313] |
| 2 | PSC clade basal to Gloobacter | gi\|33864091\|ref\|NP_895651.1\| soluble lytic transglycosylase [Prochlorococcus marinus str. MIT 9313] |
| 2 | PSC clade basal to Gloobacter | gi\|33864098\|ref\|NP_895658.1\| CRP family global nitrogen regulatory protein [Prochlorococcus marinus str. MIT 9313] |
| 2 | PSC clade basal to Gloobacter | gi\|33864106\|ref\|NP_895666.1\| glycoside hydrolase family protein [Prochlorococcus marinus str. MIT 9313] |
| 2 | PSC clade basal to Gloobacter | gi\|33864119\|ref\|NP_895679.1\| sugar fermentation stimulation protein A [Prochlorococcus marinus str. MIT 9313] |
| 2 | PSC clade basal to Gloobacter | gi\|33864125\|ref\|NP_895685.1\| esterase [Prochlorococcus marinus str. MIT 9313] |
| 2 | PSC clade basal to Gloobacter | gi\|33864131\|ref\|NP_895691.1\| glycosyl transferase family protein [Prochlorococcus marinus str. MIT 9313] |
| 2 | PSC clade basal to Gloobacter | gi\|33864133\|ref\|NP_895693.1\| hypothetical protein PMT1866 [Prochlorococcus marinus str. MIT 9313] |
| 2 | PSC clade basal to Gloobacter | gi\|33864135\|ref\|NP_895695.1\| orotate phosphoribosyltransferase [Prochlorococcus marinus str. MIT 9313] |
| 2 | PSC clade basal to Gloobacter | gi\|33864138\|ref\|NP_895698.1\| phosphotransferase superclass [Prochlorococcus marinus str. MIT 9313] |
| 2 | PSC clade basal to Gloobacter | gi\|33864157\|ref\|NP_895717.1\| NADH dehydrogenase subunit J [Prochlorococcus marinus str. MIT 9313] |
| 2 | PSC clade basal to Gloobacter | gi\|161350040\|ref\|NP_895731.2\| histidyl-tRNA synthetase [Prochlorococcus marinus str. MIT 9313] |
| 2 | PSC clade basal to Gloobacter | gi\|33864175\|ref\|NP_895735.1\| selenide,water dikinase [Prochlorococcus marinus str. MIT 9313] |
| 2 | PSC clade basal to Gloobacter | gi\|33864176\|ref\|NP_895736.1\| tRNA nucleotidyltransferase/poly(A) polymerase [Prochlorococcus marinus str. MIT 9313] |
| 2 | PSC clade basal to Gloobacter | gi\|33864183\|ref\|NP_895743.1\| UvrD/REP helicase [Prochlorococcus marinus str. MIT 9313] |
| 2 | PSC clade basal to Gloobacter | gi\|33864202\|ref\|NP_895762.1\| Fe-S oxidoreductase [Prochlorococcus marinus str. MIT 9313] |
| 2 | PSC clade basal to Gloobacter | gi\|33864208\|ref\|NP_895768.1\| alpha/beta fold family hydrolase [Prochlorococcus marinus str. MIT 9313] |
| 2 | PSC clade basal to Gloobacter | gi\|33864214\|ref\|NP_895774.1\| ribonuclease III [Prochlorococcus marinus str. MIT 9313] |
| 2 | PSC clade basal to Gloobacter | gi\|33864216\|ref\|NP_895776.1\| 16S rRNA-processing protein RimM [Prochlorococcus marinus str. MIT 9313] |
| 2 | PSC clade basal to Gloobacter | gi\|33864217\|ref\|NP_895777.1\| GDP-mannose pyrophosphorylase [Prochlorococcus marinus str. MIT 9313] |
| 2 | PSC clade basal to Gloobacter | gi\|33864233\|ref\|NP_895793.1\| zeta-carotene desaturase [Prochlorococcus marinus str. MIT 9313] |
| 2 | PSC clade basal to Gloobacter | gi\|33864243\|ref\|NP_895803.1\| O-acetylserine (thiol)-lyase A [Prochlorococcus marinus str. MIT 9313] |
| 2 | PSC clade basal to Gloobacter | gi\|33864252\|ref\|NP_895812.1\| ABC transporter ATP-binding protein [Prochlorococcus marinus str. MIT 9313] |
| 2 | PSC clade basal to Gloobacter | gi\|33864254\|ref\|NP_895814.1\| DNA polymerase III subunit delta' [Prochlorococcus marinus str. MIT 9313] |
| 2 | PSC clade basal to Gloobacter | gi\|33864262\|ref\|NP_895822.1\| malonyl CoA-ACP transacylase [Prochlorococcus marinus str. MIT 9313] |
| 2 | PSC clade basal to Gloobacter | gi\|33864263\|ref\|NP_895823.1\| 1-acyl-sn-glycerol-3-phosphate acyltransferase [Prochlorococcus marinus str. MIT 9313] |
| 2 | PSC clade basal to Gloobacter | gi\|33864269\|ref\|NP_895829.1\| phytoene desaturase [Prochlorococcus marinus str. MIT 9313] |
| 2 | PSC clade basal to Gloobacter | gi\|33864279\|ref\|NP_895839.1\| LuxR family regulatory protein [Prochlorococcus marinus str. MIT 9313] |
| 2 | PSC clade basal to Gloobacter | gi\|33864281\|ref\|NP_895841.1\| inorganic polyphosphate/ATP-NAD kinase [Prochlorococcus marinus str. MIT 9313] |
| 2 | PSC clade basal to Gloobacter | gi\|33864282\|ref\|NP_895842.1\| NADH dehydrogenase subunit K [Prochlorococcus marinus str. MIT 9313] |
| 2 | PSC clade basal to Gloobacter | gi\|33864283\|ref\|NP_895843.1\| NADH dehydrogenase subunit J [Prochlorococcus marinus str. MIT 9313] |
| 2 | PSC clade basal to Gloobacter | gi\|33864284\|ref\|NP_895844.1\| NADH dehydrogenase subunit I [Prochlorococcus marinus str. MIT 9313] |
| 2 | PSC clade basal to Gloobacter | gi\|33864294\|ref\|NP_895854.1\| adenylylsulfate kinase [Prochlorococcus marinus str. MIT 9313] |
| 2 | PSC clade basal to Gloobacter | gi\|33864307\|ref\|NP_895867.1\| pseudouridine synthase [Prochlorococcus marinus str. MIT 9313] |
| 2 | PSC clade basal to Gloobacter | gi\|33864322\|ref\|NP_895882.1\| O-succinylbenzoic acid--CoA ligase [Prochlorococcus marinus str. MIT 9313] |
| 2 | PSC clade basal to Gloobacter | gi\|33864330\|ref\|NP_895890.1\| hypothetical protein PMT2065 [Prochlorococcus marinus str. MIT 9313] |
| 2 | PSC clade basal to Gloobacter | gi\|33864330\|ref\|NP_895890.1\| hypothetical protein PMT2065 [Prochlorococcus marinus str. MIT 9313] |
| 2 | PSC clade basal to Gloobacter | gi\|33864332\|ref\|NP_895892.1\| para-aminobenzoate synthase component II [Prochlorococcus marinus str. MIT 9313] |
| 2 | PSC clade basal to Gloobacter | gi\|33864334\|ref\|NP_895894.1\| class-I aminotransferase [Prochlorococcus marinus str. MIT 9313] |
| 2 | PSC clade basal to Gloobacter | gi\|33864346\|ref\|NP_895906.1\| kinase [Prochlorococcus marinus str. MIT 9313] |
| 2 | PSC clade basal to Gloobacter | gi\|33864347\|ref\|NP_895907.1\| phosphopyruvate hydratase [Prochlorococcus marinus str. MIT 9313] |
| 2 | PSC clade basal to Gloobacter | gi\|33864355\|ref\|NP_895915.1\| 50S ribosomal protein L7/L12 [Prochlorococcus marinus str. MIT 9313] |
| 2 | PSC clade basal to Gloobacter | gi\|33864357\|ref\|NP_895917.1\| ribonuclease HI [Prochlorococcus marinus str. MIT 9313] |
| 2 | PSC clade basal to Gloobacter | gi\|33864362\|ref\|NP_895922.1\| general (type II) secretion pathway protein D precursor [Prochlorococcus marinus str. MIT 9313] |
| 2 | PSC clade basal to Gloobacter | gi\|33864373\|ref\|NP_895933.1\| pseudouridylate synthase specific to ribosomal large subunit [Prochlorococcus marinus str. MIT 9313] |
| 2 | PSC clade basal to Gloobacter | gi\|33864375\|ref\|NP_895935.1\| guanosine-3',5'-bis(diphosphate) 3'-diphosphatase [Prochlorococcus marinus str. MIT 9313] |
| 2 | PSC clade basal to Gloobacter | gi\|33864415\|ref\|NP_895975.1\| GntR family transcriptional regulator [Prochlorococcus marinus str. MIT 9313] |
| 2 | PSC clade basal to Gloobacter | gi\|33864422\|ref\|NP_895982.1\| SNF2/helicase domain-containing protein [Prochlorococcus marinus str. MIT 9313] |
| 2 | PSC clade basal to Gloobacter | gi\|33864423\|ref\|NP_895983.1\| hypothetical protein PMT2159 [Prochlorococcus marinus str. MIT 9313] |
| 2 | PSC clade basal to Gloobacter | gi\|33864425\|ref\|NP_895985.1\| TRAP-T family tripartite transporter substrate-binding protein [Prochlorococcus marinus str. MIT 9313] |
| 2 | PSC clade basal to Gloobacter | gi\|33864427\|ref\|NP_895987.1\| TRAP-T family tripartite transporter [Prochlorococcus marinus str. MIT 9313] |
| 2 | PSC clade basal to Gloobacter | gi\|33864428\|ref\|NP_895988.1\| metallo-beta-lactamase domain-containing protein [Prochlorococcus marinus str. MIT 9313] |
| 2 | PSC clade basal to Gloobacter | gi\|33864429\|ref\|NP_895989.1\| flavodoxin:flavin reductase-like domain-containing protein [Prochlorococcus marinus str. MIT 9313] |
| 2 | PSC clade basal to Gloobacter | gi\|33864440\|ref\|NP_896000.1\| DnaB replicative helicase [Prochlorococcus marinus str. MIT 9313] |
| 2 | PSC clade basal to Gloobacter | gi\|33864441\|ref\|NP_896001.1\| tRNA uridine 5-carboxymethylaminomethyl modification protein GidA [Prochlorococcus marinus str. MIT 9313] |
| 2 | PSC clade basal to Gloobacter | gi\|33864448\|ref\|NP_896008.1\| NAD-dependent DNA ligase LigA [Prochlorococcus marinus str. MIT 9313] |
| 2 | PSC clade basal to Gloobacter | gi\|33864463\|ref\|NP_896023.1\| pyridoxamine 5'-phosphate oxidase [Prochlorococcus marinus str. MIT 9313] |
| 2 | PSC clade basal to Gloobacter | gi\|33864467\|ref\|NP_896027.1\| ABC transporter substrate-binding protein [Prochlorococcus marinus str. MIT 9313] |
| 2 | PSC clade basal to Gloobacter | gi\|33864471\|ref\|NP_896031.1\| permease [Prochlorococcus marinus str. MIT 9313] |
| 2 | PSC clade basal to Gloobacter | gi\|161350039\|ref\|NP_896037.2\| spermidine synthase [Prochlorococcus marinus str. MIT 9313] |
| 2 | PSC clade basal to Gloobacter | gi\|33864478\|ref\|NP_896038.1\| arginase [Prochlorococcus marinus str. MIT 9313] |
| 2 | PSC clade basal to Gloobacter | gi\|33864487\|ref\|NP_896047.1\| p-aminobenzoate synthetase [Prochlorococcus marinus str. MIT 9313] |
| 2 | PSC clade basal to Gloobacter | gi\|33864491\|ref\|NP_896051.1\| ABC transporter membrane protein [Prochlorococcus marinus str. MIT 9313] |
| 2 | PSC clade basal to Gloobacter | gi\|33864493\|ref\|NP_896053.1\| urea ABC transporter substrate-binding protein [Prochlorococcus marinus str. MIT 9313] |
| 2 | PSC clade basal to Gloobacter | gi\|33864494\|ref\|NP_896054.1\| urease accessory protein UreG [Prochlorococcus marinus str. MIT 9313] |
| 2 | PSC clade basal to Gloobacter | gi\|33864495\|ref\|NP_896055.1\| urease accessory protein ureF [Prochlorococcus marinus str. MIT 9313] |
| 2 | PSC clade basal to Gloobacter | gi\|33864500\|ref\|NP_896060.1\| urease subunit alpha [Prochlorococcus marinus str. MIT 9313] |
| 2 | PSC clade basal to Gloobacter | gi\|33864501\|ref\|NP_896061.1\| uroporphyrin-III C-methyltransferase [Prochlorococcus marinus str. MIT 9313] |
| 2 | PSC clade basal to Gloobacter | gi\|33864506\|ref\|NP_896066.1\| hypothetical protein PMT2242 [Prochlorococcus marinus str. MIT 9313] |
| 2 | PSC clade basal to Gloobacter | gi\|33864508\|ref\|NP_896068.1\| major facilitator superfamily multidrug-efflux transporter [Prochlorococcus marinus str. MIT 9313] |
| 2 | PSC clade basal to Gloobacter | gi\|33864514\|ref\|NP_896074.1\| chloride channel [Prochlorococcus marinus str. MIT 9313] |
| 2 | PSC clade basal to Gloobacter | gi\|33864515\|ref\|NP_896075.1\| hypothetical protein PMT2251 [Prochlorococcus marinus str. MIT 9313] |
| 2 | PSC clade basal to Gloobacter | gi\|33864521\|ref\|NP_896081.1\| shikimate 5-dehydrogenase [Prochlorococcus marinus str. MIT 9313] |
| 2 | PSC clade basal to Gloobacter | gi\|33864523\|ref\|NP_896083.1\| 30S ribosomal protein S6 [Prochlorococcus marinus str. MIT 9313] |
| 2 | PSC clade basal to Gloobacter | gi\|33864535\|ref\|NP_896095.1\| DNA repair protein RecN [Prochlorococcus marinus str. MIT 9313] |
