## Supplementary figures and images for "Whole genome phylogeny of Cyanobacteria documents a distinct evolutionary trajectory of marine picocyanobacteria"

### supplementary figure s1.tif

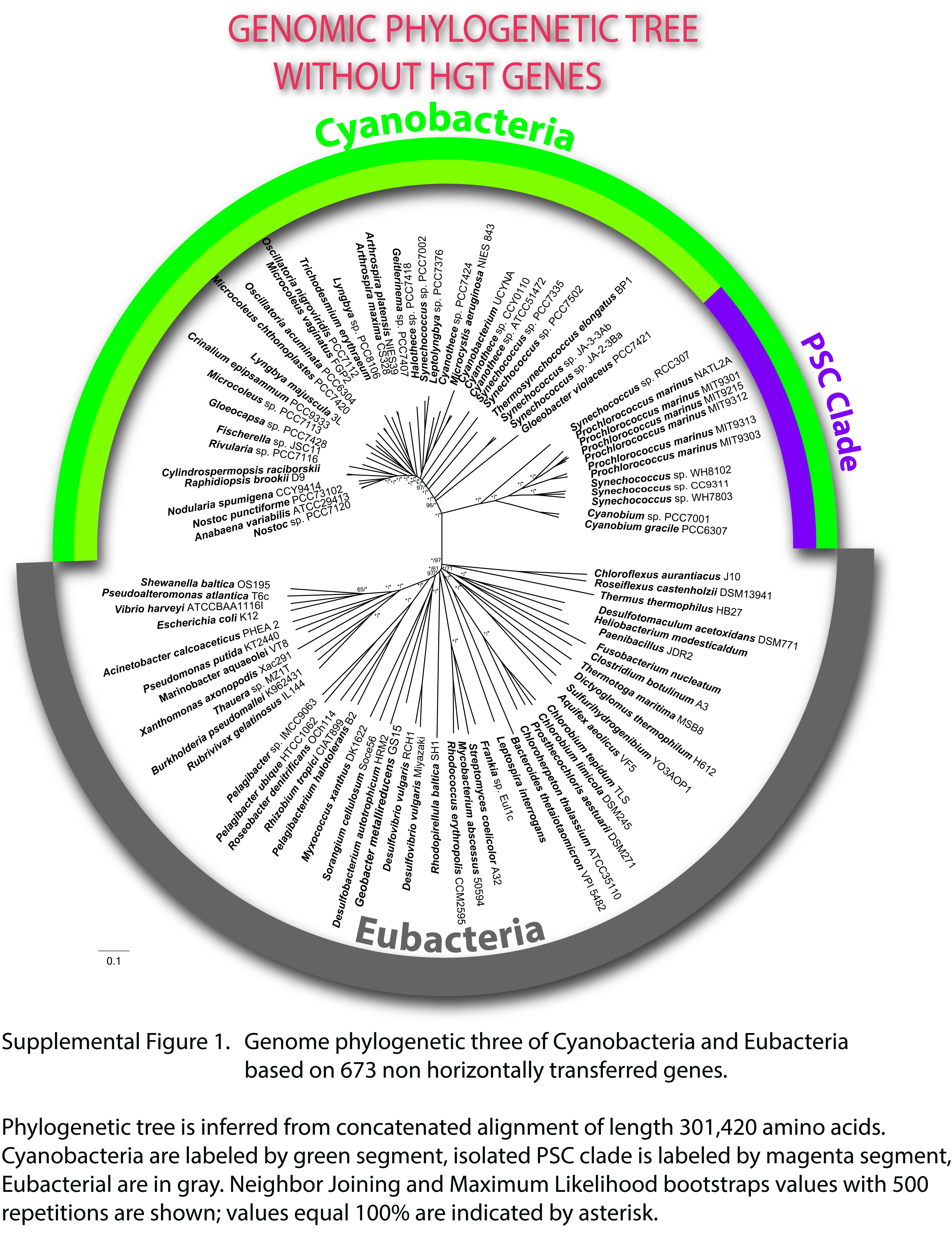

### Supplementary Figure S2.tif

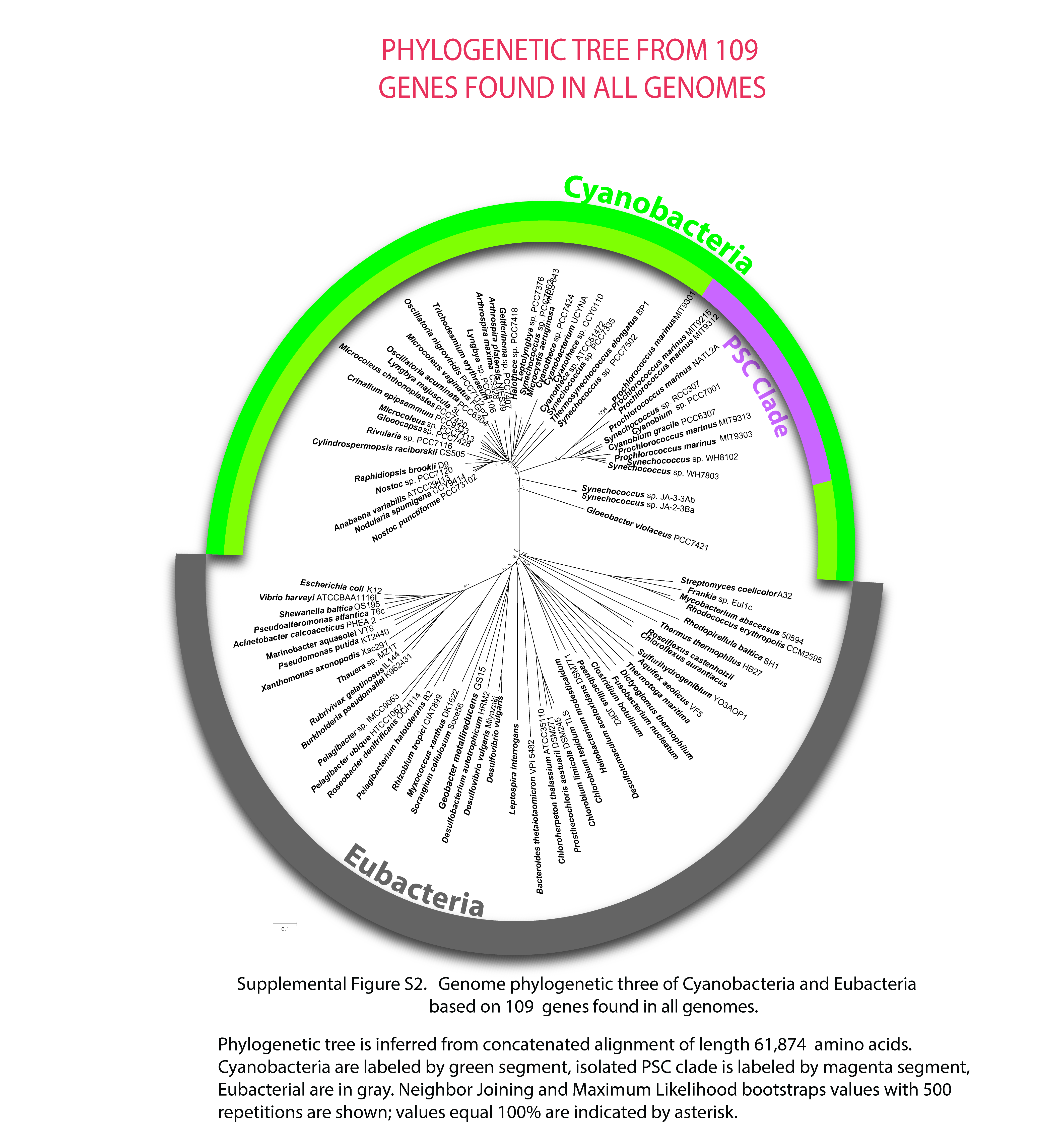

### Supplementary Figure S3.tif

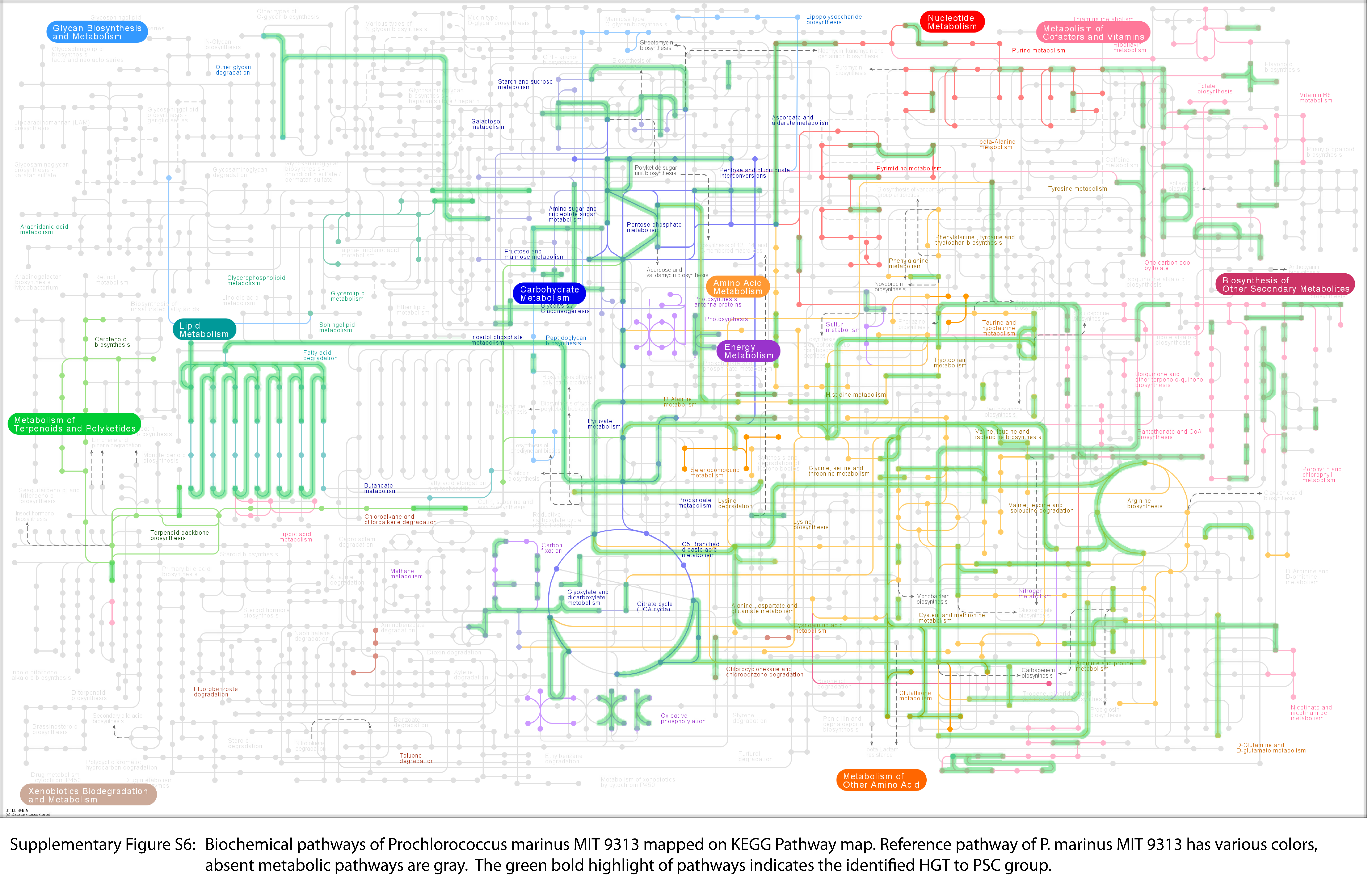
